# Chemically responsive protein switches for the precise control of biological activities

**DOI:** 10.1101/2025.05.16.654305

**Authors:** Jaime Franco Pinto, Naama Drahy, Séverine Divoux, Franck Perez, Arnaud Gautier

**Affiliations:** Sorbonne Université, École Normale Supérieure, Université PSL, CNRS, Chimie Physique et Chimie du Vivant (CPCV), Paris, France; Institut Curie, PSL University, Sorbonne Université, CNRS UMR144, Cell Biology and Cancer, 75005 Paris, France; Institut Universitaire de France, Paris, France

## Abstract

Controlling the proximity or interaction of proteins with small molecules enables researchers to chemically regulate cellular functions. Here, we leveraged CATCHFIRE (chemically assisted tethering of chimera by fluorogenic induced recognition) – a technology enabling to chemically induce dimerization in a reversible manner – to create chemically responsive proteins switches for the precise and reversible control of various biological activities. CATCHFIRE allowed us to chemically induce the assembly and thus function of various split enzymes – including luciferases, proteases, DNA recombinases. We extended this approach to develop CATCH-ON, a chemically inducible gene expression system relying on the chemically induced dimerization of the DNA-binding domain GAL4 and the truncated transcription factor p65Δ. CATCH-ON allowed us to precisely regulate the expression of cellular enzymes such as proteases, DNA recombinases, or suicide switches, as well as to control the secretion of therapeutically relevant proteins such as insulin. We showed that the CATCH-ON system is fast-acting, reversible, titratable, non-toxic and compatible with other chemically induced dimerization systems, opening exciting possibilities for its application in basic research, biotechnology and cell therapy.

## INTRODUCTION

Protein-protein interactions are key molecular recognition events controlling biological processes in living systems. Chemically induced proximity/dimerization (CIP/D) tools relying on small molecules to induce the interaction of two proteins enable to study the functional role of protein proximity in specific biological processes and to create chemically responsive systems for the precise control of biological activities. The use of non-toxic small-molecules as specific triggers present various advantages. Their ease of administration, good biodistribution and cell permeability allow simple and rapid control of biological processes in a dose-dependent and in some cases in a reversible manner. This approach has been widely used to regulate cellular processes, such as gene transcription,^1–4^ signal transduction,^5–7^ proteasome-based^8–10^ and lysosome-based^11^ protein degradation, protein post-translation modifications^12,13^ or protein folding and localization.^14–16^ Beyond basic research, chemically responsive protein switches are increasingly used to tune the function of therapeutic cells, such as CAR-T cells,^6,7,17–20^ in particular to mitigate potentially life-threatening side effects.^21–23^

A wide range of CID/P tools are now available.^21–23^ Among the most widely used are those based on rapamycin and its analogs (rapalogs), which promote the interaction of FK506-binding protein (FKBP) and FKBP-binding protein (FRB).^20,24–26^ Other CID/P systems leverage plant hormones such as abscisic acid^26^or gibberellin,^27^ which function as molecular glues to induce protein-protein interactions. Recently, CID/P tools have been designed by splitting proteins into two complementary fragments that can reassemble in presence of their cognate ligand.^4,15,28,29^ Using this approach, we introduced CATCHFIRE (chemically assisted tethering of chimera by fluorogenic-induced recognition),^15^ a unique CID system displaying fluorescence read-out. CATCHFIRE was designed by splitting pFAST,^30^ a chemogenetic fluorescent reporter made of a 14 kDa protein domain that binds and stabilizes the fluorescent state of fluorogenic ligands. The two complementary fragments of pFAST, dubbed ^FIRE^mate (114 aa, 12 kDa) and ^FIRE^tag (11 aa, 1.4 kDa), only assemble in the presence of the fluorogenic ligands. These small ligands, called match, act thus as fluorogenic molecular glues that light up in the process, enabling to monitor the dimerization process in real-time by fluorescence microscopy. The match molecules display high cell permeability, enabling rapid and reversible control of protein dimerization. This approach allowed for the precise temporal control of protein localization, organelle positioning and protein secretion. Furthermore, its intrinsic fluorogenic nature enabled the design of sensors of cellular processes^15^ CATCHFIRE offers several distinct advantages over other CIP systems, including its versatility, rapid and reversible activation, low toxicity, and seamless compatibility with existing molecular technologies. Key differentiators include (i) its fluorogenic nature, which allows self-reporting of the dimerization process, (ii) the compact size of the dimerizing domains, which facilitates easier genetic engineering and minimizes steric hindrance, as well as (iii) the rapid kinetics of dimerization and reversibility, allowing for precise temporal control. Importantly, CATCHFIRE utilizes a fully synthetic dimerizer, which a priori displays no off-target biological activity, unlike drug-based or hormone-based dimerizers such as that may elicit unintended cellular responses.^4^ A wide array of match molecules is available, each with customizable properties such as fluorescence emission, affinity, and cell permeability. These features position CATCHFIRE as a highly advantageous tool for regulating cellular processes.

Here, we present the use of CATCHFIRE for the design of chemically responsive protein switches for the precise control of protein activities and cellular processes. CATCHFIRE enabled us to chemically modulate the activity of split versions of enzymes such as luciferases, proteases and DNA recombinases, and manipulate the assembly of chimeric transcription regulators for the accurate and reversible regulation of gene expression. These chemically responsive systems enabled us to precisely regulate various cellular processes relevant for cell biology, synthetic biology or for more applied applications in biotechnology and cell therapy.

## RESULTS

### Chemically responsive luciferase

To evaluate CATCHFIRE for the control of protein function, we first engineered chemically responsive bioluminescent enzymes. Split luciferases are ideal systems as their activity can be followed by measuring their bioluminescence. We fused ^FIRE^mate and ^FIRE^tag to the SmBiT and LgBiT fragments of the NanoBiT technology derived from the blue-emitting luciferase NanoLuc^31,32^ (**Figure 1**), and tested the responsiveness of the different topologies upon match addition (**Figure 1a,b**). A preliminary screening using a two-plasmid transfection strategy identified LgBiT-^FIRE^tag and ^FIRE^mate-SmBiT as the best topology for recapitulating the NanoBiT technology (**Figure 1b**). To alleviate the mosaic effect and better control the ratios of the two protein parts, we co-expressed the two proteins using a polycistronic expression cassette containing a viral P2A sequence for ribosomal skipping during translation. We observed a 15-fold increase in bioluminescence upon match addition, demonstrating efficient bioluminescence complementation (**Figure 1c**). These performances mirror those of a rapamycin-responsive system used as a benchmark, highlighting the efficiency of CATCHFIRE in chemically reconstituting the function of NanoBiT.

**Figure 1.**
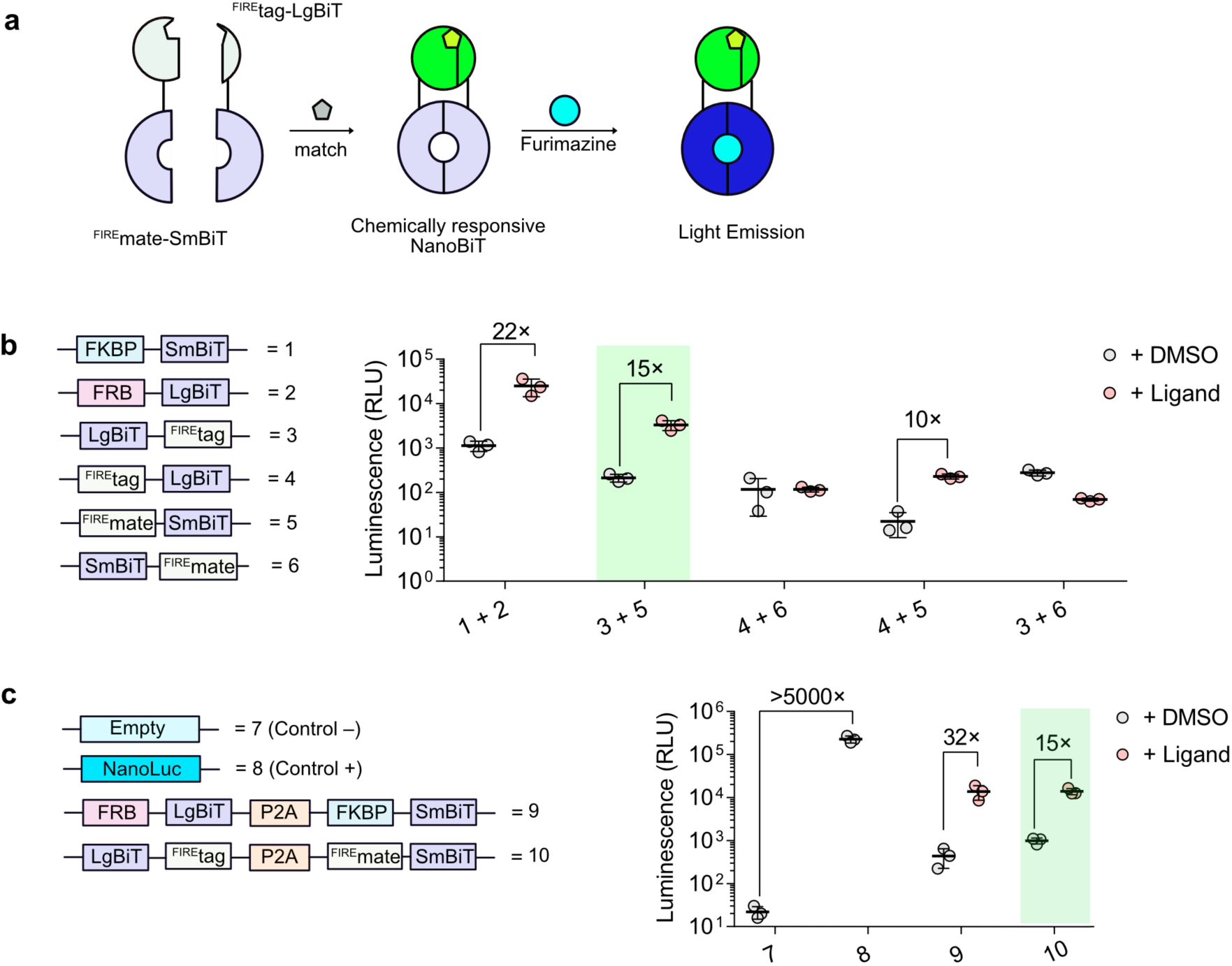
Chemically responsive luciferase. **a** Principle. **b** Initial screening and selection of best pair. Luminescence of HeLa cells expressing the chemically responsive luciferase candidates incubated for 2 h either with DMSO 0.5% or their cognate inducer (0.5 μM of rapamycin or 5 μM of match_550_). Luminescence values are shown as mean ± SD of n = 3 independent experiments (each composed of two technical replicates). **c** Luminescence of HeLa cells expressing the chemically responsive luciferase candidates incubated for 2 h either with DMSO 0.5% or their cognate inducer (0.5 μM of rapamycin or 5 μM of match_550_). Luminescence values are shown as mean ± SD of n = 3 independent experiments (each composed of two technical replicates).

### Chemically responsive protease

Next, we used CATCHFIRE to design chemically responsive split proteases. Such systems can enable precise regulation of proteins through specific cleavage.^1,5,28,29,33–36^ Specific proteolysis can change the structure and/or the localization of a protein, leading to a gain or loss of function and downstream cellular effects.^5,28,33^ One of the most popular split protease systems is derived from the tobacco etch virus protease (TEVp).^5^ Its activity was previously controlled with small molecules^5,28,29,34,35^ or light.^33^ Upon activation, TEVp cleaves a specific ENLYFQ/S sequence allowing, for example, the control of gene expression,^5,29,34^ protein secretion,^1,35^ cellular viability^36^ or the creation of logic gates for cellular control.^28,36^ We designed controllable split TEVp by fusing ^FIRE^tag and ^FIRE^mate at the N-terminus of the two split fragments of TEVp, and analyzed their match-responsiveness using a TEVp-activatable Luc2 luciferase. After 1 h of treatment with match, we observed a 5-fold increase in Luc2 bioluminescence for ^FIRE^tag-NTEVp/^FIRE^mate-CTEVp and for ^FIRE^mate-NTEVp/^FIRE^tag-CTEVp. Compared to the previously established abscisic acid (ABA)-responsive split TEVp – engineered using the ABA-dependent interaction of PYL1 and ABI^28^ – our systems displayed 2.5-fold higher dynamic range and lower background signal (**Figure 2a-c**).

**Figure 2.**
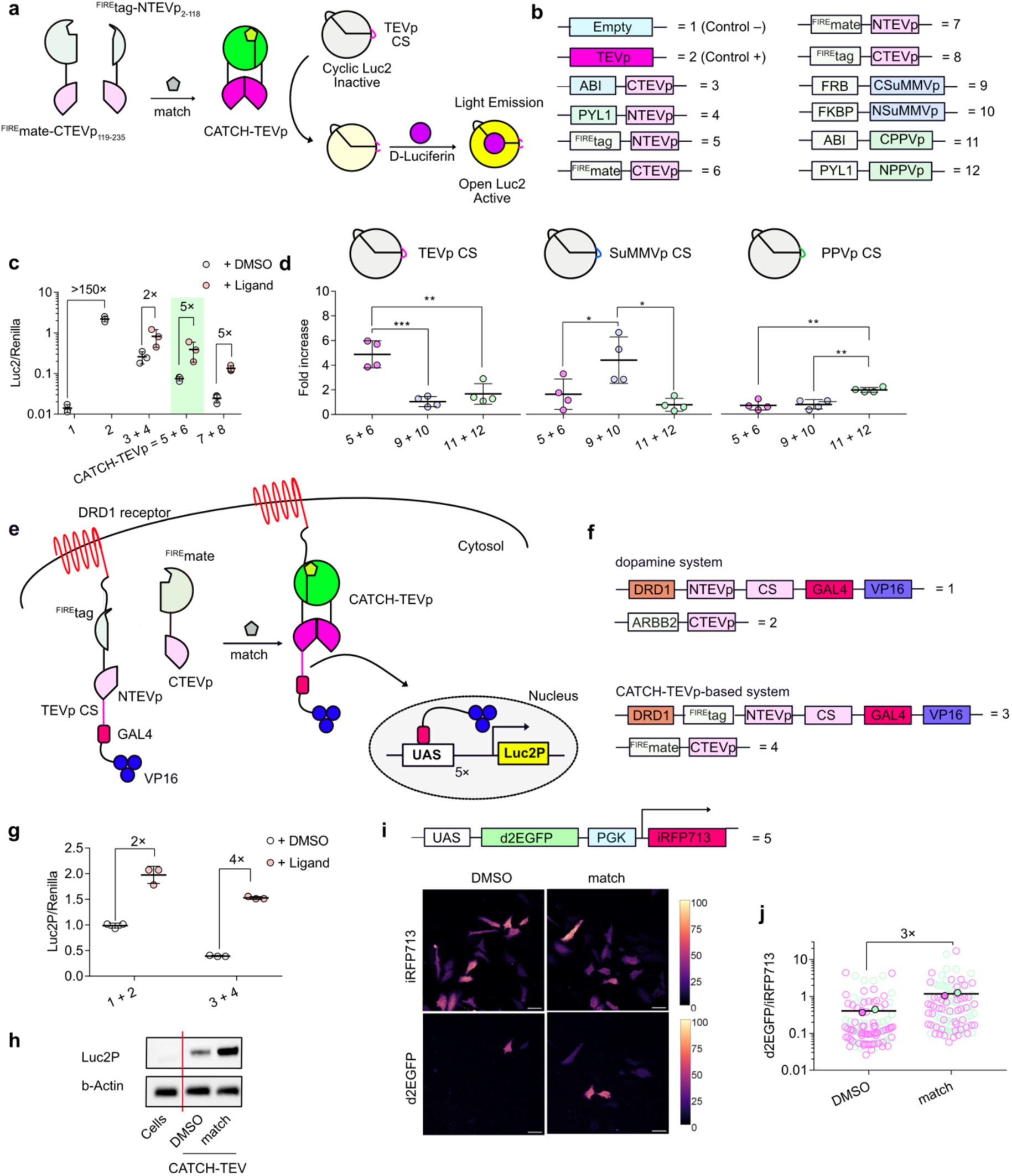
Chemically responsive protease. **a** Principle of CATCH-TEVp and the bioluminescent assay based on circularized Luc2. **b** Constructs used in this study (CS, cleavage site). **c** Bioluminescence intensity of HeLa cells co-expressing the CATCH-TEVp candidates together with the bioluminescent TEVp sensor and incubated for 1 h with 0.5% DMSO or 5 μM of match_550_. Luc2 signals were normalized by the signal of a co-expressed Renilla luciferase to overcome differences in expression levels. Control experiments using full-length TEVp or the ABA-inducible TEVp are shown (100 μM of abscisic acid (ABA) were used). Values are shown as mean ± SD of n = 3 independent experiments (each composed of two technical replicates). **d** Orthogonality of CATCH-TEVp, ABA-inducible PPVp and Rapamycin-inducible SuMMVp. The different chemically responsive proteases were expressed either with a bioluminescent TEVp sensor (left), a bioluminescent SUMMVp sensor (middle) or a bioluminescent PPVp sensor (right). Bioluminescent fold increases upon addition of the inducer are shown. ONE-way ANOVA analysis followed by a Tukey’s multiple comparisons test was performed where *p*-values are: 0.0003 and 0.0010 for TEVp, 0.0103 and 0.0406 for SUMMVp, 0.0015 and 0.0010 for PPVp. **e** Principle of CATCH-TEVp-based gene expression control. **f** Constructs used in this study (CS, cleavage site). **g** Expression fold increase upon activation of the CATCH-TEVp-based system or the dopamine-sensitive system. HeLa cells expressing the different components were treated for 24 h either with 0.5% DMSO or with the cognate inducer (5 μM of dopamine or 5 μM of match_550_). Values are shown as mean ± SD of n = 3 independent experiments (each comprised of two technical replicates). **h** Western blotting showing Luc2P expression after 24 h of incubation with DMSO 0.5% or 5 µM of match_550_. The experiment shows one representative blot out of two experiments. **i,j** Evaluation of the induced expression of d2EGFP upon activation of CATCH-TEVp by confocal microscopy experiments after 16 h of incubation with DMSO 0.5% or 5 µM of match_550_. **i** Representative confocal micrographs showing d2EGFP expression. iRFP713 was used as transfection control. Images were taken with identical microscope settings. Scale bars represent 50 µm. **j** d2EGFP fluorescence intensity (normalized by iRFP713 signal) of single cells. Each cell is color-coded according to the biological replicate it came from. The solid circles correspond to the mean of each biological replicate. The black line represents the mean values of the two biological replicates (each with n ≥ 45 cells).

An interesting feature of proteases is their composability with other modular systems to create complex networks thanks to their unique cleavage sequences.^28,36^ To test this hypothesis, we challenged our match-responsive split TEVp (hereafter called CATCH-TEVp) with other chemically responsive split proteases of the polyviral family derived from the Sunflower Mild Mosaic Virus protease (SUMMVp) and the Plum Pox Virus protease (PPVp) (**Figure 2d**). Our results showed that CATCH-TEVp is orthogonal to the rapamycin- and ABA-responsive split proteases previously reported,^28,36^ opening great prospects to create chemically-controlled cascades of events.

To demonstrate the utility of CATCH-TEVp for the control of cellular processes, we designed a TEVp activatable transcription factor. We anchored the transcriptional co-activator GAL4-VP16 at the plasma membrane by fusing it to the dopamine receptor D1 (DRD1) with a linker composed of ^FIRE^tag-NTEVp followed by a TEVp cleavage site (**Figure 2e,f**). Match-induced interaction with cytosolically expressed ^FIRE^mate-CTEVp leads to GAL4-VP16 release and translocation to the nucleus, where it activates the transcription of a UAS-controlled gene. To assess the effectiveness of our system, we compared it to a previously reported dopamine-sensitive system in which split TEVp reconstitution is triggered by dopamine-induced interaction with β-arrestin-2.^34^ After 24 h of activation, we measured the UAS-controlled expression of Luc2P (a Luc2 variant bearing a degron sequence). Match-induced activation of CATCH-TEVp led to a 4-fold increase in Luc2P expression, while the dopamine-activated system achieved only a 2-fold increase (**Figure 2g**), suggesting enhanced efficiency of our approach in modulating gene expression. Quantification of Luc2P expression level before and after match treatment by Western blot analysis confirmed our ability to control gene expression upon match-induced activation of CATCH-TEVp (**Figure 2h**). To demonstrate that our CATCH-TEVp-based expression system could be used to control the expression of other proteins, we replaced the gene coding for Luc2 by the gene coding for d2EGFP (an EGFP variant containing a degron).^37^ Confocal microscopy analysis allowed us to show that d2EGFP expression increase by 3-fold upon match addition, in agreement with the dynamic range observed in the bioluminescence assay (**Figure 2i-j**).

### Chemically responsive DNA recombinase

Next, we developed chemically responsive DNA recombinases by leveraging the split CRE recombinase. The CRE recombinase enables gene editing for the study of gene functions. It catalyzes site-specific recombination between two DNA sequences called *loxP* sites, enabling gain or loss of function by sequence removal. Split versions controllable by the addition of rapamycin^38^ or the use of light^39,40^ have been previously described. These activatable CRE allowed precise control of gene recombination in cells and organisms.^38–40^ We developed CATCH-CRE – a split CRE controllable by CATCHFIRE – by fusing ^FIRE^tag at the N-terminus of the C-terminal domain of CRE and ^FIRE^mate at the C-terminus of its N-terminal domain. We constructed a bicistronic plasmid containing a P2A sequence for co-expression of the two parts. To study the responsiveness of CATCH-CRE, we used a target plasmid containing a *loxP*-STOP-*loxP* cassette upstream of Luc2 gene.^39^ CRE-induced removal of the STOP codon enables permanent Luc2 expression (**Figure 3**). Using a 200:1 target:CATCH-CRE plasmid ratio, we observed a 13-fold increase in Luc2 expression (**Figure 3c**), reaching 80% of the expression obtained with full length CRE used as positive control. These results were confirmed by Western blot analysis (**Figure 3d**). To further demonstrate the use of CATCH-CRE for the control of gene recombination, we used CATCH-CRE to chemically control the expression of the red fluorescent protein mCherry using a *loxP*-STOP-*loxP*-mCherry target plasmid. Analysis by confocal fluorescence microscopy showed efficient match-induced gene recombination when using a 200:1 target:CATCH-CRE plasmid ratio, reaching efficiency of recombination corresponding to 27% of that observed with full length CRE (**Figure 3e-f**).

**Figure 3.**
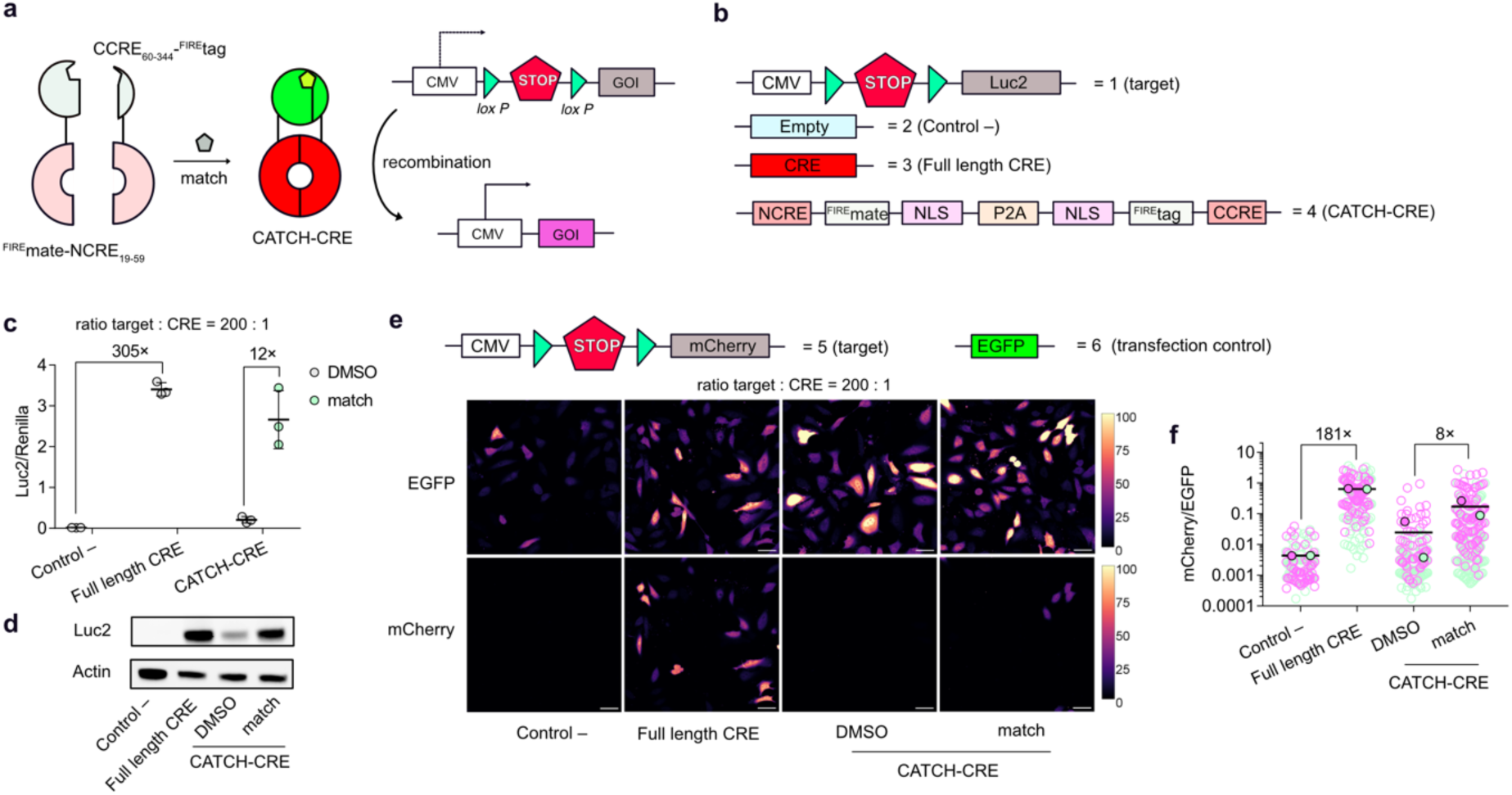
Chemically responsive DNA recombinase. **a** Principle of CATCH-CRE-based DNA editing. **b** Constructs used in this study. **c** HeLa cells transfected with the *loxP*-STOP-*loxP*-Luc2 plasmid and the CATCH-CRE plasmid (200:1 ratio) were incubated for 24 h with 0.5% DMSO or 5 μM of match_550_. Values are shown as mean ± SD of n = 3 independent experiments (each composed of two technical replicates). Control experiments with no CRE (control –) or full-length CRE are also shown. Luc2 signals were normalized by the signal of a co-expressed Renilla luciferase to overcome differences in transfection efficiency. **d** Western blot showing Luc2 expression after 24 h of incubation with DMSO 0.5% or 5 µM of match_550_. The experiment shows one representative blot out of two independent experiments. **e,f** Evaluation of mCherry expression upon activation of CATCH-CRE by confocal microscopy after 24h of incubation with DMSO 0.5% or 5 µM of match_550_. **e** Representative confocal micrographs showing mCherry expression. EGFP was used as a transfection control. Images were taken with identical microscope settings. Scale bars represent 50 µm. **f** mCherry fluorescence intensity (normalized by EGFP signal) of single cells. Each cell is color-coded according to the biological replicate it came from. The solid circles correspond to the mean of each biological replicate. The black line represents the mean values of the two biological replicates (each with n ≥ 50 cells).

### Chemically responsive transcription activators

To further validate CATCHFIRE for the design of chemically responsive control systems, we next engineered chemically responsive transcription activators by leveraging the mammalian two-hybrid system.^41^ Interaction-induced assembly of the DNA-binding domain GAL4 and the VP16 transcription activation domain enables to induce the expression of genes under the control of a UAS sequence.^41^ This approach has been used (i) to detect protein-protein interactions such as the MyoD/Id interaction,^41^ or leucine zippers,^42^ (ii) to create chemically responsive transcription activators using ABA-induced PYL1/ABI interaction^2,25^ or rapalog-induced FRB/FKBP interaction^2,24,43^ or (iii) to design light-responsive transcription activators using the light-induced Cry2-CIB1 interaction.^44^

To design controllable transcription activators, we fused ^FIRE^tag and ^FIRE^mate to GAL4 and VP16 (**Supplementary Figure 4**). Formation of the GAL4-VP16 assembly was assessed by monitoring the expression of a Luc2P gene under the control of a UAS sequence 24 h after addition of match (**Figure 4a-c** and **Supplementary Figure 1**). The best system – composed of GAL4-^FIRE^tag and VP16-^FIRE^mate (hereafter called CATCH-ON v1) – achieved a 24-fold increase in Luc2P expression upon match treatment, reaching expression level comparable to the MyoD/Id positive control,^41^ and displayed very low leakiness in absence of match (**Figure 4c**). The efficiency and low leakiness of CATCH-ON v1 was further confirmed by Western blot analysis (**Figure 4e**). Kinetics analysis showed that expression of Luc2P peaked 7 h after match addition (**Figure 4d**), similarly to what was previously observed with similar expression systems.^26,41^ We confirmed the ability of CATCH-ON v1 to chemically control gene expression by quantifying the match-induced expression of a d2EGFP gene under the control of a UAS sequence (**Figure 4f-g**).

**Figure 4.**
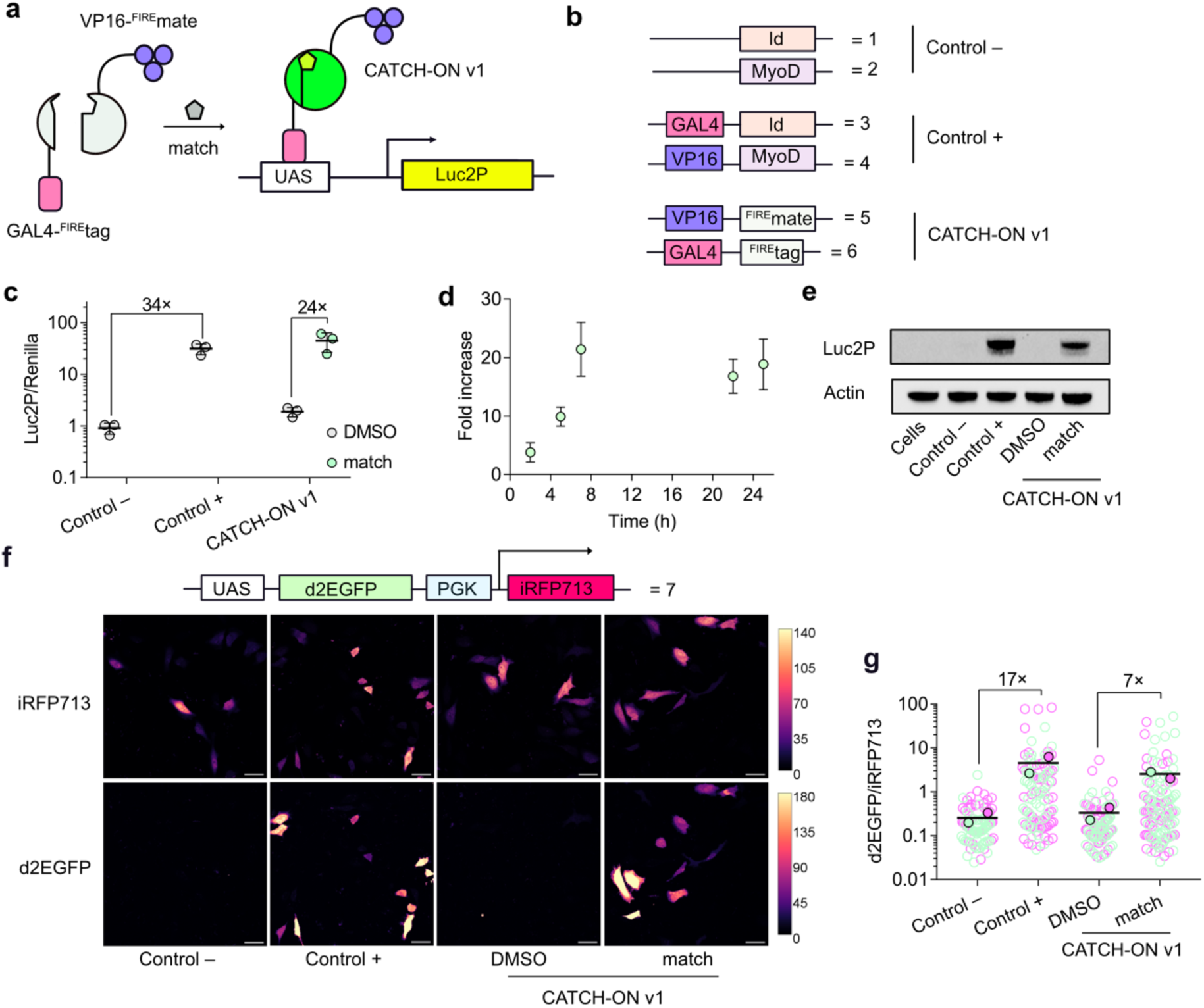
Chemical control of gene expression. **a** Principle of CATCH-ON v1. **b** Constructs used in this study. **c** HeLa cells transfected with UAS-Luc2P plasmid and the CATCH-ON v1 plasmid were incubated with DMSO 0.5% or 5 µM of match_550_ for 24 h. Results are depicted as a normalized signal of Luc2P *vs.* Renilla signal used as a transfection control. **d** Time-lapse of Luc2P gene expression upon activation of CATCH-ON v1. Expression is shown as a fold increase in the normalized Luc2P signal of match_550_-treated *vs*. DMSO-treated cells. Values are shown as mean ± SD of n = 3 independent experiments (each composed of two technical replicates). **e** Western blot showing Luc2P expression controlled with CATCH-ON v1. Transfected cells were incubated with DMSO 0.5% or 5 µM of match_550_ for 24 h. The experiment shows one representative Western blot out of two independent experiments. **f,g** Evaluation of CATCH-ON v1-controlled d2EGFP expression by confocal microscopy after 16 h of incubation with DMSO 0.5% or 5 µM of match_550_. **f** Representative confocal micrographs showing d2EGFP expression. iRFP713 was used as transfection control. Images were taken with identical microscope settings. Scale bars represent 50 µm. **g** d2EGFP fluorescence intensity (normalized by iRFP713 signal) of single cells. Each cell is color-coded according to the biological replicate it came from. The solid circles correspond to the mean of each biological replicate. The black line represents the mean values of the two biological replicates (each with n ≥ 35 cells).

To further improve the system, we created a new set of pairs using two types of modification. First, we replaced GAL4 by GAL4Δ (GAL4_1-65_), a variant with a truncated dimerization domain.^45^ Second, we replaced VP16 by p65Δ (p65_285-550_), a truncated version of the transcription activator p65 lacking the Rel Homology Domain (RHD) responsible for DNA binding and homo and heterodimerization (**Supplementary Figure 2** and **Figure 5**).^44,46^ The combination of GAL4Δ and p65Δ was previously shown to increase expression efficiency in Cry2-CIB1 optogenetics-based systems.^44^ By systematically evaluating the different combinations, we found that substituting VP16 by p65Δ significantly improved the system’s performance. The resulting system composed of GAL4-^FIRE^tag and p65Δ-^FIRE^mate – hereafter called CATCH-ON v2 – achieved a remarkable 160-fold increase of Luc2P expression upon match addition, similarly to other rapamycin-based systems described in the literature,^26,43^ while reaching, in addition, expression levels one order of magnitude higher than CATCH-ON v1 (**Figure 5a-c**). Activation with various concentrations of match showed that gene expression could be finely regulated in a dose-dependent manner (**Figure 5d**). Monitoring expression of Luc2P over 24 h allowed us to characterize the kinetics of expression upon match addition (**Figure 5e**). We showed furthermore that match washout enables to switch off CATCH-ON v2 with a halftime of about 4 h, demonstrating the high tunability and reversibility of the system (**Figure 5e**). The reversibility of CATCH-ON v2 was confirmed by Western blot analysis (**Figure 5f**). We further confirmed the superior performances of CATCH-ON v2 over CATCH-ON v1 by chemically controlling the expression of d2EGFP (**Figure 5g-h**). In accordance with the bioluminescence assay, we observed a 200-fold increase in d2EGFP expression level upon match addition (versus 20-fold with CATCH-ON v1), reaching 10-fold higher expression level.

**Figure 5.**
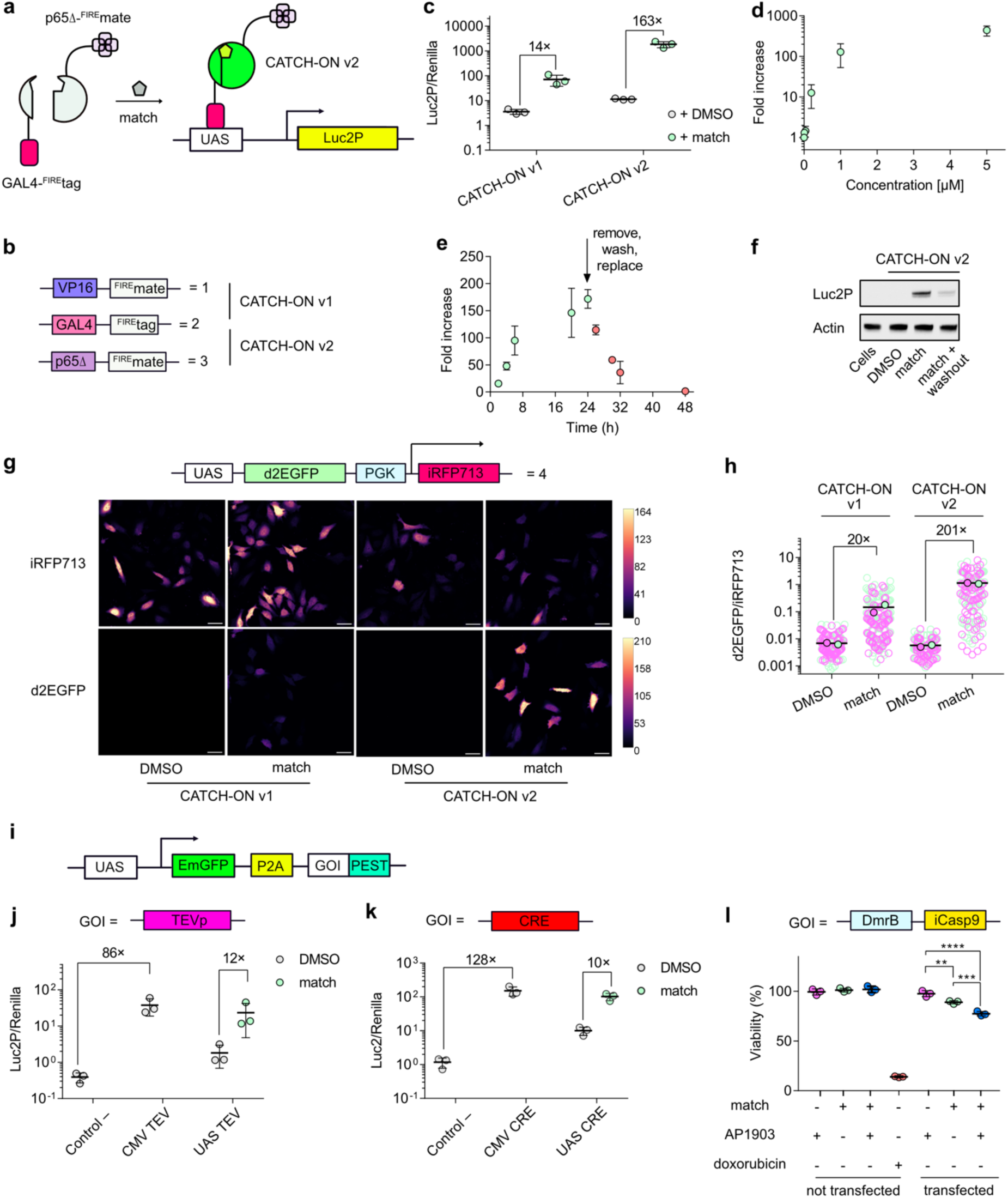
Chemical control of gene expression with improved CATCH-ON. **a** Principle of CATCH-ON v2. **b** Constructs used in this study. **c** HeLa cells transfected with UAS-Luc2 plasmid and CATCH-ON v1 or v2 plasmids were incubated with DMSO 0.5% or match_550_ 5 µM for 24 h. Results are depicted as a normalized signal of Luc2P *vs.* Renilla signal used as a transfection control. **d** Dose response of Luc2P gene expression after 26 h of incubation with DMSO 0.5% or 0-5 µM of match_550_. **e** Kinetics of Luc2P gene expression shown as a fold increase in the normalized Luc2P signal of match_550_(5 µM)-treated *vs*. DMSO-treated cells. Values are shown as mean ± SD of n = 3 independent experiments (each composed of two technical replicates). **f** Characterization of CATCH-ON v2 by Western blot. Cells transfected with UAS-Luc2 plasmid and CATCH-ON v2 plasmids were incubated with DMSO 0.5% or match_550_ 5 µM for 24 h. For the reversibility experiment, the media was removed and replaced with fresh media and incubated for 8 h. The experiment shows one representative blot out of two independent experiments. **g,h** Evaluation of CATCH-ON-controlled d2EGFP expression by confocal microscopy after 16 h of incubation with DMSO 0.5% or 5 µM of match_550_. **g** Representative confocal micrographs showing d2EGFP expression. iRFP713 was used as transfection control. Images were taken with identical microscope settings. Scale bars represent 50 µm. **h** d2EGFP fluorescence intensity (normalized by iRFP713 signal) of single cells. Each cell is color-coded according to the biological replicate it came from. The solid circles correspond to the mean of each biological replicate. The black line represents the mean values of the two biological replicates (each with n ≥ 50 cells). **i-l** Control of the expression of biologically relevant proteins using the CATCH-ON v2 system**. i** Generic plasmid for inducible UAS-based gene expression. **j** TEVp protease and **k** CRE recombinase activity in HeLa cells upon activation of CATCH-ON v2 with DMSO 0.5% or 5 µM of match_550_. Their activity was measured after 24 h. **l** Viability of HeLa cells transfected with UAS-DmrB-iCasp9 plasmid and CATCH-ON v2 plasmids after treatment with 5 µM of match_550_ and 5 µM of AP1903 for 16 h. Induction of cell death with 5 µM of doxorubicin was used as a positive control (For normalization details, see Materials and Methods section). Values are shown as mean ± SD of n = 3 independent experiments (each composed of two technical replicates). ONE-way ANOVA analysis followed by a Sidak’s multiple comparisons test was performed where *p*-values are 0.018, 0.001 and <0.001 for **, *** and **** respectively.

To showcase the use of CATCH-ON v2 for the chemical regulation of cellular functions, we tested its use to conditionally control the expression of full-length TEVp and CRE (**Figure 5i-k**). The expression of full-length TEVp was evaluated by monitoring the activation of the TEVp bioluminescent sensor described above. We observed a 12-fold increased activity of full-length TEVp upon match addition (**Figure 5j**), reaching activity comparable to that obtained with constitutively expressed TEVp. We evaluated full-length CRE expression by monitoring the expression of Luc2 upon excision of an upstream *loxP*-STOP-*loxP* cassette. We observed a 10-fold increased activity of full-length CRE upon match addition, reaching activity comparable to that obtained with constitutively expressed CRE recombinase (**Figure 5k**), further demonstrating the efficacy of CATCH-ON v2.

One exciting application of chemically controlling gene expression is to control the function of therapeutic cells on demand. To explore this idea, we used CATCH-ON v2 to chemically control the expression of DmrB-iCasp9 – a suicide switch previously used to conditionally control CAR-T cell activity.^18,20,47,48^ Made of a fusion of the truncated caspase iCasp9 and DmrB, an FKBP variant that can homodimerize in the presence of the rapalog AP1903, this suicide switch triggers the apoptotic cascade upon addition of AP1903, leading to cell death. We generated a construct with the gene coding for DmrB-iCasp9 downstream of a UAS sequence (**Figure 5i,l**). Control of its expression with CATCH-ON v2 allowed us to create an AND logic gate enabling to tightly control cell death. We experimentally showed that the presence of both match and AP1903 was required to significantly reduce cell viability in transfected cells (**Figure 5l**). It is important to mention here that, in the experiment shown on **Figure 5l**, DmrB-iCasp9 is expressed transiently, therefore only a subpopulation of cells expresses the protein and can thus be chemically controlled, while our viability test assayed the viability of all cells, explaining why the decrease in viability is only partial. The use of such AND logic gate enabled to get rid of the intrinsic leakiness of DmrB-iCasp9 in absence of AP1903. This experiment opens exciting prospects for applying CATCH-ON in cell therapy. It additionally confirmed the compatibility of CATCHFIRE with other described CID technologies, opening the possibility for the creation of various Boolean systems.

To further explore the potential of CATCH-ON for cell-based therapies, we used CATCH-ON v2 to chemically control the secretion of a therapeutically relevant protein. As a model, we focused on insulin used for regulating blood glucose level in diabetes treatment. To monitor insulin secretion, we used a chimeric protein incorporating NanoLuc as a secretion reporter.^49^ In this design, NanoLuc was flanked by the peptides A and B of insulin using Furin cleavage sites as linkers. Once synthesized and marked for secretion, this chimera travels from the endoplasmic reticulum to the trans-Golgi network where the Furin enzyme cleaves its cleavage site, releasing NanoLuc and functional insulin (**Figure 6a**).^1,35^ The NanoLuc secreted is used as a proxy for insulin secretion. CATCH-ON v2 allowed us to chemically control insulin secretion in a match-dependent manner. Treatment of cells with match increased insulin secretion by 55-fold, reaching levels comparable with constitutive insulin secretion (**Figure 6b,c**). This experiment indicates that CATCH-ON v2 could be used in the future for controlling the secretion of various therapeutically relevant proteins such as chimeric antigen receptors (CAR), cytokines or hormones.

**Figure 6.**
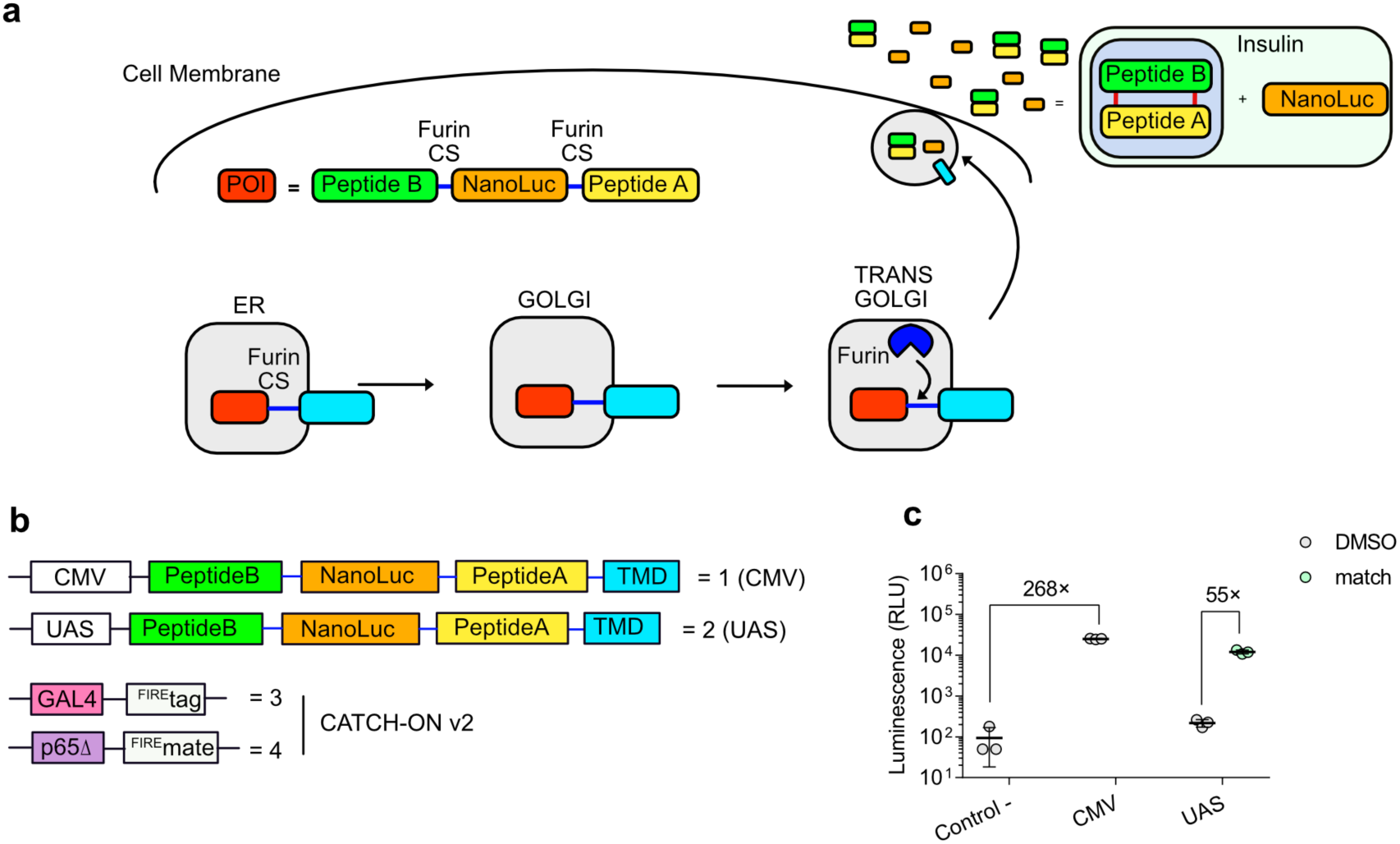
Control of insulin secretion with CATCH-ON. **a** Insulin/NanoLuc secretion scheme and **b** constructs used in this study. **c** HeLa cells were transfected and incubated with DMSO 0.5% or 5 µM of match_550_ for 24 h. The supernatant was collected and NanoLuc activity was measured. Values are shown as mean ± SD of n = 3 independent experiment.

## DISCUSSION

The development of chemically responsive protein switches offers a highly versatile and powerful tool for precise control of cellular processes, with significant implications across basic research, biotechnology and biotherapy. An ideal chemically responsive protein switch should combine versatility, rapid responsiveness, reversibility, and non-toxicity, while maintaining robust efficiency and compatibility with existing CID technologies. In this study, we show that the CATCHFIRE technology fulfils these criteria, enabling the modular design of chemically responsive switches through the controlled assembly of split enzymes, such as luciferases, the TEV protease and the CRE recombinase. This technology not only facilitates finely tuned modulation of cellular activities on demand but also provides a flexible framework for extension to other enzymatic systems.

Although the CATCH-TEVp and CATCH-CRE systems presented here have demonstrated robust performance, there is still some opportunity for further optimization. To date our work focused on previously established topologies; however, the dynamic range of these systems could be further enhanced by minimizing self-complementation among split components. Further improvements could be achieved by exploring alternative topologies, leveraging advances in de novo protein design,^50^ implementing directed protein evolution, or employing diverse protein homologs, as we recently did for optimizing split fluorescent reporters.^51^

Targeting multidomain proteins for splitting, as opposed to monodomain ones, simplifies the design of chemically responsive protein-based switches with high efficiency and minimal leakiness. The CATCH-ON system exemplifies this principle, providing near-undetectable basal activation and a remarkable dynamic range – up to 200-fold for CATCH-ON v2 – enabling dose-dependent and reversible gene expression control. CATCH-ON provides a generalizable platform, capable of regulating a wide spectrum of proteins including luciferases, fluorescent proteins, proteases, DNA recombinase. Its compatibility with existing CID technologies paves the way for sophisticated Boolean logic systems, opening exciting prospects for synthetic biology and biotechnology.

The precise, tunable, and reversible gene control afforded by CATCH-ON is especially advantageous in scenarios where conventional inducible gene expression systems, such as doxycycline-based platforms, may be unsuitable due to off-target effects. Doxycycline, for instance, has been shown, to negatively impact mammalian cell proliferation, disrupt mitochondrial protein homeostasis, trigger the unfolded protein response, induce cell cycle arrest and apoptosis, and even cause transcriptional changes,^52^ raising concerns for sensitive and therapeutic applications.^53^ In contrast, CATCH-ON relies on a fully synthetic dimerizer, designed to exhibit no adverse effects on cell viability and should display no off-target biological activity, thus minimizing unintended cellular effects, making CATCH-ON a powerful and flexible alternative to conventional inducible gene expression system. We demonstrate the ability to regulate suicide switches or therapeutic proteins like insulin, paving the way for controllable cellular therapies with enhanced control and increased safety. Although the road towards such applications is long and will require further validation, particularly demonstrating robust in vivo functionality and efficacy in therapeutic cells, the versatility and performance of CATCHFIRE and its derivatives mark a significant step forward in the design of smart, responsive cellular control systems.

## ACKNOWLEDGMENTS

pUAS-NanoLuc was a gift from Robert Campbell (Addgene plasmid # 87696), pCMV-IRES-Renilla Luciferase-IRES-Gateway-Firefly Luciferase (pIRIGF) was a gift from William Kaelin (Addgene plasmid # 101139), pcDNA3.1 TEV (full-length) was a gift from Xiaokun Shu (Addgene plasmid # 64276), FRB_nSuMMVp was a gift from Roman Jerala (Addgene plasmid # 118970), FRB_nSuMMVp was a gift from Roman Jerala (Addgene plasmid # 118970), cycLuc_TEVS was a gift from Roman Jerala (Addgene plasmid # 119207), cycLuc_SuMMVs was a gift from Roman Jerala (Addgene plasmid # 119210), PYL1_nPPVp was a gift from Roman Jerala (Addgene plasmid # 119211), ABI_cPPVp was a gift from Roman Jerala (Addgene plasmid # 119212), ABI_cPPVp was a gift from Roman Jerala (Addgene plasmid # 119212), PYL1_nTEVp was a gift from Roman Jerala (Addgene plasmid # 119213), pcDNA3_DRD1-NTEV-TCS-GV-2xHA was a gift from Michael Wehr (Addgene plasmid # 194358), pcDNA3.1_Zeo_ARRB2-1-383-CTEV-2xHA was a gift from Michael Wehr (Addgene plasmid # 194382); pGL4_10xUAS-MLPmin-luc2 was a gift from Michael Wehr (Addgene plasmid # 194383), pCMV-IRES-Renilla Luciferase-IRES-Gateway-Firefly Luciferase (pIRIGF) was a gift from William Kaelin (Addgene plasmid # 101139), pBS185 CMV-Cre was a gift from Brian Sauer (Addgene plasmid # 11916), pcDNA3.1_PA-Cre was a gift from Moritoshi Sato (Addgene plasmid # 122960), Proinsulin-NanoLuc in pLX304 was a gift from David Altshuler (Addgene plasmid # 62057), pCMMP-MCS-IRES-eGFP-P2A-iCas9 was a gift from Ryoji Yao (Addgene plasmid # 178533), FURIN-bio-His was a gift from Gavin Wright (Addgene plasmid # 51755), pHR_Gal4UAS-dGFP_PGK_iRFP (pLZA042) was a gift from Jared Toettcher (Addgene plasmid # 203908).

We thank the microscopy facility of the Institut de Biologie Paris Seine of Sorbonne University, and more particularly France Lam and Chloé Chaumeton for their assistance. This work has been supported by the Agence Nationale de la Recherche (ANR-23-CE44-0014-01CATCHFIRE), a prematuration grant from Sorbonne University and the Institut Universitaire de France. ND thanks the École Normale Supérieure for PhD funding.

## AUTHOR CONTRIBUTIONS

J.F.P and A.G. designed the overall project and wrote the paper with the help of the other authors. J.F.P., N.D., S.D., F.P. and A.G. designed the experiments. J.F.P., N.D., S.D. performed the experiments. J.F.P., N.D., F.P. and A.G. analyzed the experiments.

## MATERIALS AND METHODS

### Plasmid construction

Synthetic oligonucleotides used for cloning were purchased from Integrated DNA Technology. PCR reactions were performed with Q5® polymerase (New England Biolabs) in the buffer provided. PCR products were purified using QIAquick® PCR purification kit or QIAquick® Gel Extraction Kit (QIAGEN). Isothermal assemblies (Gibson Assembly) were performed using a homemade mix prepared according to previously described protocols.^54^ GAL4-Id VP16-MyoD plasmid were bought from PROMEGA as the CheckMate™ Mammalian Two-Hybrid System. Small-scale isolation of plasmid DNA was conducted using a QIAprep miniprep kit (QIAGEN) from 2-5 mL overnight DH10β in LB culture supplemented with appropriate antibiotic. Large-scale isolation of plasmid DNA was conducted using the QIAprep maxiprep kit (QIAGEN) from 150-180 mL overnight bacterial culture supplemented with the corresponding selection antibiotics. All plasmid sequences were confirmed by Sanger sequencing with appropriate sequencing primers (Eurofins). See **Supplementary Information** and **Supplementary Table 1** for ORFs and primers used to synthesize them, and **Supplementary Table 2** for primers sequence.

### Cell culture experiments

#### General considerations

HeLa cells (ATCC, CCL2 Lot 700756809) were grown in Minimal Essential Medium media (MEM, Gibco™) supplemented with 10% (v/v) Fetal Bovine Serum (FBS, Gibco™), 1% (v/v) of Sodium Pyruvate (Gibco™) and 1% (v/v) of Penicillin-Streptomycin (Sigma-Aldrich) at 37 °C in a 5% CO_2_ atmosphere.

#### Ligands

Match_550_ (a.k.a. HBR-2,5DM) (in-house synthesized),^55^ rapamycin (TCI), abscisic acid (Sigma-Aldrich), dopamine (Sigma-Aldrich), doxorubicin (Sigma-Aldrich), AP1903 (TOCRIS) were dissolved in DMSO (Sigma-Aldrich) at 3 to 20 mM stock solutions and kept at –20 °C. Intermediate solutions were prepared at 200-400× final concentration in DMSO.

### Chemically responsive Luciferase study

130 µL of a 2 × 10^4^ cells/mL suspension were added in a 96-white well plate (Corning®) and incubated for 24 h in the abovementioned conditions. Each sample to be added to the wells for transfection contains half volume of GeneJuice® (Sigma-Aldrich): in OptiMEM™ (Gibco™, 8% v/v) and half volume of DNA: Opti-MEM™ that was incubated for 15 to 20 minutes at RT before addition to the wells according to GeneJuice® protocol. For each well, cells were transfected with 6.3 µL of the mixture to obtain a final amount of 20 ng of DNA per well. Cells were incubated for 24 h, media was removed and replaced with 100 µL of OptiMEM™ containing a final concentration of DMSO 0.5% (v/v), match_550_ 5 µM, or rapamycin 0.5 µM, and then incubated for 2 h. Cells were treated with Nano-Glo® Live Cell Assay System (PROMEGA) as described in the protocol. Luminescence was recorded with a POLARstar OPTIMA plate reader (BMG Biotech) at 37 °C for 90 minutes with a 2-minute scan interval and 1-second integration time. Results show the luminescence values 40 minutes after reagent addition. Values are shown as mean ± SD of n ≥ 3 independent experiments (each composed of two technical replicates).

### CATCH-TEVp and orthogonality with other inducible proteases

130 µL of a 4 × 10^4^ cells/mL suspension was added in a 96-white well plate and incubated for 24 h in the abovementioned conditions. For each well, cells were transfected as reported above with 30 ng of a circularly permuted Firefly Luciferase (Luc2) encoding plasmid (target plasmid), 15 ng of each plasmid encoding the inducible split protease system, and 2.5 ng of a Renilla encoding plasmid used as a transfection control. After for 24 h, media was removed and replaced with 200 µL of fresh media containing DMSO 0.5%, Absicic Acid (ABA) 100 µM, rapamycin 3 µM or match_550_ 5 µM, and cells were incubated for 1 h at 37 °C. Luciferase activity was measured using the Dual-Luciferase® Reporter System (PROMEGA). Briefly, media was removed, and wells were washed with 150 µL of DPBS. The supernatant was removed, and cells were lysed with 20 µL per well of Passive Lysis Buffer (PROMEGA) while gently rocking at room temperature for 20 minutes. Luciferase activity was measured as indicated by the supplier. Luminescence was recorded 2-3 minutes after reagent addition, using a 1-second integration time at 25 °C. Values represent the mean of the Luc2 signal divided by the signal of the Renilla luciferase signal, constitutively expressed and used as a transfection reporter, ± SD of n ≥ 3 independent experiments (each composed of two technical replicates). Fold increase represents the mean of the normalized signal of cells treated with match, rapamycin or ABA *vs.* cells treated with DMSO.

### Gene expression induced by CATCH-TEVp

130 µL of a 4 × 10^4^ cells/mL suspension was added in a 96-white well plate and incubated for 24 h in the abovementioned conditions. For each well, cells were transfected as reported above with, in each well, 30 ng of target plasmid (UAS-Luc2P plasmid), 15 ng of each plasmid encoding the inducible splitTEVp system and 2.5 ng of Renilla encoding plasmid as transfection control. DMSO 0.5%, dopamine 5 µM or match_550_ 5 µM were added and cells were incubated for 24 h. Luc2P expression was studied, gene expression was measured by using the Dual-Luciferase® Reporter System as indicated by the supplier and as described before. Fold increase represents the mean of the Luc2P signal divided by the signal of a Renilla protein used as reporter, of dopamine or match treated *vs.* DMSO treated cells, ± SD of n = 3 independent experiments (each composed of two technical replicates). One-Way ANOVA analysis followed by a Tukey’s multiple comparison was performed.

### Control of luciferase expression using CATCH-CRE

130 µL of a 4 × 10^4^ cells/mL suspension was added in a 96-white well plate and incubated for 24 h in the abovementioned conditions. For each well, cells were transfected as reported above using 50 ng of *loxP*_STOP_*loxP*_Luc2 target plasmid in each well, 0.25 ng of CATCH-CRE encoding plasmid, CMV_CRE or empty plasmid and 2.5 ng of Renilla encoding plasmid as transfection control. After 24 h, media was removed and replaced with 150 µL of fresh media containing DMSO 0.5% or match_550_ 5 µM and incubated for 24 h. As indicated before, Luc2 expression was measured using the Dual-Luciferase® Reporter System. Values are shown as mean ± SD of 3 independent experiments (each composed of two technical replicates).

### Control of gene expression with CATCH-ON

#### Control of luciferase expression with CATCH-ON

130 µL of a 4 × 10^4^ cells/mL suspension was added in a 96-white well plate and incubated for 24 h in the abovementioned conditions. For each well, cells were transfected with 30 ng of reporter (UAS-Luc2P plasmid) and 15 ng of each plasmid encoding CATCH-ON parts or the MyoD/Id parts were added. DMSO 0.5% or match_550_ 5 µM final concentration were added and co-incubated for 2–26 h with the transfection mixture. For degradation experiments, media was removed, wells were washed with 150 µL of DPBS and replaced by 150 µL of fresh media. Luciferase expression was measured by using the Dual-Luciferase® Reporter System as indicated above. Values represent the mean of the Luc2P signal divided by the signal of the Renilla protein used as a transfection reporter, ± S. D of n = 3 independent experiments (composed of two technical replicates).

#### Control of a suicide gene with CATCH-ON

130 µL of a suspension of 6 × 10^4^ cells/mL was added in a 96-white well plate and incubated for 24 h in the abovementioned conditions. For each well, cells were transfected with 20 ng of target (UAS-DmrB-iCasp9 plasmid) and 10 ng of each plasmid encoding CATCH-ON parts for 8h, media was removed and replaced by 200 µL of fresh media containing DMSO 0.25%, match_550_ 5 µM, AP1903 5 µM or the combination of both as final concentration. To control molecules’ cytotoxicity, non-transfected cells were treated equally, using doxorubicin at 5 µM as the positive control. Cells were treated for 16 h and viability was measured using Cell-Titer Glo® (PROMEGA) where 100 µL were removed from the well and the remaining supernatant was treated with 100 µL of reagent, the mixture was incubated at 30 minutes at room temperature. Luminescence was recorded at 25 °C using a 1-second integration time. Values are shown as mean ± SD of 3 independent experiments (each composed of two technical replicates). Viability (%) is expressed as V = 100 × (Luminescence treatment – Luminescence Blank) / (Luminescence DMSO control – Luminescence Blank).

#### Control of TEVp expression with CATCH-ON

130 µL of a suspension of 4 × 10^4^ cells/mL was added in a 96-white well plate and incubated for 24 h in the abovementioned conditions. For each well, cells were transfected as reported above where in each well with 30 ng of a plasmid containing a circularly permuted Luc2 used as a target and 10 ng of GAL4-^FIRE^tag plasmid that contains also a Renilla cassette used as a transfection control. For the treatments were added for positive control 10 ng of CMV_TEVp encoding plasmid and 10 ng of an EGFP encoding plasmid (as mock plasmid), for negative control 10 ng of empty plasmid and 10 ng of EGFP encoding plasmid and for the test 10 ng of UAS-TEVp encoding plasmid and 10 ng of p65Δ-^FIRE^mate encoding plasmid. The wells were co-incubated with DMSO 0.5% or match_550_ at 5 µM for 24 h. TEVp activity was assessed by measuring Luc2 expression using the Dual-Luciferase® Reporter System (PROMEGA). Values are shown as mean ± SD of 3 independent experiments (each composed of two technical replicates).

#### Control of CRE expression with CATCH-ON

130 µL of a suspension of 4 × 10^4^ cells/mL was added in a 96-white well plate and incubated for 24 h in the abovementioned conditions. For each well, cells were transfected as reported above with 50 ng of target plasmid and 5 ng of GAL4-^FIRE^tag plasmid. For the treatments were added for positive control 0.05 ng of CMV_CRE plasmid and 5 ng of EGFP encoding plasmid (as mock plasmid), for negative control 0.05 ng of empty plasmid and 5 ng of EGFP encoding plasmid (as mock plasmid) and for the test 0.05 ng of UAS-CRE plasmid and 5 ng of p65Δ-^FIRE^mate plasmid. The cells were cultured for 24 h, media was removed and replaced with fresh media containing DMSO 0.5% or match_550_ at 5 µM for 24 h as above. CRE activity was assessed by measuring Firefly signal as described above using the Dual-Luciferase® Reporter System. Values are shown as mean ± SD of 3 independent experiments (each composed of two technical replicates).

#### Control of insulin secretion with CATCH-ON (NanoLuc measurement)

1 mL of a 2.5 × 10^5^ cells/mL suspension was added to a 12-well plate and incubated for 24 h under the aforementioned conditions. The media was removed and replaced with 930 µL of fresh MEM, then completed with 70 µL of a transfection mixture containing 680 ng of DNA. For all conditions, 280 ng of Furin encoding DNA were added; for the negative control, 400 ng of EGFP encoding plasmid (as mock plasmid) were used. For the positive control, 100 ng of CMV_Insulin-NanoLuc encoding plasmid and 300 ng of EGFP encoding plasmid (as mock plasmid) were used. For the test, 100 ng of UAS-insulin-NanoLuc encoding plasmid, 100 ng of GAL4-^FIRE^tag encoding plasmid, 100 ng of p65Δ-^FIRE^mate encoding plasmid, and 100 ng of EGFP encoding plasmid (as mock plasmid) were used. Additionally, DMSO at 0.5% or match_550_ at a final concentration of 5 µM was added and co-incubated for 24 h with the transfection mixture. The media was collected and centrifuged for 5 min at 500 g rpm at 4°C. NanoLuc secretion was measured at room temperature from 100 µL of the supernatant using the Nano-Glo® Luciferase system with a 1-second integration time. Values are shown as mean ± SD of 3 independent experiments.

### Western Blot analysis

2 mL of a HeLa cells suspension (1-2.5 × 10^5^ cells/mL) were seeded in a 6-well plate and incubated for 24 h in the described conditions. Media was removed and replaced with 1.9 mL of fresh media and 100 µL of transfection solution containing 1 µg of DNA per well. See **Supplementary Table 3** for transfection mixture. For CATCH-TEVp and CATCH-ON experiments, cells were co-incubated with DMSO 0.5% or match_550_ at 5 µM. For CATCH-CRE experiments, cells were transfected for 24 h, media was removed and replaced by 2 mL of fresh one containing DMSO 0.5% or match_550_ at 5 µM. After 24 hours, the media was removed, and the cells were washed with DPBS (2 × 1 mL). Lysates were obtained by adding 200 µL of a mixture 1:100 of a protease cocktail (Halt™ Protease Inhibitor Cocktail, 100X, Thermo Scientific™) and RIPA lysis and extraction buffer (Thermo Scientific™) on ice for 5 minutes. The supernatant containing the proteins was collected after centrifugation (15000 g at 4°C for 30 min). In the case of CATCHFIRE reversibility, media was removed and washed as stated and replaced by 2 mL of media for 8 h. Samples were treated as before. Protein concentrations were evaluated thanks to a calibration curve following Pierce™ BCA Protein Assay Kit (Thermo Scientific™) instructions. The calibration samples (0-2000 µg/mL) were prepared from a Serum Bovine Albumin solution following the instructions provided in the kit. The absorbance was read after 30 min incubation at 37°C at 562 nm on a TECAN Spark® 10 plate reader. Solutions of 10–40 µg of the protein mixture and Laemmli Buffer (5×) were boiled for 5 min at 95°C and loaded onto a gel (Premade NuPAGE™ Bis-Tris, 10% MES buffer, Thermo Fisher Scientific). After 45 min migration at 150 V using a PowerPac™ Basic device (BioRad), the proteins were wet-transferred to a nitrocellulose membrane (Nitrocellulose Blotting membrane 0.2 µm, General Electric) applying current for 90 minutes at 100 V in using Transfer Buffer (Towbin 1×, MeOH 20% (v/v) and milliQ H_2_O). The membranes were blocked in PBS (PB 50 mM, NaCl 150 mM, pH = 7.4) and skimmed milk 5% (w/v) for one hour at RT. After blocking, the membrane was incubated with the primary antibody diluted in PBS + skimmed milk 0.5% (w/v) overnight at 4°C with gentle rocking. The membranes were washed 3 times in PBS + skimmed milk 0.5% (w/v). The membrane was then probed with the secondary antibody diluted in PBS + skimmed milk 5% (w/v) for 45 min at RT. The membranes were washed 1 time in PBS + skimmed milk 0.5% and then 2 times in PBS. The proteins were visualized using the following primary antibodies: monoclonal mouse anti-LgBit (N7100, PROMEGA, 1:500), rabbit anti-FAST (Covalab, 1:1000), and polyclonal rabbit anti-β-actin (4967S, Cell Signaling, 1:1000). Secondary antibodies were used as follow: goat anti-rabbit IgG HRP-linked antibody (7074S, Cell Signalling, 1:2500), goat anti-mouse IgG (H+L) antibody (W4021, PROMEGA, 1:2500). The membranes were revealed by Chemiluminescent detection with Chemi-Doc™ imager (Biorad) after being incubated for one minute in 3-4 mL of a 1:1 solution of SuperSignal West Pico PLUS (Horse Radish Peroxidase) and luminol (Thermo Fisher Scientific). The membranes were stripped by washing them with PBS (2 × 5 mL) and then using Restore™ Western Blot Stripping Buffer (Thermo Fisher Scientific) for 15-20 minutes while shaking at room temperature, then washed again with PBS (2 × 5 mL) and re-probed with the corresponding antibodies.

### Confocal microscopy experiments

A suspension of 4.2 to 6 × 10^4^ HeLa cells in a total volume of 2 mL was seeded in a 35 mm µDish (Ibidi®) and incubated for 24 h in the conditions described before. Medium was removed, replaced with 1.9 mL of fresh media and 100 µL of transfection solution containing 1 µg of DNA per dish. See **Supplementary Table 4** for transfection mixture. For CATCH-TEVp and CATCH-ON experiments, cells were incubated for 8 h and media was removed and replaced by fresh media containing DMSO 0.5% or match_550_ at 5 µM for 16 h. For CATCH-CRE, media was removed after 24 h and replaced by fresh media containing DMSO 0.5% or match_550_ 5 µM for 24 h. After incubation, the media was removed, and the cells were washed with DPBS (2 × 1 mL). 1 mL of fresh Dubelcco’s Modified Eagle Medium (DMEM, Gibco) without phenol red was added and samples were analysed on a Zeiss LSM 980 Laser Scanning Microscope equipped with a 20× or a 63× objective. ZEN software was used to collect the data and then images were analysed using Icy software (v 2.5.2.0) or Fiji software. To track fluorescence signal in the cytoplasm in the CATCH-TEVp and CATCH-ON experiments, cell contours were determined by masking the signal in the iRFP713 channel with the plug-in HK-means with the following parameters: intensity class equals 100, min object size (px) equals 20000, max object size (px) equals 50000. For the CATCH-CRE experiments, cell contours were determined by masking the signal in the EGFP channel with the plug-in HK-means with the following parameters: intensity class equals 75, min object size (px) equals 10000, max object size (px) equals 80000. The ROI’s signal intensity was measured for each channel using plug-in Active Contours.

## REFERENCES

(1) Praznik, A.; Fink, T.; Franko, N.; Lonzarić, J.; Benčina, M.; Jerala, N.; Plaper, T.; Roškar, S.; Jerala, R. Regulation of Protein Secretion through Chemical Regulation of Endoplasmic Reticulum Retention Signal Cleavage. Nature Communications 2022, 13 (1), 1323. 10.1038/s41467-022-28971-9.

(2) Liang, F.-S.; Ho, W. Q.; Crabtree, G. R. Engineering the ABA Plant Stress Pathway for Regulation of Induced Proximity. Science Signaling 2011, 4 (164), rs2. 10.1126/scisignal.2001449.

(3) Gao, Y.; Xiong, X.; Wong, S.; Charles, E. J.; Lim, W. A.; Qi, L. S. Complex Transcriptional Modulation with Orthogonal and Inducible dCas9 Regulators. Nature Methods 2016, 13 (12), 1043–1049. 10.1038/nmeth.4042.

(4) Rihtar, E.; Lebar, T.; Lainšček, D.; Kores, K.; Lešnik, S.; Bren, U.; Jerala, R. Chemically Inducible Split Protein Regulators for Mammalian Cells. Nat Chem Biol 2023, 19 (1), 64–71. 10.1038/s41589-022-01136-x.

(5) Wehr, M. C.; Laage, R.; Bolz, U.; Fischer, T. M.; Grünewald, S.; Scheek, S.; Bach, A.; Nave, K. A.; Rossner, M. J. Monitoring Regulated Protein-Protein Interactions Using Split TEV. Nature Methods 2006, 3 (12), 985–993. 10.1038/nmeth967.

(6) Zheng, Y.; Gao, N.; Fu, Y. L.; Zhang, B. Y.; Li, X. L.; Gupta, P.; Wong, A. J.; Li, T. F.; Han, S. Y. Generation of Regulable EGFRvIII Targeted Chimeric Antigen Receptor T Cells for Adoptive Cell Therapy of Glioblastoma. Biochemical and Biophysical Research Communications 2018, 507 (1–4), 59–66. 10.1016/j.bbrc.2018.10.151.

(7) Wu, C.-Y.; Roybal, K. T.; Puchner, E. M.; Onuffer, J.; Lim, W. A. Remote Control of Therapeutic T Cells through a Small Molecule–Gated Chimeric Receptor. Science 2015, 350 (6258), aab4077. 10.1126/science.aab4077.

(8) Simpson, L. M.; Macartney, T. J.; Nardin, A.; Fulcher, L. J.; Röth, S.; Testa, A.; Maniaci, C.; Ciulli, A.; Ganley, I. G.; Sapkota, G. P. Inducible Degradation of Target Proteins through a Tractable Affinity-Directed Protein Missile System. Cell Chemical Biology 2020, 27 (9), 1164–1180.e5. 10.1016/j.chembiol.2020.06.013.

(9) Hamilton, E. P.; Ma, C.; De Laurentiis, M.; Iwata, H.; Hurvitz, S. A.; Wander, S. A.; Danso, M.; Lu, D. R.; Perkins Smith, J.; Liu, Y.; Tran, L.; Anderson, S.; Campone, M. VERITAC-2: A Phase III Study of Vepdegestrant, a PROTAC ER Degrader, versus Fulvestrant in ER+/HER2-Advanced Breast Cancer. Future Oncology 2024, 20 (32), 2447–2455. 10.1080/14796694.2024.2377530.

(10) Nowak, R. P.; Deangelo, S. L.; Buckley, D.; He, Z.; Donovan, K. A.; An, J.; Safaee, N.; Jedrychowski, M. P.; Ponthier, C. M.; Ishoey, M.; Zhang, T.; Mancias, J. D.; Gray, N. S.; Bradner, J. E.; Fischer, E. S. Plasticity in Binding Confers Selectivity in Ligand-Induced Protein Degradation Article. Nature Chemical Biology 2018, 14 (7), 706–714. 10.1038/s41589-018-0055-y.

(11) Banik, S. M.; Pedram, K.; Wisnovsky, S.; Ahn, G.; Riley, N. M.; Bertozzi, C. R. Lysosome-Targeting Chimaeras for Degradation of Extracellular Proteins. Nature 2020, 584 (7820), 291–297. 10.1038/s41586-020-2545-9.

(12) Wang, W. W.; Chen, L. Y.; Wozniak, J. M.; Jadhav, A. M.; Anderson, H.; Malone, T. E.; Parker, C. G. Targeted Protein Acetylation in Cells Using Heterobifunctional Molecules. Journal of the American Chemical Society 2021, 143 (40), 16700–16708. 10.1021/jacs.1c07850.

(13) Yamazoe, S.; Tom, J.; Fu, Y.; Wu, W.; Zeng, L.; Sun, C.; Liu, Q.; Lin, J.; Lin, K.; Fairbrother, W. J.; Staben, S. T. Heterobifunctional Molecules Induce Dephosphorylation of Kinases-A Proof of Concept Study. Journal of Medicinal Chemistry 2020, 63 (6), 2807–2813. 10.1021/acs.jmedchem.9b01167.

(14) Neklesa, T. K.; Tae, H. S.; Schneekloth, A. R.; Stulberg, M. J.; Corson, T. W.; Sundberg, T. B.; Raina, K.; Holley, S. A.; Crews, C. M. Small-Molecule Hydrophobic Tagging-Induced Degradation of HaloTag Fusion Proteins. Nature Chemical Biology 2011, 7 (8), 538–543. 10.1038/nchembio.597.

(15) Bottone, S.; Joliot, O.; Cakil, Z. V.; El Hajji, L.; Rakotoarison, L.-M.; Boncompain, G.; Perez, F.; Gautier, A. A Fluorogenic Chemically Induced Dimerization Technology for Controlling, Imaging and Sensing Protein Proximity. Nature Methods 2023, 20 (10), 1553–1562. 10.1038/s41592-023-01988-8.

(16) Pecot, M. Y.; Malhotra, V. Golgi Membranes Remain Segregated from the Endoplasmic Reticulum during Mitosis in Mammalian Cells. Cell 2004, 116 (1), 99–107. 10.1016/S0092-8674(03)01068-7.

(17) Zhou, X.; Dotti, G.; Krance, R. A.; Martinez, C. A.; Naik, S.; Kamble, R. T.; Durett, A. G.; Dakhova, O.; Savoldo, B.; Di Stasi, A.; Spencer, D. M.; Lin, Y.-F.; Liu, H.; Grilley, B. J.; Gee, A. P.; Rooney, C. M.; Heslop, H. E.; Brenner, M. K. Inducible Caspase-9 Suicide Gene Controls Adverse Effects from Alloreplete T Cells after Haploidentical Stem Cell Transplantation. Blood 2015, 125 (26), 4103–4113. 10.1182/blood-2015-02-628354.

(18) Straathof, K. C.; Pulè, M. A.; Yotnda, P.; Dotti, G.; Vanin, E. F.; Brenner, M. K.; Heslop, H. E.; Spencer, D. M.; Rooney, C. M. An Inducible Caspase 9 Safety Switch for T-Cell Therapy. Blood 2005, 105 (11), 4247–4254. 10.1182/blood-2004-11-4564.

(19) Adams, E. L.; McGovern, A. C.; So, V.; Srinivasan, S.; Deiters, A.; Lohmueller, J. Small-Molecule Control of CAR T Cells. Nat Rev Chem 2025, 9 (12), 809–825. 10.1038/s41570-025-00768-6.

(20) Di Stasi, A.; Tey, S.-K.; Dotti, G.; Fujita, Y.; Kennedy-Nasser, A.; Martinez, C.; Straathof, K.; Liu, E.; Durett, A. G.; Grilley, B.; Liu, H.; Cruz, C. R.; Savoldo, B.; Gee, A. P.; Schindler, J.; Krance, R. A.; Heslop, H. E.; Spencer, D. M.; Rooney, C. M.; Brenner, M. K. Inducible Apoptosis as a Safety Switch for Adoptive Cell Therapy. N Engl J Med 2011, 365 (18), 1673–1683. 10.1056/NEJMoa1106152.

(21) Liu, X.; Ciulli, A. Proximity-Based Modalities for Biology and Medicine. ACS Central Science 2023, 9 (7), 1269–1284. 10.1021/acscentsci.3c00395.

(22) Robinson, S. A.; Co, J. A.; Banik, S. M. Molecular Glues and Induced Proximity: An Evolution of Tools and Discovery. Cell Chemical Biology 2024, 31 (6), 1089–1100. 10.1016/j.chembiol.2024.04.001.

(23) Stanton, B. Z.; Chory, E. J.; Crabtree, G. R. Chemically Induced Proximity in Biology and Medicine. Science 2018, 359 (6380), eaao5902. 10.1126/science.aao5902.

(24) Bayle, J. H.; Grimley, J. S.; Stankunas, K.; Gestwicki, J. E.; Wandless, T. J.; Crabtree, G. R. Rapamycin Analogs with Differential Binding Specificity Permit Orthogonal Control of Protein Activity. Chemistry & Biology 2006, 13 (1), 99–107. 10.1016/j.chembiol.2005.10.017.

(25) Hathaway, N. A.; Bell, O.; Hodges, C.; Miller, E. L.; Neel, D. S.; Crabtree, G. R. Dynamics and Memory of Heterochromatin in Living Cells. Cell 2012, 149 (7), 1447–1460. 10.1016/j.cell.2012.03.052.

(26) Liang, F.-S.; Ho, W. Q.; Crabtree, G. R. Engineering the ABA Plant Stress Pathway for Regulation of Induced Proximity. Sci. Signal. 2011, 4 (164). 10.1126/scisignal.2001449.

(27) Miyamoto, T.; DeRose, R.; Suarez, A.; Ueno, T.; Chen, M.; Sun, T.; Wolfgang, M. J.; Mukherjee, C.; Meyers, D. J.; Inoue, T. Rapid and Orthogonal Logic Gating with a Gibberellin-Induced Dimerization System. Nat Chem Biol 2012, 8 (5), 465–470. 10.1038/nchembio.922.

(28) Fink, T.; Lonzarić, J.; Praznik, A.; Plaper, T.; Merljak, E.; Leben, K.; Jerala, N.; Lebar, T.; Strmšek, Ž.; Lapenta, F.; Benčina, M.; Jerala, R. Design of Fast Proteolysis-Based Signaling and Logic Circuits in Mammalian Cells. Nat Chem Biol 2019, 15 (2), 115–122. 10.1038/s41589-018-0181-6.

(29) Dolberg, T. B.; Meger, A. T.; Boucher, J. D.; Corcoran, W. K.; Schauer, E. E.; Prybutok, A. N.; Raman, S.; Leonard, J. N. Computation-Guided Optimization of Split Protein Systems. Nature Chemical Biology 2021, 17 (5), 531–539. 10.1038/s41589-020-00729-8.

(30) Benaissa, H.; Ounoughi, K.; Aujard, I.; Fischer, E.; Goïame, R.; Nguyen, J.; Tebo, A. G.; Li, C.; Le Saux, T.; Bertolin, G.; Tramier, M.; Danglot, L.; Pietrancosta, N.; Morin, X.; Jullien, L.; Gautier, A. Engineering of a Fluorescent Chemogenetic Reporter with Tunable Color for Advanced Live-Cell Imaging. Nature Communications 2021, 12 (1), 6989. 10.1038/s41467-021-27334-0.

(31) Dixon, A. S.; Schwinn, M. K.; Hall, M. P.; Zimmerman, K.; Otto, P.; Lubben, T. H.; Butler, B. L.; Binkowski, B. F.; MacHleidt, T.; Kirkland, T. A.; Wood, M. G.; Eggers, C. T.; Encell, L. P.; Wood, K. V. NanoLuc Complementation Reporter Optimized for Accurate Measurement of Protein Interactions in Cells. ACS Chemical Biology 2016, 11 (2), 400–408. 10.1021/acschembio.5b00753.

(32) Hall, M. P.; Unch, J.; Binkowski, B. F.; Valley, M. P.; Butler, B. L.; Wood, M. G.; Otto, P.; Zimmerman, K.; Vidugiris, G.; MacHleidt, T.; Robers, M. B.; Benink, H. A.; Eggers, C. T.; Slater, M. R.; Meisenheimer, P. L.; Klaubert, D. H.; Fan, F.; Encell, L. P.; Wood, K. V. Engineered Luciferase Reporter from a Deep Sea Shrimp Utilizing a Novel Imidazopyrazinone Substrate. ACS Chemical Biology 2012, 7 (11), 1848–1857. 10.1021/cb3002478.

(33) Chung, H. K.; Lin, M. Z. On the Cutting Edge: Protease-Based Methods for Sensing and Controlling Cell Biology. Nature Methods 2020, 17 (9), 885–896. 10.1038/s41592-020-0891-z.

(34) Wu, Y.; von Hauff, I.; Jensen, N.; Rossner, M.; Wehr, M. Improved Split TEV GPCR β-Arrestin-2 Recruitment Assays via Systematic Analysis of Signal Peptide and β-Arrestin Binding Motif Variants. Biosensors 2022, 13 (1), 48. 10.3390/bios13010048.

(35) Vlahos, A. E.; Kang, J.; Aldrete, C. A.; Zhu, R.; Chong, L. S.; Elowitz, M. B.; Gao, X. J. Protease-Controlled Secretion and Display of Intercellular Signals. Nature Communications 2022, 13 (1), 912. 10.1038/s41467-022-28623-y.

(36) Gao, X. J.; Chong, L. S.; Kim, M. S.; Elowitz, M. B. Programmable Protein Circuits in Living Cells. Science 2018, 361 (6408), 1252–1258. 10.1126/science.aat5062.

(37) Zhu, L.; McNamara, H. M.; Toettcher, J. E. Light-Switchable Transcription Factors Obtained by Direct Screening in Mammalian Cells. Nature Communications 2023, 14 (1). 10.1038/s41467-023-38993-6.

(38) Jullien, N.; Goddard, I.; Selmi-Ruby, S.; Fina, J.-L.; Cremer, H.; Herman, J.-P. Conditional Transgenesis Using Dimerizable Cre (DiCre). PLoS ONE 2007, 2 (12), e1355. 10.1371/journal.pone.0001355.

(39) Kawano, F.; Okazaki, R.; Yazawa, M.; Sato, M. A Photoactivatable Cre-loxP Recombination System for Optogenetic Genome Engineering. Nature Chemical Biology 2016, 12 (12), 1059–1064. 10.1038/nchembio.2205.

(40) Morikawa, K.; Furuhashi, K.; de Sena-Tomas, C.; Garcia-Garcia, A. L.; Bekdash, R.; Klein, A. D.; Gallerani, N.; Yamamoto, H. E.; Park, S.-H. E.; Collins, G. S.; Kawano, F.; Sato, M.; Lin, C.-S.; Targoff, K. L.; Au, E.; Salling, M. C.; Yazawa, M. Photoactivatable Cre Recombinase 3.0 for in Vivo Mouse Applications. Nature Communications 2020, 11 (1), 2141. 10.1038/s41467-020-16030-0.

(41) Ray, P.; Pimenta, H.; Paulmurugan, R.; Berger, F.; Phelps, M. E.; Iyer, M.; Gambhir, S. S. Noninvasive Quantitative Imaging of Protein–Protein Interactions in Living Subjects. Proceedings of the National Academy of Sciences 2002, 99 (5), 3105–3110. 10.1073/pnas.052710999.

(42) Luan, H.; Diao, F.; Scott, R. L.; White, B. H. The Drosophila Split Gal4 System for Neural Circuit Mapping. Frontiers in Neural Circuits 2020, 14 (November), 603397. 10.3389/fncir.2020.603397.

(43) Ma, D.; Yuan, Q.; Peng, F.; Paredes, V.; Zeng, H.; Osikpa, E. C.; Yang, Q.; Peddi, A.; Patel, A.; Liu, M. S.; Sun, Z.; Gao, X. Engineered PROTAC-CID Systems for Mammalian Inducible Gene Regulation. J. Am. Chem. Soc. 2023, 145 (3), 1593–1606. 10.1021/jacs.2c09129.

(44) Yamada, M.; Nagasaki, S. C.; Suzuki, Y.; Hirano, Y.; Imayoshi, I. Optimization of Light-Inducible Gal4/UAS Gene Expression System in Mammalian Cells. iScience 2020, 23 (9), 101506. 10.1016/j.isci.2020.101506.

(45) Hong, M.; Fitzgerald, M. X.; Harper, S.; Luo, C.; Speicher, D. W.; Marmorstein, R. Structural Basis for Dimerization in DNA Recognition by Gal4. Structure 2008, 16 (7), 1019–1026. 10.1016/j.str.2008.03.015.

(46) Hayden, M. S.; Ghosh, S. Shared Principles in NF-κB Signaling. Cell 2008, 132 (3), 344–362. 10.1016/j.cell.2008.01.020.

(47) Marin, V.; Cribioli, E.; Philip, B.; Tettamanti, S.; Pizzitola, I.; Biondi, A.; Biagi, E.; Pule, M. Comparison of Different Suicide-Gene Strategies for the Safety Improvement of Genetically Manipulated T Cells. Human Gene Therapy Methods 2012, 23 (6), 376–386. 10.1089/hgtb.2012.050.

(48) Okamoto, T.; Natsume, Y.; Yamanaka, H.; Fukuda, M.; Yao, R. A Protocol for Efficient CRISPR-Cas9-Mediated Knock-in in Colorectal Cancer Patient-Derived Organoids. STAR Protocols 2021, 2 (4), 100780. 10.1016/j.xpro.2021.100780.

(49) Burns, S. M.; Vetere, A.; Walpita, D.; Dančík, V.; Khodier, C.; Perez, J.; Clemons, P. A.; Wagner, B. K.; Altshuler, D. High-Throughput Luminescent Reporter of Insulin Secretion for Discovering Regulators of Pancreatic Beta-Cell Function. Cell Metabolism 2015, 21 (1), 126–137. 10.1016/j.cmet.2014.12.010.

(50) Marchand, A.; Buckley, S.; Schneuing, A.; Pacesa, M.; Elia, M.; Gainza, P.; Elizarova, E.; Neeser, R. M.; Lee, P.-W.; Reymond, L.; Miao, Y.; Scheller, L.; Georgeon, S.; Schmidt, J.; Schwaller, P.; Maerkl, S. J.; Bronstein, M.; Correia, B. E. Targeting Protein–Ligand Neosurfaces with a Generalizable Deep Learning Tool. Nature 2025, 639 (8054), 522–531. 10.1038/s41586-024-08435-4.

(51) Rakotoarison, L. M.; Tebo, A. G.; Böken, D.; Board, S.; El Hajji, L.; Gautier, A. Improving Split Reporters of Protein-Protein Interactions through Orthology-Based Protein Engineering. ACS Chemical Biology 2024, 19 (2), 428–441. 10.1021/acschembio.3c00631.

(52) De Boeck, J.; Verfaillie, C. Doxycycline Inducible Overexpression Systems: How to Induce Your Gene of Interest without Inducing Misinterpretations. Molecular Biology of the Cell 2021, 32 (17), 1517–1522. 10.1091/mbc.E21-04-0177.

(53) Adams, E. L.; McGovern, A. C.; So, V.; Srinivasan, S.; Deiters, A.; Lohmueller, J. Small-Molecule Control of CAR T Cells. Nature Reviews Chemistry 2025, 9 (12), 809–825. 10.1038/s41570-025-00768-6.

(54) Gibson, D. G.; Young, L.; Chuang, R.-Y.; Venter, J. C.; Hutchison, C. A.; Smith, H. O. Enzymatic Assembly of DNA Molecules up to Several Hundred Kilobases. Nature Methods 2009, 6 (5), 343–345. 10.1038/nmeth.1318.

(55) Li, C.; Plamont, M. A.; Sladitschek, H. L.; Rodrigues, V.; Aujard, I.; Neveu, P.; Le Saux, T.; Jullien, L.; Gautier, A. Dynamic Multicolor Protein Labeling in Living Cells. Chemical Science 2017, 8 (8), 5598–5605. 10.1039/c7sc01364g.

(56) Plamont, M. A.; Billon-Denis, E.; Maurin, S.; Gauron, C.; Pimenta, F. M.; Specht, C. G.; Shi, J.; Quérard, J.; Pan, B.; Rossignol, J.; Morellet, N.; Volovitch, M.; Lescop, E.; Chen, Y.; Triller, A.; Vriz, S.; Le Saux, T.; Jullien, L.; Gautier, A. Small Fluorescence-Activating and Absorption-Shifting Tag for Tunable Protein Imaging in Vivo. Proceedings of the National Academy of Sciences of the United States of America 2016, 113 (3), 497–502. 10.1073/pnas.1513094113.

