## Supporting Information for "Chemically responsive protein switches for the precise control of biological activities"

**This PDF file includes:**

Supplementary Figs 1-2

Supplementary Tables 1-4

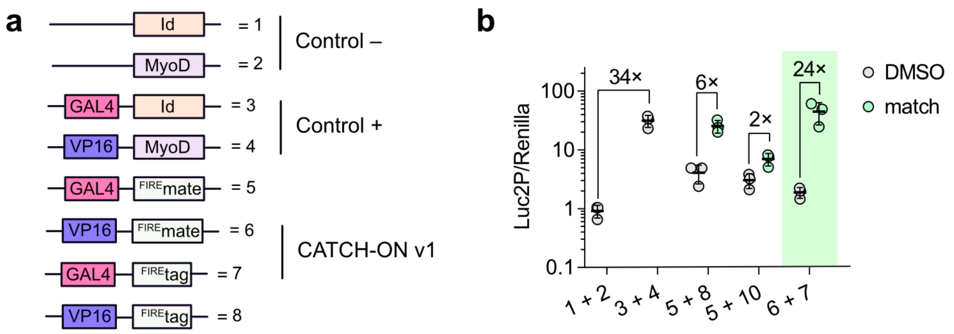


**Supplementary Figure 1. Chemically controlled gene expression – preliminary screening. a** Constructs used in this study. **b** HeLa cells were transfected and incubated with DMSO 0.5% or 5 µM of $match_{550}$ for 24 h. Results are depicted as a normalized signal of Luc2P *vs.* Renilla signal used as a transfection control. Values are shown as mean ± SD of n = 3 independent experiments (each composed of two technical replicates).

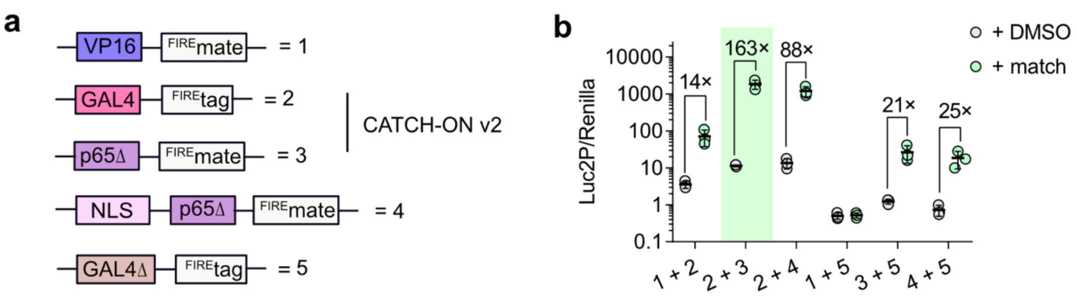


**Supplementary Figure 2. Chemically controlled gene expression with optimized CATCH-ON candidates. a** Constructs used in this study. **b** HeLa cells transfected with UAS-Luc2P plasmid and the CATCH-ON candidates were incubated with DMSO 0.5% or 5 µM of $match_{550}$ for 24 h. Results are depicted as a normalized signal of Luc2P *vs.* Renilla signal used as a transfection control. Values are shown as mean ± SD of n = 3 independent experiments (each composed of two technical replicates).

### Supplementary Tables

### Supplementary Table 1 Plasmid synthesis

| Plasmid number | Used in Fig. | ORF | Origin (Backbone/Insert) and Reference | Primers |
|---|---|---|---|---|
| pAG354 | 1,2,6 | CMV_Empty | Unpublished | |
| pAG1420 | 1 | CMV_Nanoluciferase_IRES_HA-iRFP670 | pAG1151[1]/ pAG1418 (Addgene 87696) | ag313,ag2350, ag2351, ag2352, ag2353, ag314 |
| pAG1473 | 1 | CMV_CMyc-FRB-LgBiT_IRES_HA-mTQ2 | pAG1142[1]/ Gblock | ag313, ag 2533, ag 2534, ag 2535, ag 2536, ag 314 |
| pAG1474 | 1 | CMV_FKBP-SmBiT_IRES_HA-iRFP670 | pAG1151/ Gblock | ag 313, ag 2537, ag 2538, ag 2539, ag 2540, ag 314 |
| pAG1494 | 1 | CMV_CMyc-LgBiT-FIREtag_IRES_HA-iRFP670 | pAG1142/ Gblock | ag313,ag2605, ag2606,ag2607, ag2608,ag314 |
| pAG1495 | 1 | CMV_CMyc-FIREmate-SmBiT_IRES_HA-mTQ2 | pAG1151/+ Gblock | ag313,ag2609, ag2610,ag2611, ag2612,ag314 |
| pAG1496 | 1 | CMV_CMyc-FIREtag-LgBiT_IRES_HA-iRFP670 | pAG1142/ Gblock | ag313,ag2632, ag2633,ag2634, ag2635,ag314 |
| pAG1497 | 1 | CMV_CMyc-SmBiT-FIREmate_IRES_HA-mTQ2 | pAG1151/ Gblock | ag313,ag2636, ag2637,ag2638, ag2639,ag314 |
| pAG1511 | 1 | CMV_FKBP-SmBiT-P2A-FRB-LgBiT_IRES_HA-iRFP-CMyc | pAG580[2]/ pAG1473 / pAG1474 | ag313,ag2688,ag2689,ag2690,ag2691,ag2692,ag2693,ag314 |
| pAG1512 | 1 | CMV_FIREmate-SmBiT-P2A-LgBiT-FIREtag_IRES_HA-iRFP-CMyc | pAG580/ pAG1494 / pAG1495 | ag3131,ag2694,ag2695,ag2696,ag2697,ag2698,ag2699 |
| pAG1605 | 2,4,5 | pGL4.31[luc2P/GAL4UAS/Hygro] | CheckMate™/Flexi® Vector Mammalian Two-Hybrid System PROMEGA C9360 | |
| pAG1606 | 4,S1 | pFN11 Act Vector | CheckMate™/Flexi® Vector Mammalian Two-Hybrid System PROMEGA C9360 | |
| pAG1607 | 4,S1 | pFN10 BIND FlexiVector | CheckMate™/Flexi® Vector Mammalian Two-Hybrid System PROMEGA C9360 | |
| pAG1608 | 4,S1 | pBIND-Id Control Vector | CheckMate™/Flexi® Vector Mammalian Two-Hybrid System PROMEGA C9360 | |
| pAG1609 | 4,S1 | pACT-MyoD Control Vector | CheckMate™/Flexi® Vector Mammalian Two-Hybrid System PROMEGA C9360 | |
| pAG1611 | 4,S1 | CMV_GAL4-FIREmate | pFN11 BIND Flexi vector, PROMEGA/ pAG1512 | ag2913,ag2914,ag2915,ag2916,ag2917,ag2918 |
| pAG1612 | 4,5,6,S2,S1,S1 | CMV_GAL4-FIREtag | pFN11 BIND Flexi vector, PROMEGA | ag2913,ag2919,ag2920,ag2918 |

| | | | | |
|---|---|---|---|---|
| pAG1614 | 4, 5, S1, S2 | CMV_VP16-FIREmate | pFN10 ACT Flexivector PROMEGA/ pAG1512 | ag2913,ag2914,ag2915,ag2916,ag2917,ag2918 |
| pAG1615B | 4, S1 | CMV_VP16-FIREtag | pFN10 ACT Flexivector PROMEGA | ag2913,ag2919,ag2920,ag2918 |
| pAG1673 | 2, 5 | CMV_Full TEVp | Addgene 64276 | |
| pAG1674 | 2 | CMV_cMyc-FRB-nSuMMVp | Addgene 118970 | |
| pAG1675 | 2 | CMV_FKBP-cSuMMVp-HA | Addgene 119218 | |
| pAG1676 | 2 | CMV_Cyc_Fluc_TEV | Addgene 119207 | |
| pAG1677 | 2 | CMV_Cyc_Fluc_PPV | Addgene 119208 | |
| pAG1678 | 2 | CMV_Cyc_Fluc_SuMMVp | Addgene 119210 | |
| pAG1679 | 2 | CMV_cMyc-PYL1_nPPVp | Addgene 119211 | |
| pAG1680 | 2 | CMV_cMyc-ABI_cPPVp | Addgene 119212 | |
| pAG1681 | 2 | CMV_HA-PYL1_NTEVp | Addgene 119213 | |
| pAG1682 | 2 | CMV_cMyc-ABI_CTEVp | Addgene 119214 | |
| pAG1683 | 2 | CMV_DRD1-NTEVp-cMyc-TEVcs-GAL4-VP16-HA-HA | Addgebe 194358 | |
| pAG1684 | 2 | CMV_b-Arrestin-2-CTEVp-HA-HA | Addgene 194382 | |
| pAG1687 | 2 | CMV_AU1-FIREtag-NTEVp | Addgene 119213 | ag2918,ag3003,ag3004,ag313,ag314,ag2913 |
| pAG1688 | 2 | CMV_AU1-FIREmate-NTEVp | Addgene 119214 (pAG1682)/ pAG1512 | ag2918,ag3005,ag3006,ag3007,ag3008,ag314,ag313,ag2913 |
| pAG1689 | 2 | CMV_CMyc-FIREtag-CTEVp | Addgene 119213 (pAG1681) | ag2918,ag3009,ag3010,ag314,ag313,ag314,ag2913 |
| pAG1690 | 2 | CMV_CMyc_FIREmate-CTEVp | Addgene 119214 (pAG1682)/ pAG1512 | ag2918,ag3011,ag3012,ag3013,ag3014,ag314,ag313,ag2913 |
| pAG1691 | 2 | CMV_CMyc-Renilla-IRES-HA-iRFP670 | pAG580/Addgene 101139 (pAG1681) | ag313,ag3027,ag3028,ag3029,ag3030,ag314 |
| pAG1735 | 2 | CMV_DRD1-FIREtag-NTEVp-TEVcs-CMyc-GAL4-SV40NLS-VP16AD-2xHA | Addgene 194358 (pAG1683)/pAG1687 | ag2918,ag3102,ag3103,ag3104,ag3105,ag314,ag3113,ag2913 |
| pAG1738 | 2,4,5 | 10xUAS-MLPmin-d2EGFP-hPEST PGK iRFP713 | Addgene 194383 (pAG1685) / Addgene 203908 (pAG1686) | ag2905,ag3114,ag3115,ag3116,ag3117,ag2912 |
| pAG1803 | 3, 5 | CMV_Cre_Recombinase | Addgene 11916 (pAG1803) | |
| pAG1807 | 3 | CMV_loxP_STOP_loxP_Fluc | Addgene 122962 (pAG1807) | |
| pAG1808 | 3 | CMV_loxP_STOP_mCherry | Addgene 122963 (pAG1808) | |
| pAG2013 | 6 | UAS_PeptideB-FCS-NanoLuc-FCS | pAG1812/ Addgene 179633 (pAG1742) | ag2905,ag4074,ag4075,ag2912 |

| | | | | |
|---|---|---|---|---|
| | | -PeptideA-FCS-3XTM-TEVpCS-AAMP | + Addgene 62057 (pAG1741) | |
| pAG2014 | 6 | CMV_PeptideB-FCS-NanoLuc-FCS -PeptideA-FCS-3XTM-TEVpCS-AAMP | pAG1813/ Addgene 179633 (pAG1742) + Addgene 62057 (pAG1741) | ag2905,ag4074,ag4075,ag2912 |
| pAG1819 | 3 | CMV_CreN59-FIREmate-NLS_P2A_NLS_FIREtag-CreC60 | pAG1817 /122960 Addgene (pAG1806) | ag2918,ag3396,ag3397,ag314,ag313,ag2913 |
| pAG1867 | 5, 6, S2 | CMV_p65Δ-FIREmate | pAG1614/pAG1244 | ag2913,ag3605,ag3606,ag3607,ag3608,ah2918 |
| pAG1868 | 5, S2 | CMV_GAL4Δ-FIREtag | pAG1612 | ag2913,ag3609,ag3610,ag2918 |
| pAG1869 | S2 | CMV_NLS-p65Δ-FIREmate | pAG1614/pAG1244 | ag2913,ag3611,ag3612,ag3607,ag3608,ag2918 |
| pAG1870 | 5 | pGL4.31[EmGFP-P2A-DmrB-iCasp9-hPEST_GAL4UAS_Hygro] | pAG1605/Addgene 178533 (pAG1400) | ag2908,ag3613,ag3614,ag3615,ag3616,ag2910,ag2911,ag2912 |
| pAG1930 | 5 | pGL4.31[EmGFP-P2A-TEV-hPEST_GAL4UAS_Hygro] | pAG1870 / Addgene 62537 (pAG1673) | ag2905,ag3746,ag3747,ag3748,ag3749,ag2910,ag2911,ag2912 |
| pAG1931 | 5 | pGL4.31[EmGFP-P2A-CRE-hPEST_GAL4UAS_Hygro] | pAG1870/ Addgene 119616 (pAG1803) | ag2905,ag3750,ag3751,ag3752,ag3753,ag2910,ag2911,ag2912 |
| pAG1672 | 6 | CMV_Furin_6xHis | Addgene 57155 | |
| pAG1052 | 3 | CMV_FRB-EGFP | pAG1052[3] | |

### Supplementary Table 2 Primers sequence

| Primer | Sequence 5' to 3' |
|---|---|
| ag313 | ctcaccttgctcctgccgagaaagtatcca |
| ag314 | tggatactttctcggcaggagcaaggtgag |
| ag2688 | CCTGCACTCCCATggtggcagatctgagtccgg |
| ag2689 | actcagatctgccaccATGGGAGTGCAGGTGGAAACC |
| ag2690 | TCCACGTCTCCAGCCTGCTTCAGCAGGCTGAAGTTAGTAGCTCCGCTTCCGAGTATCTCTTCAAACAACCTGTAACC |
| ag2691 | AGCCTGCTGAAGCAGGCTGGAGACGTGGAGGAGAACCCTGGACCTGAGATGTGGCATGAAGGCCTG |
| ag2692 | ggggc ggatCCCGGGTTAGGAATTAATTGTTACGCGGAAGAGCATAG |
| ag2693 | CGTAACAATTAATTCCTAACCCGGGatccgcccctc |
| ag2694 | CTGACCCGAATGCGACATGCTCCATggtggcagatctgagtccgg |
| ag2695 | actcagatctgccaccATGGAGCATGTCGCATTCGG |
| ag2696 | TCCACGTCTCCAGCCTGCTTCAGCAGGCTGAAGTTAGTAGCTCCGCTTCC GAGAATTTCCTCAAACAAGCGATAC |
| ag2697 | CAGCCTGCTGAAGCAGGCTGGAGACGTGGAGGAGAACCCTGGACCT ATGGTGTTTACATTGGAGGATTTTG |
| ag2698 | ggggc ggatCCCGGGTTAAACGCGCTTAACAAATACCC |
| ag2699 | AAGCGCGTT TAACCCGGGatccgcccctc |
| ag2700 | GACATGCTCCATCAAGTCCTCTTCAGAAATAAGCTTTTGTTCCATgg |
| ag2701 | GCTTATTTCTGAAGAGGACTTGATGGAGCATGTCGCGTTCGGAAG |
| ag2702 | cgCATggttGTGGccatattatcatcgtg |
| ag2703 | cacgatgataatatggCCACaaccATGcg |
| ag2704 | CTCATAATCATCAAGTCCTCTTCAGAAATAAGCTTTTGTTCCATgg |
| ag2705 | TTTCTGAAGAGGACTTGATGATTATGAGCGGTTATGTCAATAACCCCG |
| ag2706 | AATATCTATCGCCCAAGTCCTCTTCAGAAATAAGCTTTTGTTCCATgg |
| ag2707 | CTGAAGAGGACTTGGGCGATAGATATTGGGTTTTCGTGAAACG |
| ag2708 | TTAGCGTCCTCCATCAAGTCCTCTTCAGAAATAAGCTTTTGTTCCATgg |
| ag2709 | TTATTTCTGAAGAGGACTTGATGGAGGACGCTAAGAATATCAAGAAAGG |
| ag2905 | CCAGTGCTTGATCAGTGAGGCACCGATC |
| ag2906 | tcctcgcccttgctcaccatGGTGGCTTTACCAACAGTACCGGATTG |
| ag2907 | GTACTGTTGGTAAAGCCACCatggtgagcaagggcgaggagg |
| ag2908 | GGGAGGGAAGCCGTGAGAATTCAAGTCCTCTTCAGAAATAAGCTTTTGTTCGGATC |
| ag2909 | CAAAAGCTTATTTCTGAAGAGGACTTGAATTCTCACGGCTTCCCTCCCGAGG |
| ag2910 | GAAGATGTTGGCCACCTCGTACTGACTG |
| ag2911 | CAGTCAGTACGAGGTGGCCAACATCTTC |
| ag2912 | GATCGGTGCCTCACTGATCAAGCACTGG |
| ag2913 | CTTTTTTGCACAACATGGGGGATCATGTAACTC |
| ag2914 | GAATGCGACATGCTCCATTTCCATGGCGATCGCTGGAGAGG |
| ag2915 | CCAGCGATCGCCATGGAAATGGAGCATGTCGCATTCGGGTC |
| ag2916 | CGAATTCGTTTAAACATTAAGACAGTGCCTTCTTGAGATGCAC |
| ag2917 | TCAAGAAGGCACTGTCTTAATGTTTAAACGAATTCGGGCTCGGTACC |
| ag2918 | GAGTTACATGATCCCCCATGTTGTGCAAAAAAG |
| ag2919 | AACGCGCTTAACAAATACCCAATATCTGTCTCCTTCCATGGCGATCGCTGGAGAGG |
| ag2920 | ACAGATATTGGGTATTTGTTAAGCGCGTTTAATGTTTAAACGAATTCGGGCTCGGTACC |
| ag2921 | ATCGCCATGGAAGGAGACAGATATTGGGTATTTGTTAAGCGCG |

| Primer | Sequence 5' to 3' |
|---|---|
| ag2922 | GAATTCGTTTAAACATTAGGATAACGCTTTCTTCATGTGGACTTTC |
| ag2923 | ATGAAGAAAGCGTTATCCTAATGTTTAAACGAATTCGGGCTCGGTACC |
| ag3003 | AACGCGCTTAACAAATACCCAATATCTGTCTCCgatgtacctgtaggtgtccatggtgg |
| ag3004 | CAGATATTGGGTATTTGTTAAGCGCGTTggatccggaagtggagaaagcttgtttaag |
| ag3005 | CCGAATGCGACATGCTCCATgatgtacctgtaggtgtccatggtgg |
| ag3006 | catggacacctacaggtacatcATGGAGCATGTCGCATTCGGGTC |
| ag3007 | agctttctccacttccggatccAGACAGTGCCTTCTTGAGATGCAC |
| ag3008 | GTGCATCTCAAGAAGGCACTGTCTggatccggaagtggagaaagcttgtttaag |
| ag3009 | AACGCGCTTAACAAATACCCAATATCTGTCTCCcagatcctcctcgctgatcagcttc |
| ag3010 | GATATTGGGTATTTGTTAAGCGCGTTggatccggctccggcagcaag |
| ag3011 | TTCTGACCCGAATGCGACATGCTCCATcagatcctcctcgctgatcagcttc |
| ag3012 | ggagcagaagctgatcagcgaggaggatctgATGGAGCATGTCGCATTCGGGTC |
| ag3013 | acatgctcttgctgccggagccggatccAGACAGTGCCTTCTTGAGATGCACTTTGAC |
| ag3014 | GTGCATCTCAAGAAGGCACTGTCTggatccggctccggcagcaag |
| ag3027 | tttcgaagtcatgaattcCAAGTCCTCTTCAGAAATAAGCTTTTGTTCCATgg |
| ag3028 | GCTTATTTCTGAAGAGGACTTGgaattcatgacttcgaaagtttatgatccagaac |
| ag3029 | CTTGTCGTCGTCGTCCTTGTAGTCCTCGAGTTAtcattgttcatttttgagaactcgc |
| ag3030 | gagttctcaaaaatgaacaatgaTAACTCGAGGACTACAAGGACGACG |
| ag3102 | CTTAACAAATACCCAATATCTGTCTCCGCTACCTCCACCTCCGCTACCTC |
| ag3103 | ggaggtggaggtagcGGAGACAGATATTGGGTATTTGTTAAGCGC |
| ag3104 | accctggaagtacaggttctcagtttggaagttggttgtcacaag |
| ag3105 | gacaaccaacttccaaactgagaacctgtacttccagggttctagagaac |
| ag3106 | GCGACATGCTCCATcagatcctcctcgctgatcagcttctgctc |
| ag3107 | tgatcagcgaggaggatctgATGGAGCATGTCGCATTCGGGTC |
| ag3108 | gaacatcgtatgggtattgcgagtacaccaattcactcatgag |
| ag3109 | gaattggtgtactcgcaatacccatacgatgttcctgactatgc |
| ag3110 | ttgggagatctCCTCATcatGGCCCGGGCCAGATCCTC |
| ag3111 | cagaagaggatctggcccgggccatgATGAGGagatctcccaagaagagg |
| ag3112 | aacatcgtatgggtagcccCTTGTACAGCTCGTCCATGCCGC |
| ag3113 | GCATGGACGAGCTGTACAAGggctacccatacgatgttcctgactatg |
| ag3114 | ctcgcccttgctcaccatGGTGGCTTTACCAACAGTACCGGATTG |
| ag3115 | gtactgttggtaaagccaccATGGTGAGCAAGGGCGAGGAG |
| ag3116 | gccccgactctagaattaTTACTCTTCCATCACGCCGATCTGC |
| ag3117 | gcgtgatggaagagtaaTAATTCTAGAGTCGGGGCGGCCG |
| ag3118 | GAACGCATTCTGGCGtaattctagagtcggggcggccgg |
| ag3119 | TGTGAAGACCATggtggctttaccaacagtaccggattgc |
| ag3120 | actgttggtaaagccaccATGGTCTTCACACTCGAAGATTTCGTTGG |
| ag3121 | ggccgccccgactctagaaTTACGCCAGAATGCGTTCGCACAGC |
| ag3201 | atgtacctgtaggtgtcCATggtggcagatctgagtccgg |
| ag3202 | accggactcagatctgccaccATGgacacctacaggtacatcGGAGAC |
| ag3203 | TCCACGTCTCCAGCCTGCTTCAGCAGGCTGAAGTTAGTAGCTCCGCTTCCagtttggaagttggttgtcacaaga |
| ag3204 | GCCTGCTGAAGCAGGCTGGAGACGTGGAGGAGAACCCTGGACCTATGGAGCATGTCGCATTCGGGTC |
| ag3205 | ggcggatCCCGGGttattgcgagtacaccaattcactcatgagttg |

| Primer | Sequence 5' to 3' |
|---|---|
| ag3206 | ctcatgagtgaattggtgtactcgcaaTAACCCGGGatccgcccctctc |
| ag3207 | CTGACCCGAATGCGACATGCTCCATggtggcagatctgagtccgg |
| ag3208 | ggactcagatctgccaccATGGAGCATGTCGCATTCGGGTC |
| ag3209 | GTCTCCAGCCTGCTTCAGCAGGCTGAAGTTAGTAGCTCCGCTTCCttgcgagtacaccaattcactcatgag |
| ag3210 | GCCTGCTGAAGCAGGCTGGAGACGTGGAGGAGAACCCTGGACCTgacacctacaggtacatcGGAGAC |
| ag3211 | gaggggcggatCCCGGGTTAagtttggaagttggttgtcacaag |
| ag3212 | gacaaccaacttccaaactTAACCCGGGatccgcccctctc |
| ag2533 | cactgcctccgacctgctttgagattcgtcggaac |
| ag2534 | aaagcaggtcggaggcagtggaggcgg |
| ag2535 | ctcgaggttaggaattaattgttacgcggaagagcat |
| ag2536 | aattaattcctaacctcgaggactacaaggacgacg |
| ag2537 | CCTGCACTCCCATGGTGGCAGATCTGAGTCCG |
| ag2538 | tgccaccatgggagtgcaggtggaaaccat |
| ag2539 | cggaaccaccttcttccagttttagaagctccac |
| ag2540 | actggaagaaggtggttccggaggcggt |
| ag2605 | ATGTAAACACCATgaattcCAAGTCCTCTTCAGAAATAAGCTTTTGTTCC |
| ag2606 | GCTTATTTCTGAAGAGGACTTGgaattcATGGTGTTTACATTGGAGGATTTTG |
| ag2607 | TCCTTGTAGTCCTCGAGTTAAACGCGCTTAACAAATACCC |
| ag2608 | ATTTGTTAAGCGCGTTTAACTCGAGGACTACAAGGACGACG |
| ag2609 | ACATGCTCCATgaattcCAAGTCCTCTTCAGAAATAAGCTTTTGTTCC |
| ag2610 | TATTTCTGAAGAGGACTTGgaattcATGGAGCATGTCGCATTCGG |
| ag2611 | TTGTAGTCCTCGAGGTTAGAGAATTTCCTCAAACAAGCGATACC |
| ag2612 | GGAAATTCTCTAACCTCGAGGACTACAAGGACGACG |
| ag2632 | AGCGATCTCCGAATTCCAAGTCCTCTTCAGAAATAAGCTTTTGTTCC |
| ag2633 | GAAGAGGACTTGgaattc GGAGATCGCTACTGGGTCTTC |
| ag2634 | GTCCTCGAGTTA ACTGTTTATCGTGACGCGAA |
| ag2635 | GTCACGATAAACAGTTAACTCGAGGACTACAAGGACGACG |
| ag2636 | GTAGCCGGTTACGAATTCCAAGTCCTCTTCAGAAATAAGCTTTTGTTCC |
| ag2637 | GGACTTGgaattcGTAACCGGCTACCGACTGTT |
| ag2638 | TCCTCGAGGTTAAGAGAGTGCTTTTTTAAGGTGC |
| ag2639 | AGCACTCTCTTAACCTCGAGGACTACAAGGACGACG |
| ag3304 | cgcatccacagggccatGGTGGCTTTACCAACAGTACCGGATTG |
| ag3305 | ctgttggtaaagccaccatggccctgtggatg |
| ag3306 | gaattactaaggcatggcggcttcatcaataaaac |
| ag3307 | tgatgaagccgccatgccttagTAATTCTAGAGTCGGGGCGGCCG |
| ag3257 | gttcgggccctacagatatttcttttcatcaataaaactgcgtctgC |
| ag3258 | tattgatgaaaagaaatatctgtagggcccgaacaaaaactcatctc |
| ag3253 | gaagcgcatccacagggccatggtggcgaagcttaagtttaaacgcta |
| ag3254 | aagcttcgccaccATGGCCCTGTGGATGCGCTTC |
| ag3255 | ttttgcttctgccagaggatccacccgggtgcgcggcgttgcagtagttctccagctgg |
| ag3256 | CGGGTGGATCCTCTGGCAGAAGC |
| ag3257 | gttcgggccctacagatatttcttttcatcaataaaactgcgtctgC |
| ag3258 | tattgatgaaaagaaatatctgtagggcccgaacaaaaactcatctc |

| Primer | Sequence 5' to 3' |
|---|---|
| ag3336 | gcattctggcgCGTCAAaagcgtggcattgtagatcagtgctg |
| ag3337 | gcccagcagCGTACAaagagggtcttcacactcgaagatttc |
| ag3338 | caatgccacgcttTTGACGCGCCAGAATGCGTTCGCACAG |
| ag3339 | gcattctggcgCGTCAAaagcgtggcattgtagatcagtgctg |
| ag3340 | gtgaagaccctcttTGTACGctgctgggccacctccagtgc |
| ag3341 | gcccagcagCGTACAaagagggtcttcacactcgaagatttc |
| ag3342 | caatgccacgcttTTGACGCGCCAGAATGCGTTCGCACAG |
| ag3343 | gcattctggcgCGTCAAaagcgtggcattgtagatcagtgctg |
| ag3381 | AGTGTGAAGACCATggcggcgaattcataacttcgtataatg |
| ag3382 | tatgaattcgccgccATGGTCTTCACACTCGAAGATTTCG |
| ag3383 | gccctctagactcgagTTACGCCAGAATGCGTTCG |
| ag3384 | GCGAACGCATTCTGGCGTAActcgagtctagagggcccgtttaaac |
| ag3396 | AACGCGCTTAACAAATACCCAATATCTGTCTCCtccgccgactttcctcttcttcttg |
| ag3397 | GACAGATATTGGGTATTTGTTAAGCGCGTTggtaccaacaggaaatggttccctg |
| ag3398 | GAAAGCGACATGCTCCATTCCGCCGACTTTCCTCTTCTTCTTG |
| ag3399 | aggaaagtcggcggaATGGAGCATGTCGCTTTCGGATCAG |
| ag3400 | tcctgttggtaccAGAGAGTGCTTTTTTAAGGTGCACTTTAAC |
| ag3401 | AAAGCACTCTCTggtaccaacaggaaatggttccctgc |
| ag3402 | AACGCGCTTAACAAATACCCAATATCTGTCTCCggtaccgttcagcttgcaccaggc |
| ag3403 | AGATATTGGGTATTTGTTAAGCGCGTTggcggaagcggtggcgtgc |
| ag3549 | ATCTGTCTCCtccgcccacctcgagggggccggggttctc |
| ag3550 | aggagaaccccggccccctcgaggtgggcggaGGAGACAGATATTGGGTATTTG |
| ag3551 | cccgatatcaagacctgctaatttcaaggctaacacctcgagggggccgggg |
| ag3552 | ttgaaattagcaggtcttgatatcgggagcggcggaGGAGACAGATATTGGGTATTTG |
| ag3396 | AACGCGCTTAACAAATACCCAATATCTGTCTCCtccgccgactttcctcttcttcttg |
| ag3397 | GACAGATATTGGGTATTTGTTAAGCGCGTTggtaccaacaggaaatggttccctg |
| ag3401 | AAAGCACTCTCTggtaccaacaggaaatggttccctgc |
| ag3402 | AACGCGCTTAACAAATACCCAATATCTGTCTCCggtaccgttcagcttgcaccaggc |
| ag3403 | AGATATTGGGTATTTGTTAAGCGCGTTggcggaagcggtggcgtgc |
| ag3549 | ATCTGTCTCCtccgcccacctcgagggggccggggttctc |
| ag3550 | aggagaaccccggccccctcgaggtgggcggaGGAGACAGATATTGGGTATTTG |
| ag3551 | cccgatatcaagacctgctaatttcaaggctaacacctcgagggggccgggg |
| ag3552 | ttgaaattagcaggtcttgatatcgggagcggcggaGGAGACAGATATTGGGTATTTG |
| ag3605 | tatctggcaggtactgCATCTTTCAGGAGGCTTGCTTCAAG |
| ag3606 | GCAAGCCTCCTGAAAGATGcagtacctgccagatacagacgatc |
| ag3607 | GTCGACGGAATCGCTGGAGAggagctgatctgactcagcaggg |
| ag3608 | cctgctgagtcagatcagctccTCTCCAGCGATTCCGTCGACACC |
| ag3609 | GGTGGTGTCGACGGAATCGCTTCCAGTCTTTCTAGCCTTGATTCCAC |
| ag3610 | GGAATCAAGGCTAGAAAGACTGGAAGCGATTCCGTCGACACCACCTAC |
| ag3611 | ctggactttcctcttcttcttgggCATCTTTCAGGAGGCTTGCTTCAAG |
| ag3175 | gctgcaataaacaagttggggtgg |
| ag3176 | ccaccccaacttgtttattgcagc |
| ag3340 | gtgaagaccctcttTGTACGctgctgggccacctccagtgc |

| **Primer** | **Sequence 5' to 3'** |
|---|---|
| ag3341 | gcccagcagCGTACAaagagggtcttcacactcgaagatttc |
| ag3342 | caatgccacgcttTTGACGCGCCAGAATGCGTTCGCACAG |
| ag3343 | gcattctggcgCGTCAAaagcgtggcattgtagatcagtgctg |
| ag3613 | agctcctcgcccttgctcaccatGGTGGCTTTACCAACAGTACCGGATTG |
| ag3614 | CGGTACTGTTGGTAAAGCCACCatggtgagcaagggcgaggagc |
| ag3615 | CGGGAGGGAAGCCGTGAGAATTTgcgtagtctggtacgtcgtacgg |
| ag3616 | gtacgacgtaccagactacgcaAATTCTCACGGCTTCCCTCCCGAG |
| ag3746 | gaaaaggctctcgcccatctcgagcatggtaggtccaggg |
| ag3747 | cctggacctaccatgctcgagatgggcgagagccttttcaagg |
| ag3748 | tagtctggtacgtcgtacggataaagctgagtggcttccttaacgg |
| ag3749 | ggaagccactcagctttatccgtacgacgtaccagactacgc |
| ag3750 | tacggtcagtaaattggacat ctcgagcatggtaggtccaggg |
| ag3751 | gacctaccatgctcgagatgtccaatttactgaccgtacacc |
| ag3752 | gtacgtcgtacggataatcgccatcttccagcaggc |
| ag3753 | cctgctggaagatggcgattatccgtacgacgtaccagactacgc |
| ag4074 | gtgtgaagaccctcttTGTACGggacatgggtgtgtagaagaagcc |
| ag4075 | cttcttctacacacccatgtccCGTACAaagagggtcttcacactcgaag |

**Supplementary Table 3 Western Blot DNA mixtures**

| Figure | Plasmid | Protein | Amount (ng) |
|---|---|---|---|
| 2 | pAG1605 | Luc2P | 500 |
| | pAG1690 | FIREmate-CTEVp | 250 |
| | pAG1735 | FIREtag-NTEVp-CS-GAL4-VP16 | 250 |
| 3 | pAG1605 | Luc2P | 1000 |
| | pAG1803 | CRE | 5 |
| | pAG354 | Empty plasmid | 5 |
| | pAG1819 | Split CRE | 5 |
| 4 | pAG1605 | Luc2P | 500 |
| | pAG1606 | BIND | 250 |
| | pAG1607 | ACT | 250 |
| | pAG1608 | BIND-Id | 250 |
| | pAG1609 | ACT_MyoD | 250 |
| | pAG1612 | GAL4-FIREtag | 250 |
| | pAG1614 | VP16-FIREmate | 250 |
| 5 | pAG1605 | Luc2P | 500 |
| | pAG1612 | GAL4-FIREtag | 250 |
| | pAG1867 | p65Δ-FIREmate | 250 |

**Supplementary Table 4. Microscopy DNA mixtures**

| Figure | Plasmid | Protein | Amount (ng) |
|---|---|---|---|
| 2 | pAG1738 | UAS d2EGFP PGK iRFP713 | 400 |
| | pAG1690 | FIREmate-CTEVp | 300 |
| | pAG1735 | FIREtag-NTEVp-CS-GAL4-VP16 | 300 |
| 3 | pAG1808 | CMV_STOP_mCherry | 800 |
| | pAG1052 | CMV_FRB-EGFP | 200 |
| | pAG354 | Ghost plasmid | 40 |
| | pAG1803 | CMV_CRE | 40 |
| | pAG1819 | Split CRE | 40 |
| 4 | pAG1738 | UAS d2EGFP PGK iRFP713 | 400 |
| | pAG1606 | pACT | 300 |
| | pAG1607 | pACT-MyoD | 300 |
| | pAG1608 | pBIND | 300 |
| | pAG1609 | pBIND-Id | 300 |
| | pAG1612 | GAL4-FIREtag | 300 |
| | pAG1614 | VP16-FIREmate | 300 |
| 5 | pAG1738 | UAS d2EGFP PGK iRFP713 | 400 |
| | pAG1612 | GAL-FIREtag | 300 |
| | pAG1614 | VP16-FIREmate | 300 |
| | pAG1867 | p65Δ-FIREmate | 300 |

**ORF's**

pAG1420 CMV_NanoLuc_IRES_HA-iRFP-CMyc

ATGGTCTTCACACTCGAAGATTTCGTTGGGGACTGGCGACAGACAGCCGGCTACAACC
TGGACCAAGTCCTTGAACAGGGAGGTGTGTCCAGTTTGTTTCAGAATCTCGGGGTGTCC
GTAACTCCGATCCAAAGGATTGTCCTGAGCGGTGAAAATGGGCTGAAGATCGACATCCA
TGTCATCATCCCGTATGAAGGTCTGAGCGGCGACCAAATGGGCCAGATCGAAAAAATTT
TTAAGGTGGTGTACCCTGTGGATGATCATCACTTTAAGGTGATCCTGCACTATGGCACA
CTGGTAATCGACGGGGTTACGCCGAACATGATCGACTATTTCGGACGGCCGTATGAAG
GCATCGCCGTGTTCGACGGCAAAAAGATCACTGTAACAGGGACCCTGTGGAACGGCAA
CAAAATTATCGACGAGCGCCTGATCAACCCCGACGGCTCCCTGCTGTTCCGAGTAACCA
TCAACGGAGTGACCGGCTGGCGGCTGTGCGAACGCATTCTGGCG
TAACTCGAGGACTACAAGGACGACGACGACAAGCCCGGGatccgcccctctccctcccccccccta
acgttactggccgaagccgcttggaataaggccggtgtgcgtttgtctatatgttattttccaccatattgccgtcttttggcaatgtgag
ggcccggaaacctggccctgtcttcttgacgagcattcctaggggtctttcccctctcgccaaaggaatgcaaggtctgttgaatgtc
gtgaaggaagcagttcctctggaagcttcttgaagacaaacaacgtctgtagcgaccctttgcaggcagcggaaccccccacct
ggcgacaggtgcctctgcggccaaaagccacgtgtataagatacacctgcaaaggcggcacaaccccagtgccacgttgtga
gttggatagttgtggaaagagtcaaatggctctcctcaagcgtattcaacaaggggctgaaggatgcccagaaggtaccccattg
tatgggatctgatctggggcctcggtgcacatgctttacatgtgtttagtcgaggttaaaaaaacgtctaggccccccgaaccacgg
ggacgtggttttcctttgaaaaacacgatgataatatggCCACaaccATGcgatcgTACCCATACGATGTTCCAG
ATTACGCTGaattcatggcgcgtaaggtcgatctcacctcctgcgatcgcgagccgatccacatccccggcagcattcagc
cgtgcggctgcctgctagcctgcgacgcgcaggcggtgcggatcacgcgcattacggaaaatgccggcgcgttctttggacgcg
aaactccgcgggtcggtgagctactcgccgattacttcggcgagaccgaagcccatgcgctgcgcaacgcactggcgcagtctt
ccgatccaaagcgaccggcgctgatcttcggttggcgcgacggcctgaccggccgcaccttcgacatctcactgcatcgccatg
acggtacatcgatcatcgagttcgagcctgcggcggccgaacaggccgacaatccgctgcggctgacgcggcagatcatcgc
gcgcaccaaagaactgaagtcgctcgaagagatggccgcacgggtgccgcgctatctgcaggcgatgctcggctatcaccgc
gtgatgttgtaccgcttcgcggacgacggctccgggatggtgatcggcgaggcgaagcgcagcgacctcgagagctttctcggt
cagcactttccggcgtcgctggtcccgcagcaggcgcggctactgtacttgaagaacgcgatccgcgtggtctcggattcgcgcg
gcatcagcagccggatcgtgcccgagcacgacgcctccggcgccgcgctcgatctgtcgttcgcgcacctgcgcagcatctcgc
cctgccatctcgaatttctgcggaacatgggcgtcagcgcctcgatgtcgctgtcgatcatcattgacggcacgctatggggattgat
catctgtcatcattacgagccgcgtgccgtgccgatggcgcagcgcgtcgcggccgaaatgttcgccgacttcttatcgctgcactt
caccgccgcccaccaccaacgcGAACAAAAGCTTATTTCTGAAGAGGACTTGTAA

pAG1473_ CMyc-FRB-LgBiT _IRES_HA-mTQ2

ATGGAACAAAAGCTTATTTCTGAAGAGGACTTG gaattc
GAGATGTGGCATGAAGGCCTGGAAGAGGCATCTCGTTTGTACTTTGGGGAAAGGAACG
TGAAAGGCATGTTTGAGGTGCTGGAGCCCTTGCATGCTATGATGGAACGGGGCCCCCA
GACTCTGAAGGAAACATCCTTTAATCAGGCCTATGGTCGAGATTTAATGGAGGCCCAAG
AGTGGTGCAGGAAGTACATGAAATCAGGGAATGTCAAGGACCTCACCCAAGCCTGGGA
CCTCTATTATCATGTGTTCCGACGAATCTCAAAGCAGGTCGGAGGCAGTGGAGGCGGG
GGCTCAGGAGGGTCTTCCTCTGGGGGTATGGTGTTTACTCTGGAAGACTTTGTTGGTGA
TTGGGAGCAGACTGCAGCATACAACCTGGATCAAGTCTTGGAGCAGGGTGGTGTATCC
TCCCTTCTCCAGAACTTGGCAGTCTCCGTCACACCTATCCAACGGATCGTTAGGTCTGG
TGAAAACGCCTTGAAAATTGATATCCACGTTATTATCCCGTACGAAGGTCTGTCCGCTGA
CCAGATGGCGCAGATAGAAGAAGTTTTCAAGGTTGTTTACCCGGTCGACGATCATCACT
TCAAAGTCATATTGCCATATGGAACGCTGGTTATAGACGGTGTCACGCCTAACATGCTTA
ACTACTTCGGTCGGCCGTACGAGGGTATAGCTGTATTTGATGGTAAGAAAATTACAGTC
ACGGGCACTTTGTGGAACGGAAATAAAATCATCGATGAACGACTGATCACTCCTGACGG
GTCTATGCTCTTCCGCGTAACAATTAATTCCTAACCCGGGatccgcccctctccctcccccccccctaa
cgttactggccgaagccgcttggaataaggccggtgtgcgtttgtctatatgttattttccaccatattgccgtcttttggcaatgtgagg
gcccggaaacctggccctgtcttcttgacgagcattcctaggggtctttcccctctcgccaaaggaatgcaaggtctgttgaatgtcg
tgaaggaagcagttcctctggaagcttcttgaagacaaacaacgtctgtagcgaccctttgcaggcagcggaaccccccacctg
gcgacaggtgcctctgcggccaaaagccacgtgtataagatacacctgcaaaggcggcacaaccccagtgccacgttgtgagt
tggatagttgtggaaagagtcaaatggctctcctcaagcgtattcaacaaggggctgaaggatgcccagaaggtaccccattgta
tgggatctgatctggggcctcggtgcacatgctttacatgtgtttagtcgaggttaaaaaacgtctaggccccccgaaccacggg
gacgtggttttcctttgaaaaacacgatgataatatggCCACaaccATGcgatcgTACCCATACGATGTTCCAGA
TTACGCTgaattcatggtgagcaagggcgaggagctgttcaccggggtggtgcccatcctggtcgagctggacggcgacgt
aaacggccacaagttcagcgtgtccggcgagggcgagggcgatgccacctacggcaagctgaccctgaagttcatctgcacc
accggcaagctgcccgtgccctggcccaccctcgtgaccaccctgtcctggggcgtgcagtgcttcgcccgctaccccgaccac
atgaagcagcacgacttcttcaagtccgccatgcccgaaggctacgtccaggagcgcaccatcttcttcaaggacgacggcaa
ctacaagacccgcgccgaggtgaagttcgagggcgacaccctggtgaaccgcatcgagctgaagggcatcgacttcaagga
ggacggcaacatcctggggcacaagctggagtacaactacttcagcgacaacgtctatatcaccgccgacaagcagaagaac
ggcatcaaggccaacttcaagatccgccacaacatcgaggacggcggcgtgcagctcgccgaccactaccagcagaacacc
cccatcggcgacggccccgtgctgctgcccgacaaccactacctgagcacccagtccaagctgagcaaagaccccaacgag
aagcgcgatcacatggtcctgctggagttcgtgaccgccgccgggatcactctcggcatggacgagctgtacaagtaa

pAG1474 FKBP-SmBiT_IRES_HA_IRFP670

ATGGGAGTGCAGGTGGAAACCATCTCCCCAGGAGACGGGCGCACCTTCCCCAAGCGC
GGCCAGACCTGCGTGGTGCACTACACCGGGATGCTTGAAGATGGAAAGAAATTTGATT
CCTCCCGGGACAGAAACAAGCCCTTTAAGTTTATGCTAGGCAAGCAGGAGGTGATCCG
AGGCTGGGAAGAAGGGGTTGCCCAGATGAGTGTGGGTCAGAGAGCCAAACTGACTATA
TCTCCAGATTATGCCTATGGTGCCACTGGGCACCCAGGCATCATCCCACCACATGCCAC
TCTCGTCTTCGATGTGGAGCTTCTAAAACTGGAAGAAGGTGGTTCCGGAGGCGGTGGA
AGCGGAGGCTCATCATCCGGCGGAGTAACCGGTTACAGGTTGTTTGAAGAGATACTCTA
ACTCGAGGACTACAAGGACGACGACGACAAGCCCGGGatccGcccctctccctcccccccccctaac
gttactggccgaagccgcttggaataaggccggtgtgcgtttgtctatatgttattttccaccatattgccgtcttttggcaatgtgaggg
cccggaaacctggccctgtcttcttgacgagcattcctaggggtctttcccctctcgccaaaggaatgcaaggtctgttgaatgtcgt
gaaggaagcagttcctctggaagcttcttgaagacaaacaacgtctgtagcgaccctttgcaggcagcggaaccccccacctgg
cgacaggtgcctctgcggccaaaagccacgtgtataagatacacctgcaaaggcggcacaaccccagtgccacgttgtgagtt
ggatagttgtggaaagagtcaaatggctctcctcaagcgtattcaacaaggggctgaaggatgcccagaaggtaccccattgtat
gggatctgatctggggcctcggtgcacatgctttacatgtgtttagtcgaggttaaaaaaacgtctaggccccccgaaccacgggg
acgtggttttcctttgaaaaacacgatgataatatggCCACaaccATGcgatcgTACCCATACGATGTTCCAGA
TTACGCTGaattcatggcgcgtaaggtcgatctcacctcctgcgatcgcgagccgatccacatccccggcagcattcagcc
gtgcggctgcctgctagcctgcgacgcgcaggcggtgcggatcacgcgcattacggaaaatgccggcgcgttctttggacgcga
aactccgcgggtcggtgagctactcgccgattacttcggcgagaccgaagcccatgcgctgcgcaacgcactggcgcagtcttc
cgatccaaagcgaccggcgctgatcttcggttggcgcgacggcctgaccggccgcaccttcgacatctcactgcatcgccatga
cggtacatcgatcatcgagttcgagcctgcggcggccgaacaggccgacaatccgctgcggctgacgcggcagatcatcgcg
cgcaccaaagaactgaagtcgctcgaagagatggccgcacgggtgccgcgctatctgcaggcgatgctcggctatcaccgcgt
gatgttgtaccgcttcgcggacgacggctccgggatggtgatcggcgaggcgaagcgcagcgacctcgagagctttctcggtca
gcactttccggcgtcgctggtcccgcagcaggcgcggctactgtacttgaagaacgcgatccgcgtggtctcggattcgcgcggc
atcagcagccggatcgtgcccgagcacgacgcctccggcgccgcgctcgatctgtcgttcgcgcacctgcgcagcatctcgccc
tgccatctcgaatttctgcggaacatgggcgtcagcgcctcgatgtcgctgtcgatcatcattgacggcacgctatggggattgatc
atctgtcatcattacgagccgcgtgccgtgccgatggcgcagcgcgtcgcggccgaaatgttcgccgacttcttatcgctgcacttc
accgccgcccaccaccaacgcTAA

pAG1494 CMV_CMyc_LgBiT-$^{FIRE}$tag_IRES_HA_iRFP670

ATGGAACAAAAGCTTATTTCTGAAGAGGACTTGgaattcATGGTGTTTACATTGGAGGATTT
TGTTGGCGATTGGGAACAGACTGCGGCATATAATCTTGATCAGGTCTTGGAACAAGGCG
GGGTTTCATCCCTTTTGCAAAATCTGGCAGTATCTGTTACCCCCATACAGCGAATAGTAC
GCTCAGGCGAGAACGCCTTGAAAATTGATATCCATGTAATCATACCCTACGAAGGCCTG
AGTGCAGACCAGATGGCTCAAATAGAAGAAGTCTTTAAGGTAGTCTACCCTGTCGACGA
TCATCATTTCAAGGTTATCTTGCCATACGGTACGCTGGTTATAGATGGCGTCACCCCAAA
TATGCTTAACTATTTCGGACGGCCCTATGAAGGGATAGCAGTTTTCGACGGCAAAAAGA
TTACGGTCACTGGGACTCTGTGGAACGGTAACAAAATTATAGACGAGCGACTCATTACG
CCTGATGGATCAATGCTGTTCCGGGTGACTATAAACTCTGGTGGCAGCGGTGGAGGTG
GCTCCGGAGGAAGCTCATCCGGAGGTGGAGACAGATATTGGGTATTTGTTAAGCGCGT
TTAACTCGAGGACTACAAGGACGACGACGACAAGCCCGGGatccGcccctctccctcccccccccct
aacgttactggccgaagccgcttggaataaggccggtgtgcgtttgtctatatgttattttccaccatattgccgtcttttggcaatgtga
gggcccggaaacctggccctgtcttcttgacgagcattcctaggggtctttcccctctcgccaaaggaatgcaaggtctgttgaatgt
cgtgaaggaagcagttcctctggaagcttcttgaagacaaacaacgtctgtagcgaccctttgcaggcagcggaaccccccacc
tggcgacaggtgcctctgcggccaaaagccacgtgtataagatacacctgcaaaggcggcacaaccccagtgccacgttgtga
gttggatagttgtggaaagagtcaaatggctctcctcaagcgtattcaacaaggggctgaaggatgcccagaaggtaccccattg
tatgggatctgatctggggcctcggtgcacatgctttacatgtgtttagtcgaggttaaaaaaacgtctaggccccccgaaccacgg
ggacgtggttttcctttgaaaaacacgatgataatatggCCACaaccATGcgatcgTACCCATACGATGTTCCAG
ATTACGCTGaattcatggcgcgtaaggtcgatctcacctcctgcgatcgcgagccgatccacatccccggcagcattcagc
cgtgcggctgcctgctagcctgcgacgcgcaggcggtgcggatcacgcgcattacggaaaatgccggcgcgttctttggacgcg
aaactccgcgggtcggtgagctactcgccgattacttcggcgagaccgaagcccatgcgctgcgcaacgcactggcgcagtctt
ccgatccaaagcgaccggcgctgatcttcggttggcgcgacggcctgaccggccgcaccttcgacatctcactgcatcgccatg
acggtacatcgatcatcgagttcgagcctgcggcggccgaacaggccgacaatccgctgcggctgacgcggcagatcatcgc
gcgcaccaaagaactgaagtcgctcgaagagatggccgcacgggtgccgcgctatctgcaggcgatgctcggctatcaccgc
gtgatgttgtaccgcttcgcggacgacggctccgggatggtgatcggcgaggcgaagcgcagcgacctcgagagctttctcggt
cagcactttccggcgtcgctggtcccgcagcaggcgcggctactgtacttgaagaacgcgatccgcgtggtctcggattcgcgcg
gcatcagcagccggatcgtgcccgagcacgacgcctccggcgccgcgctcgatctgtcgttcgcgcacctgcgcagcatctcgc
cctgccatctcgaatttctgcggaacatgggcgtcagcgcctcgatgtcgctgtcgatcatcattgacggcacgctatggggattgat
catctgtcatcattacgagccgcgtgccgtgccgatggcgcagcgcgtcgcggccgaaatgttcgccgacttcttatcgctgcactt
caccgccgcccaccaccaacgcTAA

pAG1495 CMV_CMyc_FIREmate_SmBiT_IRES_HA_mTQ2

ATGGAACAAAAGCTTATTTCTGAAGAGGACTTGGaattcATGGAGCATGTCGCATTCGGGT
CAGAAGATATCGAAAACACATTGGCTAATATGGATGACGAACAGCTGGATCGGCTCGCT
TTCGGTGTCATACAATTGGACGGGGATGGCAACATTTTGTTGTATAATGCAGCAGAAGG
TGATATAACTGGACGGGACCCGAAGCAAGTGATCGGCAAGAACTTCTTTAAGGATGTCG
CCCCCGGTACCGACACGCCTGAGTTCTATGGAAAATTCAAGGAGGGAGCTGCCAGTGG
GAATTTGAACACGATGTTTGAGTGGACAATCCCGACGTCAAGGGGTCCGACCAAAGTCA
AAGTGCATCTCAAGAAGGCACTGTCTGGGGGTAGCGGTGGCGGAGGGTCTGGGGGTT
CATCCTCAGGCGGTGTCACCGGGTATCGCTTGTTTGAGGAAATTCTCTAACCTCGAGGA
CTACAAGGACGACGACGACAAGCCCGGGatccGcccctctccctcccccccccctaacgttactggccgaa
gccgcttggaataaggccggtgtgcgtttgtctatatgttattttccaccatattgccgtcttttggcaatgtgagggcccggaaacctg
gccctgtcttcttgacgagcattcctaggggtctttcccctctcgccaaaggaatgcaaggtctgttgaatgtcgtgaaggaagcagt
tcctctggaagcttcttgaagacaaacaacgtctgtagcgaccctttgcaggcagcggaaccccccacctggcgacaggtgcctc
tgcggccaaaagccacgtgtataagatacacctgcaaaggcggcacaaccccagtgccacgttgtgagttggatagttgtggaa
agagtcaaatggctctcctcaagcgtattcaacaaggggctgaaggatgcccagaaggtaccccattgtatgggatctgatctgg
ggcctcggtgcacatgctttacatgtgtttagtcgaggttaaaaaaacgtctaggccccccgaaccacggggacgtggttttcctttg
aaaaacacgatgataatatggCCACaaccATGcgatcgTACCCATACGATGTTCCAGATTACGCTgaatt
catggtgagcaagggcgaggagctgttcaccggggtggtgcccatcctggtcgagctggacggcgacgtaaacggccacaag
ttcagcgtgtccggcgagggcgagggcgatgccacctacggcaagctgaccctgaagttcatctgcaccaccggcaagctgcc
cgtgccctggcccaccctcgtgaccaccctgtcctggggcgtgcagtgcttcgcccgctaccccgaccacatgaagcagcacga
cttcttcaagtccgccatgcccgaaggctacgtccaggagcgcaccatcttcttcaaggacgacggcaactacaagacccgcgc
cgaggtgaagttcgagggcgacaccctggtgaaccgcatcgagctgaagggcatcgacttcaaggaggacggcaacatcctg
gggcacaagctggagtacaactacttcagcgacaacgtctatatcaccgccgacaagcagaagaacggcatcaaggccaact
tcaagatccgccacaacatcgaggacggcggcgtgcagctcgccgaccactaccagcagaacacccccatcggcgacggc
cccgtgctgctgcccgacaaccactacctgagcacccagtccaagctgagcaaagaccccaacgagaagcgcgatcacatg
gtcctgctggagttcgtgaccgccgccgggatcactctcggcatggacgagctgtacaagtaa

pAG1496 CMV_CMyc_[FIRE]tag_LgBiT_IRES_HA_iRFP670

ATGGAACAAAAGCTTATTTCTGAAGAGGACTTGGaattcGGAGATCGCTACTGGGTCTTCG
TGAAGAGAGTCGGAGGTTCCGGTGGAGGCGGTTCAGGTGGCTCTAGCAGCGGTGGTA
TGGTGTTCACACTTGAAGATTTCGTGGGTGATTGGGAGCAGACCGCAGCTTACAACCTC
GACCAGGTCTTGGAGCAAGGTGGTGTTAGTTCATTGCTCCAAAACCTCGCGGTGAGCG
TAACTCCAATCCAGCGCATAGTTCGGAGTGGGGAGAACGCCCTCAAGATAGATATTCAC
GTCATAATACCATACGAGGGTCTTTCTGCGGATCAAATGGCGCAGATAGAAGAGGTCTT
TAAGGTAGTCTACCCCGTAGATGATCATCACTTTAAAGTCATATTGCCCTATGGCACCCT
GGTAATAGACGGGGTGACGCCGAACATGTTGAATTACTTCGGGCGACCATACGAAGGA
ATCGCTGTCTTTGACGGAAAGAAGATTACAGTGACCGGGACCCTCTGGAATGGGAATAA
GATTATAGATGAAAGGTTGATCACACCAGATGGCAGTATGCTTTTTCGCGTCACGATAAA
CAGTTAACTCGAGGACTACAAGGACGACGACGACAAGCCCGGGatccGcccctctccctcccccc
ccctaacgttactggccgaagccgcttggaataaggccggtgtgcgtttgtctatatgttattttccaccatattgccgtcttttggcaa
tgtgagggcccggaaacctggccctgtcttcttgacgagcattcctaggggtctttcccctctcgccaaaggaatgcaaggtctgttg
aatgtcgtgaaggaagcagttcctctggaagcttcttgaagacaaacaacgtctgtagcgaccctttgcaggcagcggaacccc
ccacctggcgacaggtgcctctgcggccaaaagccacgtgtataagatacacctgcaaaggcggcacaaccccagtgccacg
ttgtgagttggatagttgtggaaagagtcaaatggctctcctcaagcgtattcaacaaggggctgaaggatgcccagaaggtacc
ccattgtatgggatctgatctggggcctcggtgcacatgctttacatgtgtttagtcgaggttaaaaaaacgtctaggccccccgaac
cacggggacgtggttttcctttgaaaaacacgatgataatatggCCACaaccATGcgatcgTACCCATACGATGTT
CCAGATTACGCTGaattcatggcgcgtaaggtcgatctcacctcctgcgatcgcgagccgatccacatccccggcagca
ttcagccgtgcggctgcctgctagcctgcgacgcgcaggcggtgcggatcacgcgcattacggaaaatgccggcgcgttctttgg
acgcgaaactccgcgggtcggtgagctactcgccgattacttcggcgagaccgaagcccatgcgctgcgcaacgcactggcgc
agtcttccgatccaaagcgaccggcgctgatcttcggttggcgcgacggcctgaccggccgcaccttcgacatctcactgcatcg
ccatgacggtacatcgatcatcgagttcgagcctgcggcggccgaacaggccgacaatccgctgcggctgacgcggcagatc
atcgcgcgcaccaaagaactgaagtcgctcgaagagatggccgcacgggtgccgcgctatctgcaggcgatgctcggctatca
ccgcgtgatgttgtaccgcttcgcggacgacggctccgggatggtgatcggcgaggcgaagcgcagcgacctcgagagctttct
cggtcagcactttccggcgtcgctggtcccgcagcaggcgcggctactgtacttgaagaacgcgatccgcgtggtctcggattcg
cgcggcatcagcagccggatcgtgcccgagcacgacgcctccggcgccgcgctcgatctgtcgttcgcgcacctgcgcagcat
ctcgccctgccatctcgaatttctgcggaacatgggcgtcagcgcctcgatgtcgctgtcgatcatcattgacggcacgctatgggg
attgatcatctgtcatcattacgagccgcgtgccgtgccgatggcgcagcgcgtcgcggccgaaatgttcgccgacttcttatcgct
gcacttcaccgccgcccaccaccaacgcTAA

pAG1497 CMV_CMyc_SmBiT-$^{FIRE}$mate_IRES_HA_mTQ2

ATGGAACAAAAGCTTATTTCTGAAGAGGACTTGGaattcGTAACCGGCTACCGACTGTTTG
AAGAAATACTCGGCGGTTCTGGTGGTGGTGGTTCAGGGGGCTCCAGCTCCGGTGGTAT
GGAGCATGTCGCTTTCGGATCAGAAGATATTGAAAACACCCTGGCGAATATGGACGATG
AGCAACTTGATCGGTTGGCTTTCGGGGTTATCCAGTTGGACGGTGATGGCAACATCCTG
TTGTACAACGCTGCTGAGGGGGATATTACAGGACGAGACCCCAAACAAGTGATAGGTAA
AAATTTCTTTAAAGACGTTGCTCCCGGAACAGATACGCCGGAATTCTATGGTAAGTTCAA
GGAAGGGGCAGCTTCCGGCAATTTGAACACCATGTTCGAGTGGACAATTCCTACCAGTC
GCGGTCCAACAAAAGTTAAAGTGCACCTTAAAAAAGCACTCTCTTAACCTCGAGGACTA
CAAGGACGACGACGACAAGCCCGGGatccGcccctctccctcccccccccctaacgttactggccgaagccg
cttggaataaggccggtgtgcgtttgtctatatgttattttccaccatattgccgtcttttggcaatgtgagggcccggaaacctggccct
gtcttcttgacgagcattcctaggggtctttcccctctcgccaaaggaatgcaaggtctgttgaatgtcgtgaaggaagcagttcctct
ggaagcttcttgaagacaaacaacgtctgtagcgaccctttgcaggcagcggaaccccccacctggcgacaggtgcctctgcg
gccaaaagccacgtgtataagatacacctgcaaaggcggcacaaccccagtgccacgttgtgagttggatagttgtggaaaga
gtcaaatggctctcctcaagcgtattcaacaaggggctgaaggatgcccagaaggtaccccattgtatgggatctgatctggggc
ctcggtgcacatgctttacatgtgtttagtcgaggttaaaaaaacgtctaggccccccgaaccacggggacgtggttttcctttgaaa
aacacgatgataatatggCCACaaccATGcgatcgTACCCATACGATGTTCCAGATTACGCTgaattcat
ggtgagcaagggcgaggagctgttcaccggggtggtgcccatcctggtcgagctggacggcgacgtaaacggccacaagttc
agcgtgtccggcgagggcgagggcgatgccacctacggcaagctgaccctgaagttcatctgcaccaccggcaagctgcccg
tgccctggcccaccctcgtgaccaccctgtcctggggcgtgcagtgcttcgcccgctaccccgaccacatgaagcagcacgactt
cttcaagtccgccatgcccgaaggctacgtccaggagcgcaccatcttcttcaaggacgacggcaactacaagacccgcgccg
aggtgaagttcgagggcgacaccctggtgaaccgcatcgagctgaagggcatcgacttcaaggaggacggcaacatcctggg
gcacaagctggagtacaactacttcagcgacaacgtctatatcaccgccgacaagcagaagaacggcatcaaggccaacttc
aagatccgccacaacatcgaggacggcggcgtgcagctcgccgaccactaccagcagaacacccccatcggcgacggccc
cgtgctgctgcccgacaaccactacctgagcacccagtccaagctgagcaaagaccccaacgagaagcgcgatcacatggtc
ctgctggagttcgtgaccgccgccgggatcactctcggcatggacgagctgtacaagtaa

pAG1511 CMV_FKBP-SmBiT-P2A-FRB-LgBiT_IRES_HA-iRFP670-CMyc

ATGGGAGTGCAGGTGGAAACCATCTCCCCAGGAGACGGGCGCACCTTCCCCAAGCGC
GGCCAGACCTGCGTGGTGCACTACACCGGGATGCTTGAAGATGGAAAGAAATTTGATT
CCTCCCGGGACAGAAACAAGCCCTTTAAGTTTATGCTAGGCAAGCAGGAGGTGATCCG
AGGCTGGGAAGAAGGGGTTGCCCAGATGAGTGTGGGTCAGAGAGCCAAACTGACTATA
TCTCCAGATTATGCCTATGGTGCCACTGGGCACCCAGGCATCATCCCACCACATGCCAC
TCTCGTCTTCGATGTGGAGCTTCTAAAACTGGAAGAAGGTGGTTCCGGAGGCGGTGGA
AGCGGAGGCTCATCATCCGGCGGAGTAACCGGTTACAGGTTGTTTGAAGAGATACTCG
GAAGCGGAGCTACTAACTTCAGCCTGCTGAAGCAGGCTGGAGACGTGGAGGAGAACCC
TGGACCTGAGATGTGGCATGAAGGCCTGGAAGAGGCATCTCGTTTGTACTTTGGGGAA
AGGAACGTGAAAGGCATGTTTGAGGTGCTGGAGCCCTTGCATGCTATGATGGAACGGG
GCCCCCAGACTCTGAAGGAAACATCCTTTAATCAGGCCTATGGTCGAGATTTAATGGAG
GCCCAAGAGTGGTGCAGGAAGTACATGAAATCAGGGAATGTCAAGGACCTCACCCAAG
CCTGGGACCTCTATTATCATGTGTTCCGACGAATCTCAAAGCAGGTCGGAGGCAGTGGA
GGCGGGGGCTCAGGAGGGTCTTCCTCTGGGGGTATGGTGTTTACTCTGGAAGACTTTG
TTGGTGATTGGGAGCAGACTGCAGCATACAACCTGGATCAAGTCTTGGAGCAGGGTGG
TGTATCCTCCCTTCTCCAGAACTTGGCAGTCTCCGTCACACCTATCCAACGGATCGTTA
GGTCTGGTGAAAACGCCTTGAAAATTGATATCCACGTTATTATCCCGTACGAAGGTCTGT
CCGCTGACCAGATGGCGCAGATAGAAGAAGTTTTCAAGGTTGTTTACCCGGTCGACGAT
CATCACTTCAAAGTCATATTGCCATATGGAACGCTGGTTATAGACGGTGTCACGCCTAA
CATGCTTAACTACTTCGGTCGGCCGTACGAGGGTATAGCTGTATTTGATGGTAAGAAAA
TTACAGTCACGGGCACTTTGTGGAACGGAAATAAAATCATCGATGAACGACTGATCACT
CCTGACGGGTCTATGCTCTTCCGCGTAACAATTAATTCCAACCCGGGatccgcccctctccctcc
cccccccctaacgttactggccgaagccgcttggaataaggccggtgtgcgtttgtctatatgttattttccaccatattgccgtcttttg
gcaatgtgagggcccggaaacctggccctgtcttcttgacgagcattcctaggggtctttcccctctcgccaaaggaatgcaaggt
ctgttgaatgtcgtgaaggaagcagttcctctggaagcttcttgaagacaaacaacgtctgtagcgaccctttgcaggcagcggaa
ccccccacctggcgacaggtgcctctgcggccaaaagccacgtgtataagatacacctgcaaaggcggcacaaccccagtgc
cacgttgtgagttggatagttgtggaaagagtcaaatggctctcctcaagcgtattcaacaaggggctgaaggatgcccagaagg
taccccattgtatgggatctgatctggggcctcggtgcacatgctttacatgtgtttagtcgaggttaaaaaacgtctaggccccc
gaaccacggggacgtggttttcctttgaaaaacacgatgataatatggCCACaaccATGcgatcgTACCCATACGAT
GTTCCAGATTACGCTGaattcatggcgcgtaaggtcgatctcacctcctgcgatcgcgagccgatccacatccccggc
agcattcagccgtgcggctgcctgctagcctgcgacgcgcaggcggtgcggatcacgcgcattacggaaaatgccggcgtt
ctttggacgcgaaactccgcgggtcggtgagctactcgccgattacttcggcgagaccgaagcccatgcgctgcgcaacgcact
ggcgcagtcttccgatccaaagcgaccggcgctgatcttcggttggcgcgacggcctgaccggccgcaccttcgacatctcactg
catcgccatgacggtacatcgatcatcgagttcgagcctgcggcggccgaacaggccgacaatccgctgcggctgacgcggc
agatcatcgcgcgcaccaaagaactgaagtcgctcgaagagatggccgcacgggtgccgcgctatctgcaggcgatgctcgg
ctatcaccgcgtgatgttgtaccgcttcgcggacgacggctccgggatggtgatcggcgaggcgaagcgcagcgacctcgaga
gctttctcggtcagcactttccggcgtcgctggtcccgcagcaggcgcggctactgtacttgaagaacgcgatccgcgtggtctcg
gattcgcgcggcatcagcagccggatcgtgcccgagcacgacgcctccggcgccgcgctcgatctgtcgttcgcgcacctgcgc
agcatctcgccctgccatctcgaatttctgcggaacatgggcgtcagcgcctcgatgtcgctgtcgatcatcattgacggcacgcta
tggggattgatcatctgtcatcattacgagccgcgtgccgtgccgatggcgcagcgcgtcgcggccgaaatgttcgccgacttctta
tcgctgcacttcaccgccgcccaccaccaacgcGAACAAAAGCTTATTTCTGAAGAGGACTTGTAA

pAG1512 CMV_$^{FIRE}$mate-SmBiT-P2A-LgBiT-$^{FIRE}$tag_IRES_HA-iRFP-CMyc

ATGGAGCATGTCGCATTCGGGTCAGAAGATATCGAAAACACATTGGCTAATATGGATGA
CGAACAGCTGGATCGGCTCGCTTTCGGTGTCATACAATTGGACGGGGATGGCAACATTT
TGTTGTATAATGCAGCAGAAGGTGATATAACTGGACGGGACCCGAAGCAAGTGATCGG
CAAGAACTTCTTTAAGGATGTCGCCCCCGGTACCGACACGCCTGAGTTCTATGGAAAAT
TCAAGGAGGGAGCTGCCAGTGGGAATTTGAACACGATGTTTGAGTGGACAATCCCGAC
GTCAAGGGGTCCGACCAAAGTCAAAGTGCATCTCAAGAAGGCACTGTCTGGGGGTAGC
GGTGGCGGAGGGTCTGGGGGTTCATCCTCAGGCGGTGTCACCGGGTATCGCTTGTTTG
AGGAAATTCTCGGAAGCGGAGCTACTAACTTCAGCCTGCTGAAGCAGGCTGGAGACGT
GGAGGAGAACCCTGGACCTATGGTGTTTACATTGGAGGATTTTGTTGGCGATTGGGAAC
AGACTGCGGCATATAATCTTGATCAGGTCTTGGAACAAGGCGGGGTTTCATCCCTTTTG
CAAAATCTGGCAGTATCTGTTACCCCCATACAGCGAATAGTACGCTCAGGCGAGAACGC
CTTGAAAATTGATATCCATGTAATCATACCCTACGAAGGCCTGAGTGCAGACCAGATGG
CTCAAATAGAAGAAGTCTTTAAGGTAGTCTACCCTGTCGACGATCATCATTTCAAGGTTA
TCTTGCCATACGGTACGCTGGTTATAGATGGCGTCACCCCAAATATGCTTAACTATTTCG
GACGGCCCTATGAAGGGATAGCAGTTTTCGACGGCAAAAAGATTACGGTCACTGGGAC
TCTGTGGAACGGTAACAAAATTATAGACGAGCGACTCATTACGCCTGATGGATCAATGC
TGTTCCGGGTGACTATAAACTCTGGTGGCAGCGGTGGAGGTGGCTCCGGAGGAAGCTC
ATCCGGAGGTGGAGACAGATATTGGGTATTTGTTAAGCGCGTTTAACCCGGGatccgcccct
ctccctcccccccccctaacgttactggccgaagccgcttggaataaggccggtgtgcgtttgtctatatgttattttccaccatattgc
cgtcttttggcaatgtgagggcccggaaacctggccctgtcttcttgacgagcattcctaggggtctttcccctctcgccaaaggaat
gcaaggtctgttgaatgtcgtgaaggaagcagttcctctggaagcttcttgaagacaaacaacgtctgtagcgaccctttgcaggc
agcggaaccccccacctggcgacaggtgcctctgcggccaaaagccacgtgtataagatacacctgcaaaggcggcacaac
cccagtgccacgttgtgagttggatagttgtggaaagagtcaaatggctctcctcaagcgtattcaacaaggggctgaaggatgc
ccagaaggtaccccattgtatgggatctgatctggggcctcggtgcacatgctttacatgtgtttagtcgaggttaaaaaaacgtcta
ggccccccgaaccacggggacgtggttttcctttgaaaaacacgatgataatatggCCACaaccATGcgatcgTACCCA
TACGATGTTCCAGATTACGCTGaattcatggcgcgtaaggtcgatctcacctcctgcgatcgcgagccgatccaca
tccccggcagcattcagccgtgcggctgcctgctagcctgcgacgcgcaggcggtgcggatcacgcgcattacggaaaatgcc
ggcgcgttctttggacgcgaaactccgcgggtcggtgagctactcgccgattacttcggcgagaccgaagcccatgcgctgcgc
aacgcactggcgcagtcttccgatccaaagcgaccggcgctgatcttcggttggcgcgacggcctgaccggccgcaccttcgac
atctcactgcatcgccatgacggtacatcgatcatcgagttcgagcctgcggcggccgaacaggccgacaatccgctgcggctg
acgcggcagatcatcgcgcgcaccaaagaactgaagtcgctcgaagagatggccgcacgggtgccgcgctatctgcaggcg
atgctcggctatcaccgcgtgatgttgtaccgcttcgcggacgacggctccgggatggtgatcggcgaggcgaagcgcagcgac
ctcgagagctttctcggtcagcactttccggcgtcgctggtcccgcagcaggcgcggctactgtacttgaagaacgcgatccgcgt
ggtctcggattcgcgcggcatcagcagccggatcgtgcccgagcacgacgcctccggcgccgcgctcgatctgtcgttcgcgca
cctgcgcagcatctcgccctgccatctcgaatttctgcggaacatgggcgtcagcgcctcgatgtcgctgtcgatcatcattgacgg
cacgctatggggattgatcatctgtcatcattacgagccgcgtgccgtgccgatggcgcagcgcgtcgcggccgaaatgttcgcc
gacttcttatcgctgcacttcaccgccgcccaccaccaacgcGAACAAAAGCTTATTTCTGAAGAGGACTTGT
AA

pAG1611 CMV_GAL4-$^{FIRE}$mate

ATGAAGCTACTGTCTTCTATCGAACAAGCATGCGATATTTGCCGACTTAAAAGCTCAAG
TGCTCCAAAGAAAAACCGAAGTGCGCCAAGTGTCTGAAGAACAACTGGGAGTGTCGCT
ACTCTCCCAAAACCAAAAGGTCTCCGCTGACTAGGGCACATCTGACAGAAGTGGAATCA
AGGCTAGAAAGACTGGAACAGCTATTTCTACTGATTTTTCCTCGAGAAGACCTTGACATG
ATTTTGAAAATGGATTCTTTACAGGATATAAAAGCATTGTTAACAGGATTATTTGTACAAG
ATAATGTGAATAAAGATGCCGTCACAGATAGATTGGCTTCAGTGGAGACTGATATGCCT
CTAACATTGAGACAGCATAGAATAAGTGCGACATCATCATCGGAAGAGAGTAGTAACAA
AGGTCAAAGACAGTTGACTGTAGCGATTCCGTCGACACCACCTACTCCCTCTCCAGCGA
TCGCCATGGAAATGGAGCATGTCGCATTCGGGTCAGAAGATATCGAAAACACATTGGCT
AATATGGATGACGAACAGCTGGATCGGCTCGCTTTCGGTGTCATACAATTGGACGGGG
ATGGCAACATTTTGTTGTATAATGCAGCAGAAGGTGATATAACTGGACGGGACCCGAAG
CAAGTGATCGGCAAGAACTTCTTTAAGGATGTCGCCCCCGGTACCGACACGCCTGAGTT
CTATGGAAAATTCAAGGAGGGAGCTGCCAGTGGGAATTTGAACACGATGTTTGAGTGGA
CAATCCCGACGTCAAGGGGTCCGACCAAAGTCAAAGTGCATCTCAAGAAGGCACTGTC
T

pAG1612 CMV_GAL4-$^{FIRE}$tag

ATGAAGCTACTGTCTTCTATCGAACAAGCATGCGATATTTGCCGACTTAAAAGCTCAAG
TGCTCCAAAGAAAAACCGAAGTGCGCCAAGTGTCTGAAGAACAACTGGGAGTGTCGCT
ACTCTCCCAAAACCAAAAGGTCTCCGCTGACTAGGGCACATCTGACAGAAGTGGAATCA
AGGCTAGAAAGACTGGAACAGCTATTTCTACTGATTTTTCCTCGAGAAGACCTTGACATG
ATTTTGAAAATGGATTCTTTACAGGATATAAAAGCATTGTTAACAGGATTATTTGTACAAG
ATAATGTGAATAAAGATGCCGTCACAGATAGATTGGCTTCAGTGGAGACTGATATGCCT
CTAACATTGAGACAGCATAGAATAAGTGCGACATCATCATCGGAAGAGAGTAGTAACAA
AGGTCAAAGACAGTTGACTGTAGCGATTCCGTCGACACCACCTACTCCCTCTCCAGCGA
TCGCCATGGAAGGAGACAGATATTGGGTATTTGTTAAGCGCGTT

pAG1614 CMV_VP16-$^{FIRE}$mate

ATGAAGCTACTGTCTTCTATCGAACAAGCATGCCCAAAAAAGAAGAGAAAGGTAGATGA
ATTTCCTGGGATCTCTACTGCTCCTCCAACCGATGTCAGCCTGGGCGACGAACTCCACT
TAGACGGCGAGGACGTGGCGATGGCGCATGCCGACGCGCTAGACGATTTCGATCTGG
ACATGTTGGGGGACGGGGATTCCCCGGGTCCGGGATCTCCAGCGATTCCGTCGACAC
CACCTACTCCCTCTCCAGCGATCGCCATGGAAATGGAGCATGTCGCATTCGGGTCAGA
AGATATCGAAAACACATTGGCTAATATGGATGACGAACAGCTGGATCGGCTCGCTTTCG
GTGTCATACAATTGGACGGGGATGGCAACATTTTGTTGTATAATGCAGCAGAAGGTGAT
ATAACTGGACGGGACCCGAAGCAAGTGATCGGCAAGAACTTCTTTAAGGATGTCGCCC
CCGGTACCGACACGCCTGAGTTCTATGGAAAATTCAAGGAGGGAGCTGCCAGTGGGAA
TTTGAACACGATGTTTGAGTGGACAATCCCGACGTCAAGGGGTCCGACCAAAGTCAAAG
TGCATCTCAAGAAGGCACTGTCTTAA

pAG1615 CMV_VP16-$^{\text{FIRE}}$tag

ATGAAGCTACTGTCTTCTATCGAACAAGCATGCCCAAAAAAGAAGAGAAAGGTAGATGA
ATTTCCTGGGATCTCTACTGCTCCTCCAACCGATGTCAGCCTGGGCGACGAACTCCACT
TAGACGGCGAGGACGTGGCGATGGCGCATGCCGACGCGCTAGACGATTTCGATCTGG
ACATGTTGGGGGACGGGGATTCCCCGGGTCCGGGATCTCCAGCGATTCCGTCGACAC
CACCTACTCCCTCTCCAGCGATCGCCATGGAAGGAGACAGATATTGGGTATTTGTTAAG
CGCGTT TAA

pAG1687 CMV_AU1-$^{\text{FIRE}}$tag-NTEVp

atggacacctacaggtacatcGGAGACAGATATTGGGTATTTGTTAAGCGCGTTggatccggaagtgga
gaaagcttgtttaagggaccacgtgattacaacccgatatcgagcaccatttgtcacttgacgaatgaatctgatgggcacacaac
atcgttgtatggtattggatttggtcccttcatcattacaaacaagcacttgtttagaagaaataatggaacactgttggtccaatcact
acatggtgtattcaaggtcaagaacaccacgactttgcaacaacacctcattgatgggagggacatgataattattcgcatgccta
aggatttcccaccatttcctcaaaagctgaaatttagagagccacaaagggaagagcgcatatgtcttgtgacaaccaacttcca
aacttaa

pAG1688 CMV_ AU1-$^{\text{FIRE}}$mate-NTEVp

atggacacctacaggtacatcATGGAGCATGTCGCATTCGGGTCAGAAGATATCGAAAACACATT
GGCTAATATGGATGACGAACAGCTGGATCGGCTCGCTTTCGGTGTCATACAATTGGACG
GGGATGGCAACATTTTGTTGTATAATGCAGCAGAAGGTGATATAACTGGACGGGACCCG
AAGCAAGTGATCGGCAAGAACTTCTTTAAGGATGTCGCCCCCGGTACCGACACGCCTG
AGTTCTATGGAAAATTCAAGGAGGGAGCTGCCAGTGGGAATTTGAACACGATGTTTGAG
TGGACAATCCCGACGTCAAGGGGTCCGACCAAAGTCAAAGTGCATCTCAAGAAGGCAC
TGTCTggatccggaagtggagaaagcttgtttaagggaccacgtgattacaacccgatatcgagcaccatttgtcacttgacg
aatgaatctgatgggcacacaacatcgttgtatggtattggatttggtcccttcatcattacaaacaagcacttgtttagaagaaataa
tggaacactgttggtccaatcactacatggtgtattcaaggtcaagaacaccacgactttgcaacaacacctcattgatgggaggg
acatgataattattcgcatgcctaaggatttcccaccatttcctcaaaagctgaaatttagagagccacaaagggaagagcgcata
tgtcttgtgacaaccaacttccaaacttaa

pAG1689 CMV_CMyc-$^{\text{FIRE}}$tag-CTEVp

ATGgagcagaagctgatcagcgaggaggatctgGGAGACAGATATTGGGTATTTGTTAAGCGCGTT
ggatccggctccggcagcaagagcatgtctagcatggtgtcagacaccagttgcacattcccttcatctgatggcatattctggaa
gcattggattcaaaccaaggatgggcagtgtggcagtccattagtatcaactagagatgggttcattgttggtatacactcagcatc
gaatttcaccaacacaaacaattatttcacaagcgtgccgaaaaacttcatggaattgttgacaaatcaggaggcgcagcagtg
ggttagtggttggcgattaaatgctgactcagtattgtgggggggccataaagttttcatgagcaaacctgaagagccttttcagcca
gttaaggaagcgactcaactcatgagtgaattggtgtactcgcaataa

pAG1690 CMV_CMyc-FIREmate-CTEVp

ATGgagcagaagctgatcagcgaggaggatctgATGGAGCATGTCGCATTCGGGTCAGAAGATATCG
AAAACACATTGGCTAATATGGATGACGAACAGCTGGATCGGCTCGCTTTCGGTGTCATA
CAATTGGACGGGGATGGCAACATTTTGTTGTATAATGCAGCAGAAGGTGATATAACTGG
ACGGGACCCGAAGCAAGTGATCGGCAAGAACTTCTTTAAGGATGTCGCCCCCGGTACC
GACACGCCTGAGTTCTATGGAAAATTCAAGGAGGGAGCTGCCAGTGGGAATTTGAACA
CGATGTTTGAGTGGACAATCCCGACGTCAAGGGGTCCGACCAAAGTCAAAGTGCATCT
CAAGAAGGCACTGTCTggatccggctccggcagcaagagcatgtctagcatggtgtcagacaccagttgcacattcc
cttcatctgatggcatattctggaagcattggattcaaaccaaggatgggcagtgtggcagtccattagtatcaactagagatgggtt
cattgttggtatacactcagcatcgaatttcaccaacacaaacaattatttcacaagcgtgccgaaaaacttcatggaattgttgaca
aatcaggaggcgcagcagtgggttagtggttggcgattaaatgctgactcagtattgtggggggccataaagttttcatgagcaa
acctgaagagccttttcagccagttaaggaagcgactcaactcatgagtgaattggtgtactcgcaataa

pAG1691 CMV_CMyc-Renilla_IRES_HA-iRFP670

ATGGAACAAAAGCTTATTTCTGAAGAGGACTTGgaattc
atgacttcgaaagtttatgatccagaacaaaggaaacggatgataactggtccgcagtggtgggccagatgtaaacaaatgaat
gttcttgattcatttattaattattatgattcagaaaaacatgcagaaaatgctgttattttttacatggtaacgcggcctcttcttatttatg
gcgacatgttgtgccacatattgagccagtagcgcggtgtattataccagaccttattggtatgggcaaatcaggcaaatctggtaa
tggttcttataggttacttgatcattacaaatatcttactgcatggtttgaacttcttaatttaccaaagaagatcattttgtcggccatgatt
ggggtgcttgtttggcatttcattatagctatgagcatcaagataagatcaaagcaatagttcacgctgaaagtgtagtagatgtgatt
gaatcatgggatgaatggcctgatattgaagaagatattgcgttgatcaaatctgaagaaggagaaaaaatggttttggagaata
acttcttcgtggaaaccatgttgccatcaaaaatcatgagaaagttagaaccagaagaatttgcagcatatcttgaaccattcaaa
gagaaaggtgaagttcgtcgtccaacattatcatggcctcgtgaaatcccgttagtaaaaggtggtaaacctgacgttgtacaaatt
gttaggaattataatgcttatctacgtgcaagtgatgatttaccaaaaatgtttattgaatcggacccaggattcttttccaatgctattgtt
gaaggtgccaagaagtttcctaatactgaatttgtcaaagtaaaaggtcttcatttttcgcaagaagatgcacctgatgaaatggga
aaatatatcaaatcgttcgttgagcgagttctcaaaaatgaacaatgaTAACTCGAGGACTACAAGGACGACGA
CGACAAGCCCGGGatccgcccctctccctcccccccccctaacgttactggccgaagccgcttggaataaggccggtgt
gcgtttgtctatatgttattttccaccatattgccgtcttttggcaatgtgagggcccggaaacctggccctgtcttcttgacgagcattcct
aggggtctttcccctctcgccaaaggaatgcaaggtctgttgaatgtcgtgaaggaagcagttcctctggaagcttcttgaagacaa
acaacgtctgtagcgaccctttgcaggcagcggaaccccccacctggcgacaggtgcctctgcggccaaaagccacgtgtata
agatacacctgcaaaggcggcacaaccccagtgccacgttgtgagttggatagttgtggaaagagtcaaatggctctcctcaag
cgtattcaacaaggggctgaaggatgcccagaaggtaccccattgtatgggatctgatctggggcctcggtgcacatgctttacat
gtgtttagtcgaggttaaaaaaacgtctaggccccccgaaccacggggacgtggttttcctttgaaaaacacgatgataatatggC
CACaaccATGcgatcgTACCCATACGATGTTCCAGATTACGCTgaattcatggcgcgtaaggtcgatctca
cctcctgcgatcgcgagccgatccacatccccggcagcattcagccgtgcggctgcctgctagcctgcgacgcgcaggcggtgc
ggatcacgcgcattacggaaaatgccggcgcgttctttggacgcgaaactccgcgggtcggtgagctactcgccgattacttcgg
cgagaccgaagcccatgcgctgcgcaacgcactggcgcagtcttccgatccaaagcgaccggcgctgatcttcggttggcgcg
acggcctgaccggccgcaccttcgacatctcactgcatcgccatgacggtacatcgatcatcgagttcgagcctgcggcggccg
aacaggccgacaatccgctgcggctgacgcggcagatcatcgcgcgcaccaaagaactgaagtcgctcgaagagatggccg
cacgggtgccgcgctatctgcaggcgatgctcggctatcaccgcgtgatgttgtaccgcttcgcggacgacggctccgggatggt
gatcggcgaggcgaagcgcagcgacctcgagagctttctcggtcagcactttccggcgtcgctggtcccgcagcaggcgcggc
tactgtacttgaagaacgcgatccgcgtggtctcggattcgcgcggcatcagcagccggatcgtgcccgagcacgacgcctccg
gcgccgcgctcgatctgtcgttcgcgcacctgcgcagcatctcgccctgccatctcgaatttctgcggaacatgggcgtcagcgcc
tcgatgtcgctgtcgatcatcattgacggcacgctatggggattgatcatctgtcatcattacgagccgcgtgccgtgccgatggcgc
agcgcgtcgcggccgaaatgttcgccgacttcttatcgctgcacttcaccgccgcccaccaccaacgcTAA

pAG1735 CMV_DRD1-$^{FIRE}$tag-NTEVp-TEVcs-CMyc-GAL4-SV40NLS-VP16AD-2xHA

ATGaggactctgaacacctctgccatggacgggactgggctggtggtggagagggacttctctgttcgtatcctcactgcctgtttc
ctgtcgctgctcatcctgtccacgctcctggggaacacgctggtctgtgctgccgttatcaggttccgacacctgcggtccaaggtga
ccaacttctttgtcatctccttggctgtgtcagatctcttggtggccgtcctggtcatgccctggaaggcagtggctgagattgctggctt
ctggccctttgggtccttctgtaacatctgggtggcctttgacatcatgtgctccactgcatccatcctcaacctctgtgtgatcagcgtg
gacaggtattgggctatctccagccctttccggtatgagagaaagatgacccccaaggcagccttcatcctgatcagtgtggcatg
gaccttgtctgtactcatctccttcatcccagtgcagctcagctggcacaaggcaaaacccacaagcccctctgatggaaatgcca
cttccctggctgagaccatagacaactgtgactccagcctcagcaggacatatgccatctcatcctctgtaataagcttttacatccct
gtggccatcatgattgtcacctacaccaggatctacaggattgctcagaaacaaatacggcgcattgcggccttggagagggca
gcagtccacgccaagaattgccagaccaccacaggtaatggaaagcctgtcgaatgttctcaaccggaaagttcttttaagatgt
ccttcaaaagagaaactaaagtcctgaagactctgtcggtgatcatgggtgtgtttgtgtgctgttggctacctttcttcatcttgaactg
cattttgcccttctgtgggtctggggagacgcagcccttctgcattgattccaacacctttgacgtgtttgtgtggtttgggtgggctaatt
catccttgaaccccatcatttatgcctttaatgctgattttcggaaggcattttcaaccctcttaggatgctacagactttgccctgcgac
gaataatgccatagagacggtgagtatcaataacaatgggccgcgatgttttccagccatcatgagccacgaggctccatctcc
aaggagtgcaatctggtttacctgatcccacatgctgtgggctcctctgaggacctgaaaaaggaggaggcagctggcatcgcc
agacccttggagaagctgtccccagccctatcggtcatattggactatgacactgacgtctctctggagaagatccaacccatcac
acaaaacggtcagcacccaaccgacccagctttcttgtacaaagtggttgatgtcggaggtggaggtagcggaggtggaggta
gcGGAGACAGATATTGGGTATTTGTTAAGCGCGTTggatccggaagtggagaaagcttgtttaagggacc
acgtgattacaacccgatatcgagcaccatttgtcacttgacgaatgaatctgatgggcacacaacatcgttgtatggtattggattt
ggtcccttcatcattacaaacaagcacttgtttagaagaaataatggaacactgttggtccaatcactacatggtgtattcaaggtca
agaacaccacgactttgcaacaacacctcattgatgggagggacatgataattattcgcatgcctaaggatttcccaccatttcctc
aaaagctgaaatttagagagccacaaagggaagagcgcatatgtcttgtgacaaccaacttccaaactgagaacctgtacttcc
agggttctagagaacagaagctgatctcagaagaggatctgGcccgggccatgaagctactgtcttctatcgaacaagcatgcg
atatttgccgacttaaaaagctcaagtgctccaaagaaaaaccgaagtgcgccaagtgtctgaagaacaactgggagtgtcgct
actctcccaaaaccaaaaggtctccgctgactagggcacatctgacagaagtggaatcaaggctagaaagactggaacagct
atttctactgattttcctcgagaagaccttgacatgattttgaaaatggattctttacaggatataaaagcattgttaacaggattatttgt
acaagataatgtgaataaagatgccgtcacagatagattggcttcagtggagactgatatgcctctaacattgagacagcataga
ataagtgcgacatcatcatcggaagagagtagtaacaaaggtcaaagacagttgactgtatcgccggaattcccggggatcag
cttgataattcgattcaccatgcccaagaagaagaggaaggtggcgtacagccgcgcgcgtacgaaaaacaattacgggtcta
ccatcgagggcctgctcgatctcccggacgacgacgcccccgaagaggcggggctggcggctccgcgcctgtcctttctccccg
cgggacacacgcgcagactgtcgacgGcccccccgaccgatgtcagcctgggggacgagctccacttagacggcgaggac
gtcgcgatggcgcatgccgacgcgctagacgatttcgatctggacatgttgggggacggggattccccgggtccgggatttaccc
cccacgactccgccccctacggcgctctggatatggccgacttcgagtttgagcagatgtttaccgatgcccttggaattgacgagt
acggtgggggctacccatacgatgttcctgactatgcgggctatccctatgacgtcccggactatgcatga

pAG1738 10xUAS-MLPmin d2EGFP_PGK_iRFP713

atggtgagcaagggcgaggagctgttcaccggggtggtgcccatcctggtcgagctggacggcgacgtaaacggccacaagtt
cagcgtgtccggcgagggcgagggcgatgccacctacggcaagctgaccctgaagttcatctgcaccaccggcaagctgccc
gtgccctggcccaccctcgtgaccaccctgacctacggcgtgcagtgcttcagccgctaccccgaccacatgaagcagcacga
cttcttcaagtccgccatgcccgaaggctacgtccaggagcgcaccatcttcttcaaggacgacggcaactacaagacccgcgc
cgaggtgaagttcgagggcgacaccctggtgaaccgcatcgagctgaagggcatcgacttcaaggaggacggcaacatcctg
gggcacaagctggagtacaactacaacagccacaacgtctatatcatggccgacaagcagaagaacggcatcaaggtgaac
ttcaagatccgccacaacatcgaggacggcagcgtgcagctcgccgaccactaccagcagaacacccccatcggcgacggc
cccgtgctgctgcccgacaaccactacctgagcacccagtccgccctgagcaaagaccccaacgagaagcgcgatcacatg

gtcctgctggagttcgtgaccgccgccgggatcactctcggcatggacgagctgtacaagaagcttagccatggcttcccgccgg
aggtggaggagcaggatgatggcacgctgcccatgtcttgtgcccaggagagcgggatggaccgtcaccctgcagcctgtgctt
ctgctaggatcaatgtgtaggatccttgacttgcggccgcaactcccacctgcaacatgcgtgactgactgaggccgcgactctag
agtcgacctgcatctaggcgccggaattagatctctcgaggttaacgaattctaccgggtaggggaggcgcttttcccaaggcagt
ctggagcatgcgctttagcagccccgctgggcacttggcgctacacaagtggcctctggcctcgcacacattccacatccaccgg
taggcgccaaccggctccgttctttggtggccccttcgcgccaccttctactcctcccctagtcaggaagttcccccccgccccgca
gctcgcgtcgtgcaggacgtgacaaatggaagtagcacgtctcactagtctcgtgcagatggacagcaccgctgagcaatgga
agcgggtaggcctttggggcagcggccaatagcagctttgctccttcgctttctgggctcagaggctgggaaggggtgggtccgg
gggcgggctcaggggcgggctcaggggcggggcgggcgcccgaaggtcctccggaggcccggcattctgcacgcttcaaaa
gcgcacgtctgccgcgctgttctcctcttcctcatctccgggcctttcgacctgcagcccaagcttaccatggctgaaggatccgtcg
ccaggcagcctgacctcttgacctgcgacgatgagccgatccatatccccggtgccatccaaccgcatggactgctgctcgccct
cgccgccgacatgacgatcgttgccggcagcgacaaccttcccgaactcaccggactggcgatcggcgccctgatcggccgct
ctgcggccgatgtcttcgactcggagacgcacaaccgtctgacgatcgccttggccgagcccggggcggccgtcggagcaccg
atcactgtcggcttcacgatgcgaaaggacgcaggcttcatcggctcctggcatcgccatgatcagctcatcttcctcgagctcga
gcctccccagcgggacgtcgccgagccgcaggcgttcttccgccgcaccaacagcgccatccgccgcctgcaggccgccga
aaccttggaaagcgcctgcgccgccgcggcgcaagaggtgcggaagattaccggcttcgatcgggtgatgatctatcgcttcgc
ctccgacttcagcggcgaagtgatcgcagaggatcggtgcgccgaggtcgagtcaaaactaggcctgcactatcctgcctcaac
cgtgccggcgcaggcccgtcggctctataccatcaacccggtacggatcattcccgatatcaattatcggccggtgccggtcacc
ccagacctcaatccggtcaccgggcggccgattgatcttagcttcgccatcctgcgcagcgtctcgcccgtccatctggaattcatg
cgcaacataggcatgcacggcacgatgtcgatctcgattttgcgcggcgagcgactgtggggattgatcgtttgccatcaccgaa
cgccgtactacgtcgatctcgatggccgccaagcctgcgagctagtcgcccaggttctggcctggcagatcggcgtgatggaag
agtaa

pAG1819 CMV_NCRE-$^{\text{FIRE}}$mate_NLS_P2A_NLS_$^{\text{FIRE}}$tag-CCRE

atggccacctctgatgaagtcaggaagaacctgatggacatgttcagggacaggcaggccttctctgaacacacctggaagatg
ctcctgtctgtgtgcagatcctgggctgcctggtgcaagctgaacggtaccATGGAGCATGTCGCTTTCGGATCA
GAAGATATTGAAAACACCCTGGCGAATATGGACGATGAGCAACTTGATCGGTTGGCTTT
CGGGGTTATCCAGTTGGACGGTGATGGCAACATCCTGTTGTACAACGCTGCTGAGGGG
GATATTACAGGACGAGACCCCAAACAAGTGATAGGTAAAAATTTCTTTAAAGACGTTGCT
CCCGGAACAGATACGCCGGAATTCTATGGTAAGTTCAAGGAAGGGGCAGCTTCCGGCA
ATTTGAACACCATGTTCGAGTGGACAATTCCTACCAGTCGCGGTCCAACAAAAGTTAAA
GTGCACCTTAAAAAAGCACTCTCTggcggaagcggtggcgtgcccaagaagaagaggaaagtcggatccgg
cagcggcgccaccaacttcagcctgctgaagcaggccggcgacgtggaggagaaccccggccccctcgaggtgcccaaga
agaagaggaaagtcggcggaGGAGACAGATATTGGGTATTTGTTAAGCGCGTTggtaccaacaggaa
atggttccctgctgaacctgaggatgtgagggactacctcctgtacctgcaagccagaggcctggctgtgaagaccatccaacag
cacctgggccagctcaacatgctgcacaggagatctggcctgcctcgcccttctgactccaatgctgtgtccctggtgatgaggag
aatcagaaaggagaatgtggatgctggggagagagccaagcaggccctggcctttgaacgcactgactttgaccaagtcagat
ccctgatggagaactctgacagatgccaggacatcaggaacctggccttcctgggcattgcctacaacaccctgctgcgcattgc
cgaaattgccagaatcagagtgaaggacatctcccgcaccgatggtgggagaatgctgatccacattggcaggaccaagacc
ctggtgtccacagctggtgtggagaaggccctgtccctggggttaccaagctggtggagagatggatctctgtgtctggtgtggct
gatgaccccaacaactacctgttctgccgggtcagaaagaatggtgtggctgccccttctgccacctcccaactgtccacccggg
ccctggaagggatctttgaggccacccaccgcctgatctatggtgccaaggatgactctgggcagagatacctggcctggtctgg
ccactctgccagagtgggtgctgccagggacatggccagggctggtgtgtccatccctgaaatcatgcaggctggtggctggacc
aatgtgaacattgtgatgaactacatcagaaacctggactctgagactggggccatggtgaggctgctcgaagatggggac

pAG1867 CMV_p65Δ(286-550)-$^{\text{FIRE}}$mate

ATGcagtacctgccagatacagacgatcgtcaccggattgaggagaaacgtaaaaggacatatgagaccttcaagagcatc
atgaagaagagtcctttcagcggacccaccgacccccggcctccacctcgacgcattgctgtgccttcccgcagctcagcttctgt
ccccaagccagcaccccagccctatccctttacgtcatccctgagcaccatcaactatgatgagtttcccaccatggtgtttccttctg
ggcagatcagccaggcctcggccttggccccggcccctccccaagtcctgccccaggctccagcccctgcccctgctccagcc
atggtatcagctctggcccaggccccagcccctgtcccagtcctagccccaggccctcctcaggctgtggccccacctgcccccа
agcccacccaggctggggaaggaacgctgtcagaggccctgctgcagctgcagtttgatgatgaagacctgggggccttgcttg
gcaacagcacagacccagctgtgttcacagacctggcatccgtcgacaactccgagtttcagcagctgctgaaccagggcata
cctgtggcccccacacaactgagcccatgctgatggagtaccctgaggctataactcgcctagtgacaggggcccagaggcc
ccccgacccagctcctgctccactgggggccccggggctccccaatggcctcctttcaggagatgaagacttctcctccattgcgg
acatggacttctcagccctgctgagtcagatcagctccTCTCCAGCGATTCCGTCGACACCACCTACTCCC
TCTCCAGCGATCGCCATGGAAATGGAGCATGTCGCATTCGGGTCAGAAGATATCGAAAA
CACATTGGCTAATATGGATGACGAACAGCTGGATCGGCTCGCTTTCGGTGTCATACAAT
TGGACGGGGATGGCAACATTTTGTTGTATAATGCAGCAGAAGGTGATATAACTGGACGG
GACCCGAAGCAAGTGATCGGCAAGAACTTCTTTAAGGATGTCGCCCCCGGTACCGACA
CGCCTGAGTTCTATGGAAAATTCAAGGAGGGAGCTGCCAGTGGGAATTTGAACACGAT
GTTTGAGTGGACAATCCCGACGTCAAGGGGTCCGACCAAAGTCAAAGTGCATCTCAAG
AAGGCACTGTCTTAA

pAG1868 CMV_ GAL4Δ(1-65)-FIREtag

ATGAAGCTACTGTCTTCTATCGAACAAGCATGCGATATTTGCCGACTTAAAAAGCTCAAG
TGCTCCAAAGAAAAACCGAAGTGCGCCAAGTGTCTGAAGAACAACTGGGAGTGTCGCT
ACTCTCCCAAAACCAAAAGGTCTCCGCTGACTAGGGCACATCTGACAGAAGTGGAATCA
AGGCTAGAAAGACTGGAAGCGATTCCGTCGACACCACCTACTCCCTCTCCAGCGATCG
CCATGGAAGGAGACAGATATTGGGTATTTGTTAAGCGCGTT

pAG1869 CMV_NLS_ p65Δ (286-550)-FIREmate

ATGcccaagaagaagaggaaagtccagtacctgccagatacagacgatcgtcaccggattgaggagaaacgtaaaagga
catatgagaccttcaagagcatcatgaagaagagtcctttcagcggacccaccgacccccggcctccacctcgacgcattgctgt
gccttcccgcagctcagcttctgtccccaagccagcaccccagccctatccctttacgtcatccctgagcaccatcaactatgatga
gtttcccaccatggtgtttccttctgggcagatcagccaggcctcggccttggccccggcccctccccaagtcctgccccaggctcc
agcccctgcccctgctccagccatggtatcagctctggcccaggccccagcccctgtcccagtcctagccccaggccctcctcag
gctgtggccccacctgcccccaagcccacccaggctggggaaggaacgctgtcagaggccctgctgcagctgcagtttgatgat
gaagacctgggggccttgcttggcaacagcacagacccagctgtgttcacagacctggcatccgtcgacaactccgagtttcag
cagctgctgaaccagggcatacctgtggcccccacacaactgagcccatgctgatggagtaccctgaggctataactcgccta
gtgacaggggcccagaggcccccgacccagctcctgctccactgggggccccggggctccccaatggcctcctttcaggaga
tgaagacttctcctccattgcggacatggacttctcagccctgctgagtcagatcagctccTCTCCAGCGATTCCGTCG
ACACCACCTACTCCCTCTCCAGCGATCGCCATGGAAATGGAGCATGTCGCATTCGGGT
CAGAAGATATCGAAAACACATTGGCTAATATGGATGACGAACAGCTGGATCGGCTCGCT
TTCGGTGTCATACAATTGGACGGGGATGGCAACATTTTGTTGTATAATGCAGCAGAAGG
TGATATAACTGGACGGGACCCGAAGCAAGTGATCGGCAAGAACTTCTTTAAGGATGTCG
CCCCCGGTACCGACACGCCTGAGTTCTATGGAAAATTCAAGGAGGGAGCTGCCAGTGG

GAATTTGAACACGATGTTTGAGTGGACAATCCCGACGTCAAGGGGTCCGACCAAAGTCA
AAGTGCATCTCAAGAAGGCACTGTCTTAA

pAG1870 pGL4.31EmGFP-P2A-iCasp9-HA-hPEST

atggtgagcaagggcgaggagctgttcaccggggtggtgcccatcctggtcgagctggacggcgacgtaaacggccacaagtt
cagcgtgtctggcgagggcgagggcgatgccacctacggcaagctgaccctgaagttcatctgcaccaccggcaagctgccc
gtgccctggcccaccctcgtgaccaccctgacctacggcgtgcagtgcttcagccgctaccccgaccacatgaagcagcacga
cttcttcaagtccgccatgcccgaaggctacgtccaggagcgcaccatcttcttcaaggacgacggcaactacaagacccgcgc
cgaggtgaagttcgagggcgacaccctggtgaaccgcatcgagctgaagggcatcgacttcaaggaggacggcaacatcctg
gggcacaagctggagtacaactacaacagccacaacgtctatatcatggccgacaagcagaagaacggcatcaaggcgaa
cttcaagatccgccacaacatcgaggacggcagcgtgcagctcgccgaccactaccagcagaacacccccatcggcgacgg
ccccgtgctgctgcccgacaaccactacctgagcacccagtccgccctgagcaaagaccccaacgagaagcgcgatcacatg
gtcctgctggagttcgtgaccgccgccgggatcactctcggcatggacgagctgtacaagttaatcgccggaagcggagctacta
acttcagcctgctgaagcaggctggagacgtggaggagaaccctggacctaccatgctcgagtctggcggtggatccggagtc
gacggatttggtgatgtcggtgctcttgagagtttgaggggaaatgcagatttggcttacatcctgagcatggagccctgtggccact
gcctcattatcaacaatgtgaacttctgccgtgagtccgggctccgcacccgcactggctccaacatcgactgtgagaagttgcgg
cgtcgcttctcctcgctgcatttcatggtggaggtgaagggcgacctgactgccaagaaaatggtgctggctttgctggagctggcg
cggcaggaccacggtgctctggactgctgcgtggtggtcattctctctcacggctgtcaggccagccacctgcagttcccaggggc
tgtctacggcacagatggatgccctgtgtcggtcgagaagattgtgaacatcttcaatgggaccagctgccccagcctgggaggg
aagcccaagctctttttcatccaggcctgtggtggggagcagaaagaccatgggtttgaggtggcctccacttcccctgaagacg
agtcccctggcagtaaccccgagccagatgccaccccgttccaggaaggtttgaggaccttcgaccagctggacgccatatcta
gtttgcccacacccagtgacatctttgtgtcctactctactttcccaggttttgtttcctggagggaccccaagagtggctcctggtacgt
tgagaccctggacgacatctttgagcagtgggctcactctgaagacctgcagtccctcctgcttagggtcgctaatgctgtttcggtg
aaagggatttataaacagatgcctggttgctttaatttcctccggaaaaaacttttctttaaaacatcagtcgactgtaaatattgtcatg
aactatatccgtaacctggatagtgaaacaggggcaatggtgcgcctgctggaagatggcgattagtatccgtacgacgtacca
gactacgcaAATTCTCACGGCTTCCCTCCCGAGGTGGAGGAGCAGGCCGCCGGCACCCTG
CCCATGAGCTGCGCCCAGGAGAGCGGCATGGATAGACACCCTGCTGCTTGCGCCAGC
GCCAGGATCAACGTCTAA

pAG1930 pGL4.31[EmGFP-P2A-TEV-HA-hPEST]

atggtgagcaagggcgaggagctgttcaccggggtggtgcccatcctggtcgagctggacggcgacgtaaacggccacaagtt
cagcgtgtctggcgagggcgagggcgatgccacctacggcaagctgaccctgaagttcatctgcaccaccggcaagctgccc
gtgccctggcccaccctcgtgaccaccctgacctacggcgtgcagtgcttcagccgctaccccgaccacatgaagcagcacga
cttcttcaagtccgccatgcccgaaggctacgtccaggagcgcaccatcttcttcaaggacgacggcaactacaagacccgcgc
cgaggtgaagttcgagggcgacaccctggtgaaccgcatcgagctgaagggcatcgacttcaaggaggacggcaacatcctg
gggcacaagctggagtacaactacaacagccacaacgtctatatcatggccgacaagcagaagaacggcatcaaggcgaa
cttcaagatccgccacaacatcgaggacggcagcgtgcagctcgccgaccactaccagcagaacacccccatcggcgacgg
ccccgtgctgctgcccgacaaccactacctgagcacccagtccgccctgagcaaagaccccaacgagaagcgcgatcacatg
gtcctgctggagttcgtgaccgccgccgggatcactctcggcatggacgagctgtacaagttaatcgccggaagcggagctacta
acttcagcctgctgaagcaggctggagacgtggaggagaaccctggacctaccatgctcgagatgggcgagagccttttcaag
ggcccgagggactacaacccgatctccagcaccatctgtcacctgaccaacgagagcgacggtcacaccactagtctgtacgg
catcggcttcggccccttcatcatcaccaacaagcatctgttcaggaggaataacggcacactgctggtgcaaagcctgcacgg
cgtgttcaaagtgaagaacacaaccaccctgcaacagcacctgatcgacggcagggacatgattatcatcaggatgcccaag

gacttcccccccttttccccagaaactgaagttcagggagccacaaagggaggagcgaatctgcctggtgaccaccaacttcca
gaccaagtccatgagcagcatggtctctgataccagctgcaccttccccagcagcgacggcatcttctggaagcactggattcag
acgaaggatggccaatgcggcagcccattggtgagcactagggacggcttcatcgtgggcatccacagcgccagcaattttacc
aataccaacaactacttcacgagcgtgccgaaaaacttcatggagctgttgaccaatcaagaggcgcagcagtgggtgagcgg
ctggaggctgaacgccgacagcgttctttggggcggacataaggtgttcatggtcaagcccgaggaacccttccagcccgttaa
ggaagccactcagctttatccgtacgacgtaccagactacgcaAATTCTCACGGCTTCCCTCCCGAGGTGGA
GGAGCAGGCCGCCGGCACCCTGCCCATGAGCTGCGCCCAGGAGAGCGGCATGGATAG
ACACCCTGCTGCTTGCGCCAGCGCCAGGATCAACGTCTAA

pAG1931 pGL4.31[EmGFP-P2A-CRE-HA-hPEST]

atggtgagcaagggcgaggagctgttcaccggggtggtgcccatcctggtcgagctggacggcgacgtaaacggccacaagtt
cagcgtgtctggcgagggcgagggcgatgccacctacggcaagctgaccctgaagttcatctgcaccaccggcaagctgccc
gtgccctggcccaccctcgtgaccaccctgacctacggcgtgcagtgcttcagccgctaccccgaccacatgaagcagcacga
cttcttcaagtccgccatgcccgaaggctacgtccaggagcgcaccatcttcttcaaggacgacggcaactacaagacccgcgc
cgaggtgaagttcgagggcgacaccctggtgaaccgcatcgagctgaagggcatcgacttcaaggaggacggcaacatcctg
gggcacaagctggagtacaactacaacagccacaacgtctatatcatggccgacaagcagaagaacggcatcaaggcgaa
cttcaagatccgccacaacatcgaggacggcagcgtgcagctcgccgaccactaccagcagaacacccccatcggcgacgg
ccccgtgctgctgcccgacaaccactacctgagcacccagtccgccctgagcaaagaccccaacgagaagcgcgatcacatg
gtcctgctggagttcgtgaccgccgccgggatcactctcggcatggacgagctgtacaagttaatcgccggaagcggagctacta
acttcagcctgctgaagcaggctggagacgtggaggagaaccctggacctaccatgctcgagatgtccaatttactgaccgtac
accaaaatttgcctgcattaccggtcgatgcaacgagtgatgaggttcgcaagaacctgatggacatgttcagggatcgccaggc
gttttctgagcatacctggaaaatgcttctgtccgtttgccggtcgtgggcggcatggtgcaagttgaataaccggaaatggtttcccg
cagaacctgaagatgttcgcgattatcttctatatcttcaggcgcgcggtctggcagtaaaaactatccagcaacatttgggccagc
taaacatgcttcatcgtcggtccgggctgccacgaccaagtgacagcaatgctgtttcactggttatgcggcggatccgaaaaga
aaacgttgatgccggtgaacgtgcaaaacaggctctagcgttcgaacgcactgatttcgaccaggttcgttcactcatggaaaata
gcgatcgctgccaggatatacgtaatctggcatttctggggattgcttataacaccctgttacgtatagccgaaattgccaggatcag
ggttaaagatatctcacgtactgacggtgggagaatgttaatccatattggcagaacgaaaacgctggttagcaccgcaggtgta
gagaaggcacttagcctgggggtaactaaactggtcgagcgatggatttccgtctctggtgtagctgatgatccgaataactacct
gttttgccgggtcagaaaaaatggtgttgccgcgccatctgccaccagccagctatcaactcgcgccctggaagggatttttgaag
caactcatcgattgatttacggcgctaaggatgactctggtcagagatacctggcctggtctggacacagtgcccgtgtcggagcc
gcgcgagatatggcccgcgctggagtttcaataccggagatcatgcaagctggtggctggaccaatgtaaatattgtcatgaact
atatccgtaacctggatagtgaaacagggggcaatggtgcgcctgctggaagatggcgattatccgtacgacgtaccagactacg
caAATTCTCACGGCTTCCCTCCCGAGGTGGAGGAGCAGGCCGCCGGCACCCTGCCCAT
GAGCTGCGCCCAGGAGAGCGGCATGGATAGACACCCTGCTGCTTGCGCCAGCGCCAG
GATCAACGTCTAA

pAG2013 UAS_PeptideB-FCS-NanoLuc-FCS-PeptideA- FCS TMD

atggccctgtggatgcgcttcctgcccctgctggccctgctcttcctctgggagtcccaccccacccaggcttttgtcaagcagcacc
tttgtggttcccacctggtggaggctctctacctggtgtgtggggagcgtggcttcttctacacacccatgtccCGTACAaagagg
gtcttcacactcgaagatttcgttggggactggcgacagacagccggctacaacctggaccaagtccttgaacagggaggtgtgt
ccagtttgtttcagaatctcggggtgtccgtaactccgatccaaaggattgtcctgagcggtgaaaatgggctgaagatcgacatcc

atgtcatcatcccgtatgaaggtctgagcggcgaccaaatgggccagatcgaaaaaatttttaaggtggtgtaccctgtggatgat
catcactttaaggtgatcctgcactatggcacactggtaatcgacggggttacgccgaacatgatcgactatttcggacggccgtat
gaaggcatcgccgtgttcgacggcaaaaagatcactgtaacagggaccctgtggaacggcaacaaaattatcgacgagcgcc
tgatcaaccccgacggctccctgctgttccgagtaaccatcaacggagtgaccggctggcggctgtgcgaacgcattctggcgC
GTCAAaagcgtggcattgtagatcagtgctgcaccagcatctgctccctctaccagctggagaactactgcaacgccgcgcac
ccgggtggatcctctggcagaagcaaaagaagcgtcattgacggctttaccctgaccagcgacgaggtgtgggtggtgggcatg
ggcatcgtcatgtctctcatcgtcctggccatcgtgtttggcaatgtgctggtcatcacagccattgccaagttcgagcgtctgcagac
ggtcaccgggggcggctctctcgagggggggggctcggggctcgaaaacttgtacttccagagtggcagtggaggtaccaatta
cttcatcacttcactggcctgtgctgatctggtcatgggcctggcagtggtgccctttggggccgcccatattcttatgaaaatgtggac
ttttggcaacttctgggatatctacatctgggctcccttggccgggacttgtggggtccttctcctgtcactggttatcaccctttactgca
acaagtacaaaagcagacgcagttttattgatgaagccgccatgccttag

pAG2014 CMV_PeptideB-FCS-NanoLuc-FCS-PeptideA-TMD

atggccctgtggatgcgcttcctgcccctgctggccctgctcttcctctgggagtcccaccccacccaggcttttgtcaagcagcacc
tttgtggttcccacctggtggaggctctctacctggtgtgtggggagcgtggcttcttctacacacccatgtccCGTACAaagagg
gtcttcacactcgaagatttcgttggggactggcgacagacagccggctacaacctggaccaagtccttgaacagggaggtgtgt
ccagtttgtttcagaatctcggggtgtccgtaactccgatccaaaggattgtcctgagcggtgaaaatgggctgaagatcgacatcc
atgtcatcatcccgtatgaaggtctgagcggcgaccaaatgggccagatcgaaaaaatttttaaggtggtgtaccctgtggatgat
catcactttaaggtgatcctgcactatggcacactggtaatcgacggggttacgccgaacatgatcgactatttcggacggccgtat
gaaggcatcgccgtgttcgacggcaaaaagatcactgtaacagggaccctgtggaacggcaacaaaattatcgacgagcgcc
tgatcaaccccgacggctccctgctgttccgagtaaccatcaacggagtgaccggctggcggctgtgcgaacgcattctggcgC
GTCAAaagcgtggcattgtagatcagtgctgcaccagcatctgctccctctaccagctggagaactactgcaacgccgcgcac
ccgggtggatcctctggcagaagcaaaagaagcgtcattgacggctttaccctgaccagcgacgaggtgtgggtggtgggcatg
ggcatcgtcatgtctctcatcgtcctggccatcgtgtttggcaatgtgctggtcatcacagccattgccaagttcgagcgtctgcagac
ggtcaccgggggcggctctctcgagggggggggctcggggctcgaaaacttgtacttccagagtggcagtggaggtaccaatta
cttcatcacttcactggcctgtgctgatctggtcatgggcctggcagtggtgccctttggggccgcccatattcttatgaaaatgtggac
ttttggcaacttctgggatatctacatctgggctcccttggccgggacttgtggggtccttctcctgtcactggttatcaccctttactgca
acaagtacaaaagcagacgcagttttattgatgaagccgccatgccttag

### References


(1) Bottone, S.; Joliot, O.; Cakil, Z. V.; El Hajji, L.; Rakotoarison, L.-M.; Boncompain, G.; Perez, F.; Gautier, A. A Fluorogenic Chemically Induced Dimerization Technology for Controlling, Imaging and Sensing Protein Proximity. *Nature Methods* **2023**, *20* (10), 1553–1562. https://doi.org/10.1038/s41592-023-01988-8.

(2) Rakotoarison, L. M.; Tebo, A. G.; Böken, D.; Board, S.; El Hajji, L.; Gautier, A. Improving Split Reporters of Protein-Protein Interactions through Orthology-Based Protein Engineering. *ACS Chemical Biology* **2024**, *19* (2), 428–441. https://doi.org/10.1021/acschembio.3c00631.

(3) El Hajji, L.; Lam, F.; Avtodeeva, M.; Benaissa, H.; Rampon, C.; Volovitch, M.; Vriz, S.; Gautier, A. Multiplexed In Vivo Imaging with Fluorescence Lifetime-Modulating Tags. *Advanced Science* **2024**, *11* (32). https://doi.org/10.1002/advs.202404354.
